# Genomic insights into the differentiated population admixture structure and demographic history of North East Asians

**DOI:** 10.1101/2021.07.19.452943

**Authors:** Guanglin He, Mengge Wang, Xing Zou, Renkuan Tang, Hui-Yuan Yeh, Zheng Wang, Xiaomin Yang, Ziyang Xia, Yingxiang Li, Jianxin Guo, Rui Wang, Jing Liu, Kongyang Zhu, Jing Chen, Meiqing Yang, Qu Shen, Jinwen Chen, Jing Zhao, Hao Ma, Lan-Hai Wei, Ling Chen, Changhui Liu, Chao Liu, Gang Chen, Yiping Hou, Chuan-Chao Wang

**Author notes:** Corresponding author (**G.L. H.**), (**C.L.**), (**G.C.**), (**Y.P. H.**) and (**C.C. W.**). These authors contributed equally to this work and should be considered co-first authors (Guanglin He, Mengge Wang).

## Abstract

North China and South Siberia, mainly populated by Altaic-speaking populations, possess extensive ethnolinguistic diversity and serve as the crossroad for the initial peopling of America and western-eastern trans-continental communication. Yet, the complex scenarios of genetic origin, population structure, and admixture history of North-East Asia remain to be fully characterized, especially for Mongolic people in China with a genome-wide perspective. Thus, we genotyped genome-wide SNPs for 510 individuals from 38 Chinese Mongolic, Tungusic, and Sinitic populations to explore the sharing alleles and haplotypes within the studied groups and following merged it with 3508 modern and ancient Eurasian individuals to reconstruct the deep evolutionary and natural selection history of northern East Asians. We identified significant substructures within Altaic-speaking populations with the primary common ancestry linked to the Neolithic northern East Asians: Western Turkic people harbored more western Eurasian ancestry; Northern Mongolic people in Siberia and eastern Tungusic people in Amur River Basin (ARB) possessed dominant Neolithic Mongolian Plateau (MP) or ARB ancestry; Southern Mongolic people in China owned obvious genetic impact from Neolithic Yellow River Basin (YRB) farmers. Additionally, we found the differentiated admixture history between western and eastern Mongolians and geographically close Northeast Hans: the former received a genetic impact from western Eurasians and the latter retained the dominant YRB and ARB Neolithic ancestry. Moreover, we demonstrated that Kalmyk people from the northern Caucasus Mountain possessed a strong genetic affinity with Neolithic MP people, supporting the hypothesis of their eastern Eurasian origin and long-distance migration history. We also illuminated that historic pastoral empires in the MP contributed considerably to the gene pool of northern Mongolic people but rarely to southern ones. We finally found natural signatures in Mongolians associated with alcohol metabolism. Generally, our results not only illuminated that complex population migration and admixture of Neolithic ancestral sources from the MP or ARB played an important role in the spread of Altaic-speaking populations and Proto-Altaic language, which partly supported the Northeast Asia-origin hypothesis, but also demonstrated that the observed multi-sources of genetic diversity contributed significantly to the modern existing extensive ethnolinguistic diversity in North-East Asia.

## INTRODUCTION

North China and South Siberia have been inhabited by anatomically modern humans as early as 40 thousand years ago (kya)(1). Two hypotheses, referred to as the southern migration hypothesis(2) and northern migration hypothesis(3), have been put forward to elucidate the mechanism and process of peopling of East Asia and Siberia. The southern migration hypothesis states that the ancestor of East Asians diverged with the ancestor of Europeans at approximately 49∼54 kya in South Asia or the Arabian Peninsula and then split with the ancestor of Oceanians in Southeast Asia at approximately 44∼49 kya and finally formed ancient East Asians via this southern route(4). The northern migration hypothesis states that one wave of ancestral population diverged from the ancestor of Europeans, Australians, and East Asians and migrated via Central Asia into North Siberia, forming Ancient North Eurasian and Ancient North Siberian, and finally mixed with incoming ancestors from East Asia to form the founding populations of Americans(5, 6). Crossroads of Mongolian Plateau (**MP**) and surrounding regions between South East Asia and North Siberia played a key role in the formation of ancient and modern Eastern Eurasians. Indeed, archeological and genetic evidence from the Chinese Tianyuan Cave (40,000-year-old Tianyuan)(1) and Siberian Paleolithic ancient people (31,600-year-old Yana_RHS, 17,000-year-old AG2, 16,500-year-old AG3, and 24,000-year-old Mal’ta)(3, 7, 8) have suggested that at least two Paleolithic eastern Eurasian lineages widely existed in East Asia and Siberia, and these two lineages have participated in the formation of Northeast Eurasians and Americans in the crossroad of Baikal Lake and Amur River Basin (**ARB**) regions.

Neolithic Trans-Eurasian migration and subsequent admixture or transformation played a pivotal role in the formation of ancestral Eurasians (especially in Siberia) and also contributed to the observed modern mosaic genetic structure(9-12). Similarly, Neolithic genetic evidence in East Asia not only documented two deeply diverged eastern Eurasian lineages(13), which composed of Hoabinhian-related clade in Southeast Asia and Jomon-related lineage in the Japanese archipelago, but also identified at least five Neolithic East Asian lineages (inland/coastal southern/northern East Asian lineages respectively associated with Liangdao/Qihe in Fujian, Longlin in Guangxi, Bianbian and Yumin in **Yellow River Basin (YRB)**, and Chokhopani in Tibet, as well as the ancient northeast East Asian lineages related to Neolithic DevilsGate and Boisman in **MP** and **ARB**)(14-17). These observed patterns of the admixed ancestry among northern and southern Neolithic East Asians showed that both southward and northward gene flow events and coastal population connections from Vietnam and coastal Siberia have reshaped the subsequent genetic landscape of East Asia. Further intense interplays between the coastal and inland East Asian pre-Neolithic Hunter-Gatherers and Neolithic farmers in Fujian and Guangxi, as well as lower or middle or upper YRB, were illuminated in the perspectives of both archeology and ancient genomes. Besides, Neolithic ancient DNA evidence from the Baikal Lake region(7, 18) and ARB(7, 19) not only suggested the different extent of genetic continuity that existed in eastern Siberia but also demonstrated extensive genetic interaction and connection between YRB farmers and other Siberian ancient Hunter-Gatherers(20).

Differentiated western Eurasian ancestral sources contributed different genetic components into the gene pool of North-East Asia. In the Copper or early and middle Bronze Age, the archeologically documented material culture showed that Yamnaya pastoralists, formed in the Pontic-Caspian steppe (∼3000 BCE), revolutionized the perception of personhood, property, and family(10, 11). These early herders shaped the genetic structure of later populations in Europe and South Asia(21, 22). Subsequent cultural shifts, such as Afanasievo culture in the Minusinsk Basin and the Altaic Mountain, Sintashta culture emerged in the Urals, Andronovo culture in Central Asia and even in Xinjiang and Sayan Mountain, were significantly associated with the demic diffusion of early eastward migration of steppe pastoralists(10, 11, 22). European or Anatolian farmers following admixed with local Pontic-Caspian herders, which then formed the Late Bronze Age pastoralists with different cultural backgrounds (Koryakova, Mezhovskaya, Srubnaya, Okunevo, and Karasuk)(9, 12). Although multiple genetic turnovers and admixture events occurred in western Eurasia, the Bronze Age population history of eastern Eurasians kept a genetic continuity as the Neo-Siberians, which has been evidenced via sporadic studies of ancient DNA(7, 12). Subsistence strategies of these people kept primary hunting-gathering/fishing or reindeer different from the pastoral lifestyles in the west Eurasia. Recent 6000-year genomic dynamics in Mongolia showed the instantaneous admixture signatures between western and eastern Eurasians in central Mongolia, as the observed tripartite population structures starting to vanish(9).

More dynamic population history occurred in the eastern steppe during the Iron and historic periods. Population expansion directions have gradually changed from the previously dominant eastward expansion to radial expansion from multiple dominant centers (such as the Scythian federation) and finally to westward dispersal via historic pastoral tribes(11, 23). Historic empires of nomadic pastoralists combined and formed many elite dominance federation groups, such as Turkic, Xiongnu, Mongolia, and Tungusic et al., and the population expansion centers have been shifted from West Eurasia to East Eurasia(9, 20). And the basic patterns of the genetic background of modern populations were gradually formed, with the eastern Eurasian-related genetic diversity replaced or mixed with the Proto-Indo-European’s gene pool as well as their languages(7, 10). Although the landscapes of three genetic clines in modern Siberians have been characterized, how their genetic relationship with Chinese Altaic populations and the potentially existing differentiated evolutionary population history needed to be comprehensively clarified and characterized.

The language/farming co-dispersal hypothesis has been evidenced in the spread of Sino-Tibetan, and micro-family in South China and Southeast Asia(20), as well as the possible association between the spread of Anatolian farmer ancestry and Indo-European people(12). Linguistic and archaeologic evidence also consistently supported that early dissemination of Indo-European languages was mediated by the steppe pastoralists’ expansion(21, 22). Recent findings have revealed the influence of the co-spread of this language and the corresponding ancient Indo-European people extending to 2200-year-old Iron Age Shirenzigou people in the northeastern Xinjiang(24). Patterns of genetic structure and Indo-European distribution were also influenced and characterized by large-scale eastward expansion and replacements of local populations(7). Genomic evidence has found that these population migrations have no significant influence on the population demographic dynamics in Baikal Lake and ARB regions and North China(25). After the Bronze Age migrations, population interaction between western and eastern Eurasian steppe has emerged among highly structured Scythian groups with possible Turkic language, whose gene pool was consisted of western sources (European farmer and Late Bronze Age herder), eastern source of southern Siberian Hunter-Gatherer and Anatolian or Iran farmer-related ancestry(26). Further westward dispersal of Xiongnu and Hun Khanates, and other historical migration events (expansion of Mongolian Empire) have promoted Trans-Eurasian-speaking populations (Turkic, Mongolic and Tungusic) dominated in the Eurasian steppe and replaced previous Indo-Iranian-speaking groups (Wusun, Kangju et al.)(26). However, the detailed interaction between YRB farmers and Siberian Hunter-Gatherers, and what the extent of genetic contribution from these ancient populations to modern northern East Asians, as well as who mediated the dispersal of modern Altaic-speaking populations needed to be further explored. The geographical origins of Altaic-speaking populations have been stated to be controversial based on evidence from genetics and linguistics, including Pastoralist Hypothesis or Farming Hypothesis supported from the Altaic Mountain, West Liao River, Baikal Lake and ARB(20). These controversial hypotheses needed to be tested via dense sampling of modern Altaic people and comprehensive comparison with spatiotemporally ancient sources. As the demographic and evolutionary history of the eastern regions (Baikal, ARB, MP and YRB) is still in its infancy and unclear compared with the well-documented complex admixture history in the western Eurasian steppe. These regions have been inhabited mainly by Mongolic and Tungusic-speaking populations today and there are also some remnants of early Turkic-speaking populations and historic northward expanded Han Chinese.

To comprehensively reconstruct the deep population history, explore the ancestral origin and population dynamics of Tungusic/Mongolic-speaking populations and their neighbors, we genotyped 510 modern individuals from Mongolic, Tungusic, and Sinitic-speaking populations using a high-density SNP array and combined the basic dataset with publicly available worldwide modern and ancient genome-wide data(3, 4, 7, 9-16, 20-22, 26, 27). This work mainly aimed to explore **I**) how many ancestral sources contributed to modern Altaic-speaking populations, **II**) what’s the association between the identified ancestries and the origin and mixture of modern Altaic language macro-family, as well as **III**) what’s the detailed admixture and natural selection histories of subgroups of Altaic-speaking populations and geographically close Northeast Hans, and finally **IV**) illuminate the genetic contribution from the historic pastoral empires from the MP to the southernmost Yunnan and Guangzhou Mongolians and westernmost Kalmyk people. We reconstructed the deep population genetic history of North-East Asia using multiple computational statistical methods (including sharing-allele *f*-statistics, haplotype-based chromosome painting skills, and sharing paternal and maternal founding lineages). We found a shared Neolithic ancestry maximized in Neolithic populations from the MP and ARB, which was widely distributed among all northern Mongolic/Tungusic-speaking populations from as far east as coastal Siberia (Ulchi) to as far west as the northern Caucasus (Kalmyk). We also identified population substructures within the Altaic-speaking populations, suggesting that the geographically diverse sampling populations were the mixture results of different ancestral sources or proportions. We also identified biogeographically structured Mongolian populations: western Mongolian with substantial ancestry related to western Eurasians, and eastern Mongolian with excessive shared alleles associated with Neolithic southern East

Asians. We finally provided one case for a better understanding of deep population history via the shared alleles, haplotypes and paternal lineages.

## RESULTS

### General population structure

We successfully genotyped around 500K genome-wide SNPs from 510 individuals from Mongolian, Daur, Ewenki, Hezhen and Han in 40 populations (**Fig. 1A**). We merged our data with population data of modern and ancient East Asians included in the Human Origin dataset (110K). Eurasian PCA among 3508 individuals clustered studied populations with Eastern Eurasian North-to-South cline (**Supplementary Figs. 1∼3**). The final eastern Eurasian dataset included 2826 individuals from 291 populations. Patterns of population genetic relationship in eastern Eurasian PCA showed an obvious separation between northern and southern East Asians along PC1, in which the southern East Asians consisting of Austronesian, Austroasiatic, Tai-Kadai and Hmong-Mien people and the counterpart consisting of the Altaic-speaking populations (Mongolic, Tungusic and others, **Fig. 1B**). Hmong-Mien-speaking populations from southwestern China and Mainland Southeast Asia formed one separated clade, which was different from the Tai-Kadai, Austronesian and Austroasiatic people from South China and Southeast Asia (**Fig. 1C**). Studied Mongolic and Tungusic people and the previously published ones from North China showed a separated localization with other northern Mongolic and Tungusic speakers from southern Siberia, Outer Mongolia and the Amur River. Ancient projected populations (LateXiongnu_sarmatian, EarlyMed_Uigur and EarlyXiongnu_west) also separated from all included modern Tungusic and Mongolic people along PC2, but other ancient populations with high Neolithic northeastern ancestry overlapped with modern Tungusic and Mongolic people. This observed deviation from Bronze Age/historic Altaic Mountain populations suggested that the primary ancestry of the modern Tungusic and Mongolic people originated from the eastern ancestral source from the MP or ARB. As expected, our studied Mongolic and Tungusic populations not only separated with southern and central Han Chinese but also slightly separated with geographically close northeastern Hans. Neolithic people from YRB were clustered closely with Hans than with Mongolians. Focused on the genetic variations of modern populations from North China and southern Siberia, cluster patterns inferred from the PCA showed more interesting population substructures among them, consistent with the categories of the subbranches of Altaic languages. PC1 separated Sinitic speakers and Mongolic and Tungusic people, and PC3 separated Tungusic and Mongolic people (**Fig. 1D∼E**)

**Fig. 1.**
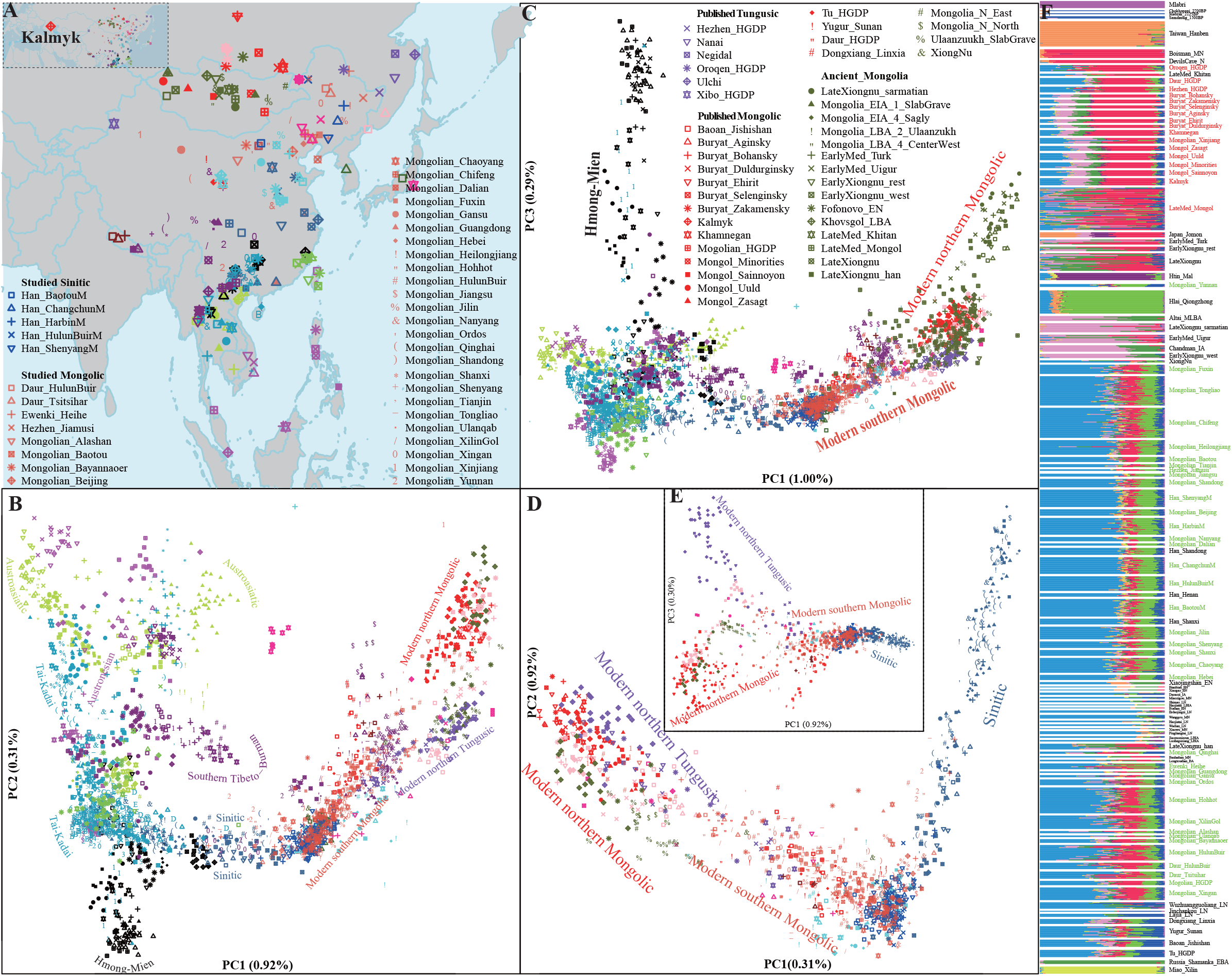
Geographic position and genetic structure of studied Hans, Mongolic and Tungusic people, as well as other modern and ancient East Asians based on the merged HO dataset (113,962 SNPs). (**A**) Map of the newly sampled 40 Chinese populations and other 251 modern and ancient reference populations. (**B∼C**) population genetic relationship patterns in the PCA analyses focused on all eastern Eurasian populations based on the top three components. Modern populations were colored-coded based on language categories (**D∼E**) Northern East Asian PCA analysis based on the top three components. Ancient samples were projected. (**F**) ADMIXTURE analysis result for K=10 which has the minimized cross-validation error.

Admixture results based on the modern and ancient Eurasians further showed the mixture nature of our studied populations and other Mongolic and Tungusic people. Different from the homogeneous genetic structure observed in some ancient or geographically/ethnically representative source populations, most of the modern East Asians were showed as the mosaic form as a mixture of at least two ancestry components. Red ancestry was maximized in Boisman_MN and DevilsCave_N Neolithic people, which was previously referred to as Neo-Siberian ancestry or northeastern Asian ancestry in ARB. Light-blue ancestry was maximized in Neolithic YRB millet farmers from Henan, Shandong and Gansu provinces (Wanggou_MN, Lajia_LN and Erdaojingzi_LN). Mongolic-speaking Buryat in Russia and Mongols in Mongolia harbored remarkable red and light-blue ancestries, which showed a strong genetic affinity with indigenous Siberian ancients. They also had pink ancestry related to western Eurasians and light-green ancestry related to Russia-Shamanka_EBA, providing clues of the effect of western Eurasians and indigenous Baikal ancients on the genetic structure of northern Mongolic populations. Different from northern Mongolic and Tungusic people, new studied Mongolians and Hans in China possessed much light-blue ancestry, and limited dark-blue, red and light-green ancestries. Light-green ancestry dominant in Qiongzhong Hlai was one of the representative ancestral sources of southern East Asians, and dark-blue ancestry related to Nepal ancients was the representative source of indigenous Tibetan ancients. Southern Altaic-speaking populations in China harbored more Neolithic YRB farmer-related ancestry and little Siberian-related ancestry (**Fig. 1F**). And more western Eurasian ancestry was visualized in Turkic people. Similar patterns of predefined ancestry sources and ancestry composition were also identified with different included reference populations in the model-based ADMIXTURE results (**Supplementary Figs. 4∼7**)

### The differentiated demographic history between western and eastern Mongolians

From the PCA results in **Fig. 1**, we observed the western Eurasian affinity of Mongolians from Xinjiang, Qinghai, Gansu and western Inner Mongolia. Ancestry composition inferred from ADMIXTURE modeling showed that Xinjiang Mongolians had 0.186 western Eurasian pink ancestry, 0.343 Neolithic Amur red ancestry, 0.283 Neolithic millet farmer light-blue ancestry and some of Neolithic Mongolia (0.070) and Tibeto-Burman (0.046) ancestry. Thus, we first explored the genetic structure of our newly genotyped 510 individuals and two previously published Hainan Hans and Hlai genotyped using the same Affymetrix array (**466K**). Plink-based PCA results showed the genetic differentiation between northern and southern East Asians (**Fig. 2A**). The number and length of the shared ancestry fragment were more powerful to dissect the finer-scale population structure. We obtained the shared haplotype via phasing and painting our generated genome-wide SNP data and then explored the coancestry matrix and clustering patterns. PCA results along the top four components extracted from the coancestry matrix showed the clear separation between Xinjiang Mongolians and other eastern Mongolians (**Fig. 2B∼D**). Heatmap of pairwise coincidence and clustering dendrogram based on individual-level and population-level shared chunk counts showed more genetically homogeneous subgroups in accordance (**Fig. 2E∼F**). Southernmost Hans formed the same branch with geographically close indigenous Hlai. New studied populations were separated into two major branches and more subbranches. One branch consisted of most of the Mongolic and Tungusic people and the other comprised of northern Hans and some Mongolians possessing more modern southern East Asian ancestry. We found geographically close populations shared longer IBD fragments, larger outgroup-*f*_*3*_ values and smaller Fst values (**Fig. 2G∼H**). Two southernmost Chinese Hainan populations had the shortest shared IBD fragments and the largest Fst values with our newly studied northern East Asians. Different from the southern indigenous Hlai (sharing IBD length within the population is 137.456), the identified shared IBD within geographically defined northern Mongolians was shorter, which suggested more recent admixture events occurred in the isolated Hainan populations consistent with the observed longer ROH in Hlai people. We also observed the Mongolians separated from Northeast Han in the clustering patterns, suggesting their differentiated demographic history. Modeling of ancestry composition based on the ADMIXTURE showed that the differentiated ancestry compositions were observed in Xinjiang and other eastern Mongolians (**Fig. 2I**). Pairwise Fst and inverse outgroup-*f*_*3*_-values among populations from the merged HO dataset also showed different genetic affinity between western and eastern Mongolians with their Eurasian references. Indeed, clustering patterns in the multidimensional scaling plots (MDS), N-J tree and heatmap based on these two-type genetic distances further confirmed that western Mongolians showed a close relationship with western Eurasians and Northern Altaic people, but eastern Mongolians were clustered closely with Sinitic people (**Supplementary Figs. 8∼22**).

**Fig. 2.**
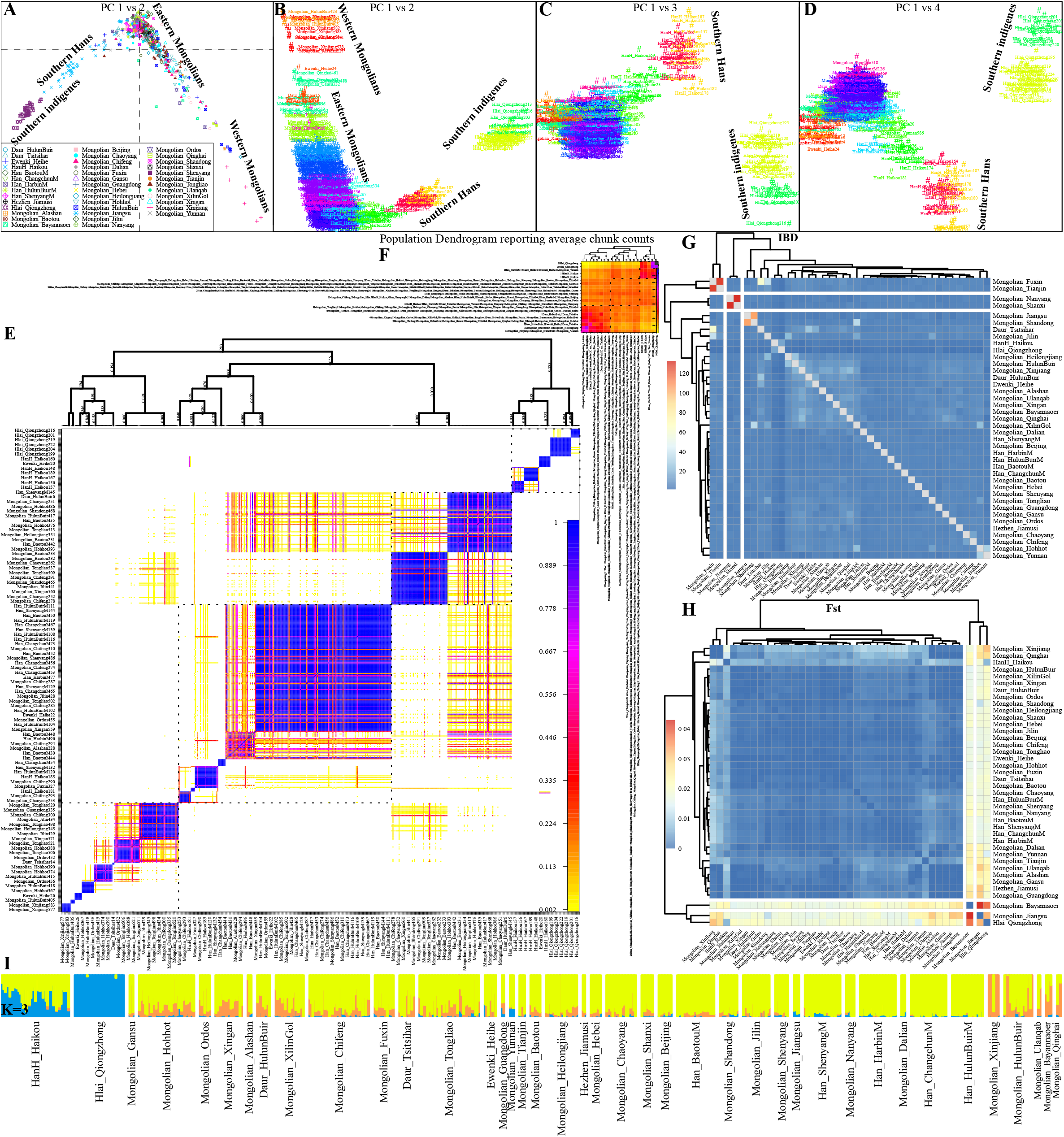
Fine-scale population genetic structure of studied populations and two Hainan populations based on 465,941SNPs genotyped using the Affymetrix chip. (A) Plink-based PCA results based on the allele frequency spectrum showed two clines within 573 individuals from 40 populations. (**B∼D**) PCA patterns based on the coancestry matrix of linked SNP markers. (**E∼F**) Pairwise coincidence matrix estimated based on the chunk counts using the linked SNPs and population-level average chunk counts. The coloring in E represented the posterior coincidence probability. (**G∼H**) Heatmap showed the pairwise sharing IBD length and pairwise Fst genetic distance within 40 populations.

We following assessed the detailed admixture landscape of Mongolians with more reference populations included in the merged 1240K dataset (**369K**). Results from the pairwise qpWave analysis focused on our western Mongolians showed multiple differentiated allele-sharing between the left and right populations, suggesting western Mongolians did not form a clade with eastern Mongolians and other reference populations relative to these used right populations (**Fig. 3A**). The different patterns of genetic affinity between western and eastern Mongolians were further confirmed via the different affinities observed in outgroup-*f*_*3*_ in the form *f*_*3*_(Eurasians, eastern/western Mongolians; Mbuti), diverse admixture signatures via the observed admixture signatures in the admixture-*f*_*3*_(Source1, Source2; western/eastern Mongolians). Rank results of p_rank1 (0.0004) and p_rank2 (0.9567) showed that at least two admixture events were needed to explain the admixture landscape of western Mongolians. When we focused on the genetic diversity of the Hezhen, Tu, Oroqen, Daur, and eastern Mongolians in the right populations in the qpWave analysis, we found they formed a clade with each other, but not with western Mongolians. Indeed, PCA analysis based on the merged 1240K-dataset showed that eastern Mongolians overlapped with Daur, Ewenki, and some of northern Hans, but western Mongolians were separated with them (**Fig. 3B**). Eastern Mongolians were overlapped with the ancient YRB populations from North China and Neolithic people in southern Siberia, suggesting their close genetic affinity. The observed significantly negative values in *f*_*4*_(Eurasians, Eastern Mongolians; YRB farmers, Mbuti) and *f*_*4*_(MP/ARB Hunter-Gatherers; YRB farmers; Eastern Mongolians, Mbuti) further showed that eastern Mongolians shared more alleles with Neolithic millet farmers. We further explored the topologies among Chinese ancients, East Asians in HGDP, Tai-Kadai-speaking Sui and Zhuang, and Hmong-Mien-speaking Miao using TreeMix-based phylogeny. Western Mongolians clustered with Uyghur people, however, eastern Mongolians clustered closely to southern East Asians and YRB ancient populations (**Fig. 3C**). Admixture results in this merged 1240K-dataset also revealed that western Mongolians harbored more western Eurasian ancestry (orange ancestry maximized in Srubnaya, Sintashta and Srubnaya). Besides, Western Mongolians, similar to Yakut people in Siberia, also harbored more ancestry related to the Neolithic MP people (light-blue). Eastern Mongolians harbored more yellow ancestry related to Neolithic YRB farmers and purple ancestry related to Tai-Kadai people (Sui and Hlai). Compared with northern Hans and geographically close northern East Asian ancient people, Mongolians had more western Eurasian orange ancestry (**Fig. 3D**). Compared to historic Mongols in Mongolia, western Mongolians shared more alleles with ancient YRB farmers and modern East Asians related to Sino-Tibetans, Austronesian, Austroasiatic and Tai-Kadai people, as the observed negative values in *f*_*4*_(Late_Mongol, western Mongolians; Eurasians, Mbuti). Differentiated sharing alleles between western and eastern Mongolians were also evidenced via the observed significant asymmetrical-*f*_*4*_ values in *f*_*4*_(western Mongolians, eastern Mongolians; Eurasians, Mbuti). Differentiated allele sharing patterns were also evidenced in the merged HO dataset (**Supplementary Figs. 23∼38**).

**Fig. 3.**
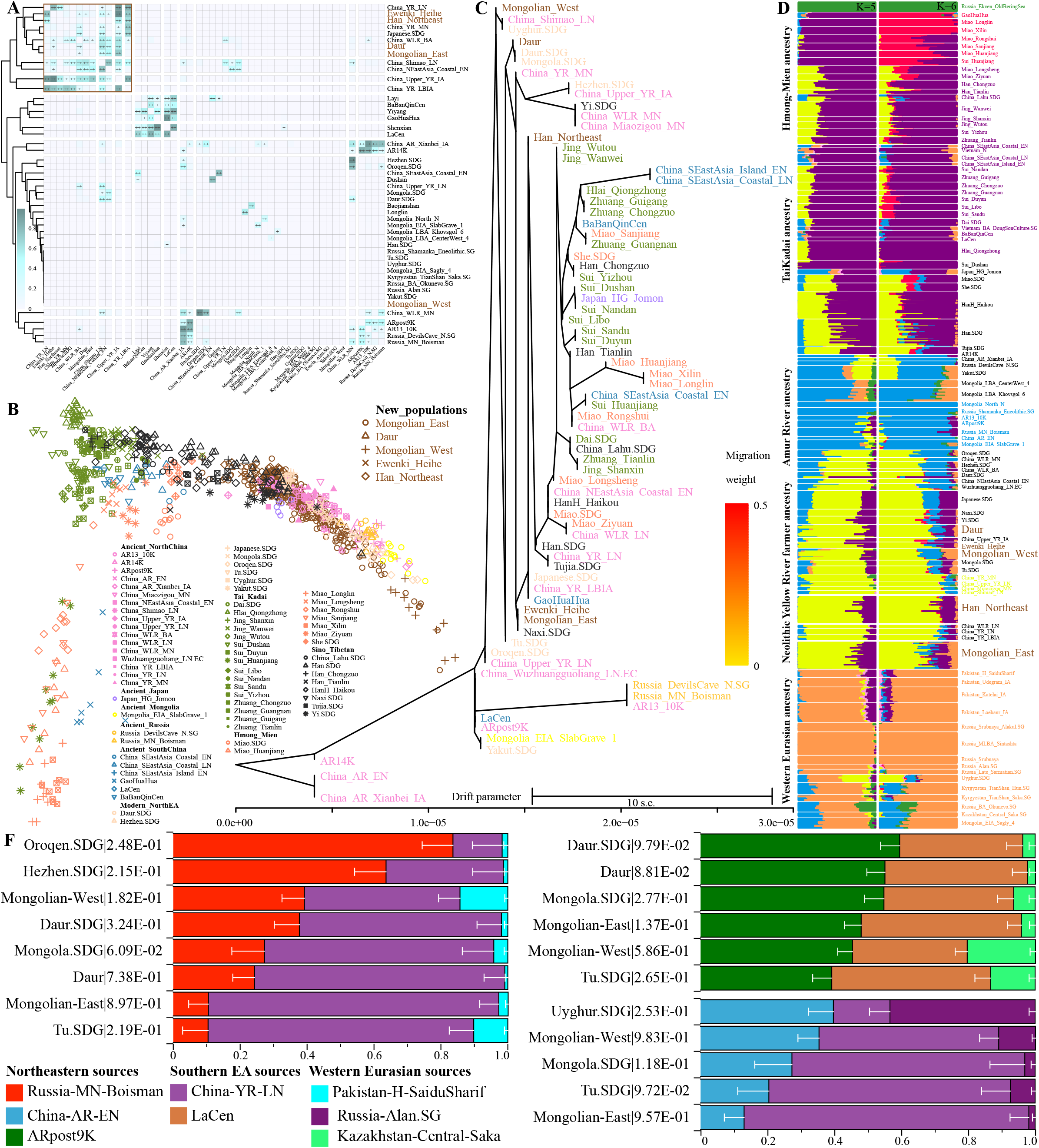
Genetic sources and mixed landscape of eastern Asians based on the merged 1240K dataset (369519 SNPs). (**A**) The p values of rank1 in the pairwise qpWave analysis. P values larger than 0.05 were marked with “++” and larger than 0.01 were marked with “+”. The color in the heatmap showed the detailed p values. (**B**). PCA results among 1073 individuals from 71 modern and ancient populations. Ancient populations were projected. (**C**) TreeMix-based phylogeny without migration events among 71 populations. (**D**) Model-based ADMIXTURE results showed the possible ancestral sources and their corresponding admixture proportion. The breadth of populations is not correlated with the included population size, which was be enlarged or reduced to be better visualization, especially for the eastern Mongolians. (**F**) Three-way admixture qpAdm models with different ancestral sources showed the mixed landscape of northern Mongolic and Tungusic people. The p_rank2 values were presented following the population name and the white bar represented the standard error of the estimated ancestry proportion.

**Fig. 4.**
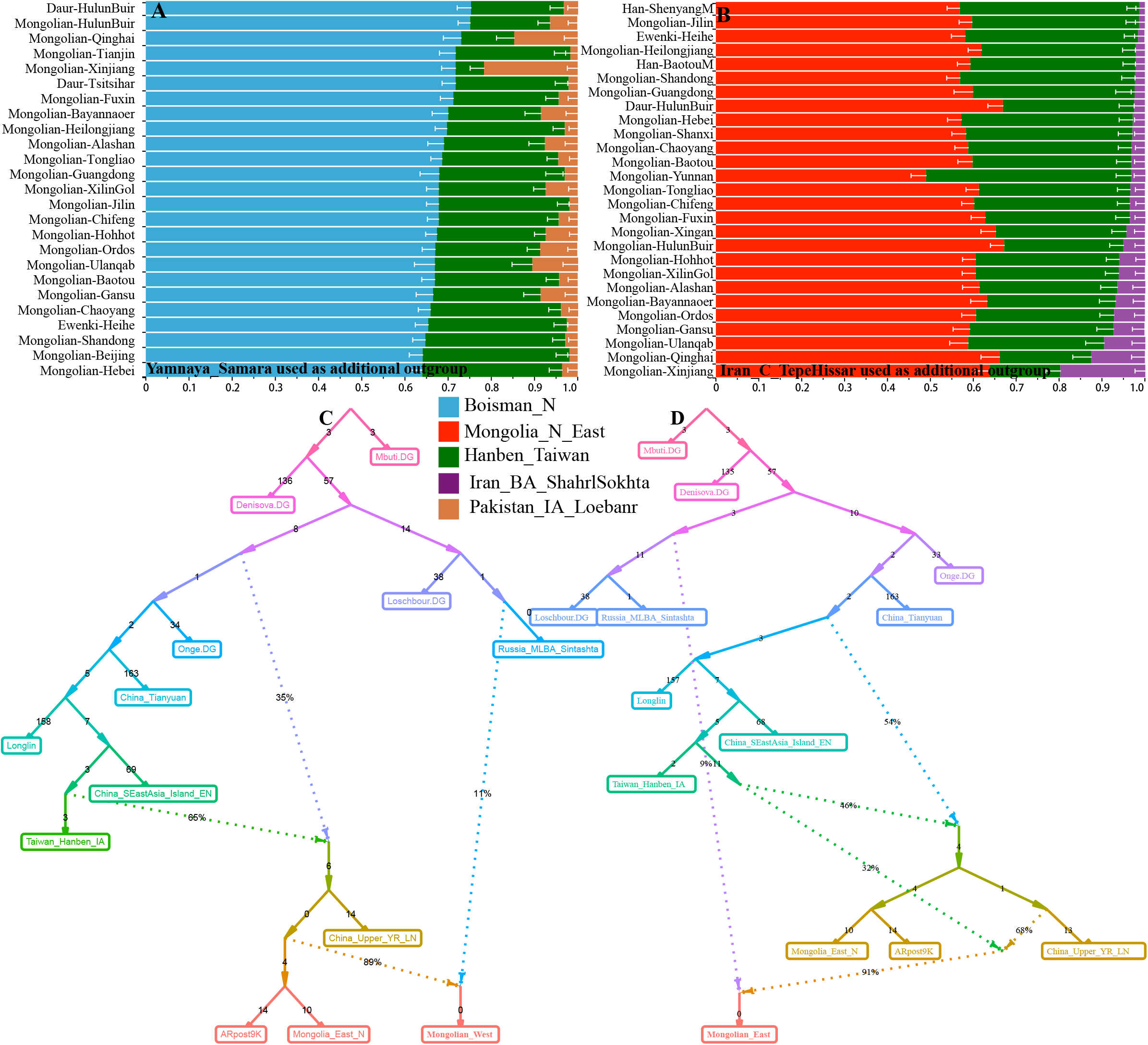
Demographic history of western and eastern Mongolians estimated via the qpAdm and qpGraph. (**A∼B**) The estimated ancestry coefficient of the representative sources in the three-way ADMIXTURE models with Northeast East Asian sources from Mongolian Plateau and Amur River Basin, East Asian source related to Hanben people and western source related Iran and Pakistan ancients. The bar plot denoted the standard error of the admixture proportion. The p values of p_rank2 were larger than 0.05. (**C∼D**) The best fitted qpGraph-based deep population admixture history of western Mongolians (**C**) and eastern Mongolians (**D**)

Considering the complex admixture sources of northern East Asians identified in admixture-*f*_*3*_-statistics, ALDER and GLOBETROTTER, we could successfully fit the admixture model via the three-way admixture models with ancestry from East Asians, Northeast Asians and western Eurasians. As we expected, western Mongolians were modeled as an admixture of more ancestry related to Alan, Saka or Historic SaiduSharif in Pakistan, and the remained ancestry related to Neolithic MP/ARB/YRB people or southern East Asian Guangxi ancient people. We also evaluated the ancestry composition of geographically separated populations using qpAdm-based models (**Fig. 4B**). We found that the three-way admixture models with the aforementioned three ancestral sources could be well-fitted for most of the included studied populations with variable ancestry proportions. We further reconstructed the deed demographic history using qpGraph. Here, we included Mbuti, Denisovan, Onge, Loschbour, Tianyuan and Longshan people to explore the basal model (**Fig. 4B**). We also used the Sintashta, island Neolithic-to-Iron-Age southern East Asians (Liangdao and Hanben), ARpost9k and Mongolia_East_N, as well as upper YRB farmers (Jinchankou and Lajia) as the ancestral source proxies from western Eurasia, Fujian, YRB, MP and ARB. We could find that western Mongolians were fitted via 11% gene flow from steppe Sintashta pastoralists and remained ancestry (89%) from eastern Eurasian ancients related to Neolithic millet farmers (**Fig. 4B**). Focused on the genomic formation of eastern Mongolians, the admixture modeling graph showed that they were produced via three ancestral sources: western Eurasian (0.090), Southern East Asian Hanben (0.291) and upper YRB farmers (0.180). Our results suggested that modern Chinese Mongolic and some of the Tungusic people were formed via the massive population movement and interaction of three ancestral sources related to East Asians, southern Siberians, and western Eurasians.

### Genetic history of Northeast Han Chinese populations

Han Chinese with the largest population size in the world have complex demographic history. Genetic traces extracted from the ancient genomes supported that Han Chinese were formed via admixture of the major ancestry from Neolithic YRB farmer ancestry and geographically different indigenes(20). Xu et al. analyzed the genome-wide SNVs of 114,783 Han Chinese (Han100K) individuals and found six population subgroups: Northwest Hans, Northeast Hans, Central Hans, Southwest Hans, Southeast Hans and Lingnan Hans(25). Our recent genetic survey focused on geographically distinct Hans also identified their differentiated demographic history: Northwest Hans from Gansu harboring more western Eurasian ancestry; Southwest Hans from Sichuan possessing more genetic influence from Tibeto-Burman speakers; Southmost Hans from Hainan having more ancestry related to Tai-Kadai people; Southeast Hans from Fujian possessing more Austronesian-related ancestry; Central Hans from Chongqing, Guizhou, Hubei, and Hunan harboring equal ancestry related to Neolithic YRB millet farmers and southern indigenes, and North Hans in Shaanxi and Shanxi possessing dominant local millet farmer ancestry. Whole-genome sequence data from Taiwan Hans, Singapore and Peranakan Chinese also showed that Hans outside of Mainland China possessed additional genetic admixture from indigenous Austronesian Ami or Malay(28, 29). However, the genetic origin and population structure of Northeast Hans from Inner Mongolia, Jilin, Liaoning and Heilongjiang keep uncharacterized. Northeast China was majorly populated by Mongolic and Tungusic people, but the recent northeastward migration of Hans, such as historic migration events of Chuanguandong and Zouhukou, as well as the established Yuan and Ming dynasties, changed the genetic landscape of this region. Thus, we collected 119 Han Chinese individuals from Baotou (25), Changchun (24), Harbin (24), HulunBuir (21) and Shenyang (25, **Fig. 1A**) to illuminate the genetic formation of Northeast Hans and explore how they interacted with adjoining Altaic-speaking people.

Northeast Hans was localized between North Hans from Henan, Shandong and Shaanxi provinces and Mongolians based on the merged HO dataset and was far away from indigenous Tungusic Ulchi and Nanai (**Fig. 1B∼E**). Five studied Hans shared a similar mixed genetic landscape with North Hans, which harbored major light-blue and light-green ancestry and minor red and dark-blue ancestry. No significant *f*_*4*_-values in *f*_*4*_(Studied Han Chinese1, Studied Han Chinese2; Ancient Eurasian populations, Yoruba) further showed a close genetic affinity within Northeast Hans (**Supplementary Figs. 39**). Han people were also clustered together based on the co-ancestry matrix-based FineSTRUCTURE and shared-IBD-based heatmap (**Fig. 2**). Pairwise qpWave focused on the meta-Northeast Hans and one of modern and ancient northern East Asians showed that Northeast Hans formed one clade with Late Neolithic-to-Iron Age YRB farmers in Henan province, suggesting a close genetic connection with YRB farmers. Genetic variations extracted from the first two components in the merged 1240K dataset further confirmed the close relationship between Northeast Hans and middle YRB farmers, as some of them were overlapped in **Fig. 2B**. Studied Hans were also clustered together in the TreeMix tree and model-based ADMIXTURE results in the 1240K dataset, in which Northeast Hans possessed more purple Tai-Kadai ancestry than Henan ancients but lack light-blue Neolithic MP ancestry (**Fig. 2C∼D**), suggesting they received more genetic influence from southern East Asians than from their ancient counterparts. Different from the Northwest Hans, we could not identify obvious western ancestry in studied Hans based on the model-based ADMIXTURE modeling, which was following confirmed by the limited western Eurasian ancestry estimated in three-way qpAdm admixture models (**Fig. 4A∼B**). The most shared genetic drift with one of geographically Northeast Han is another Han population, following by eastern Mongolian in the outgroup-*f*_*3*_-statistics. We evaluated the ancestral source composition of Hans using admixture-*f*_*3*_-statistics and found that included Hans had a similar admixture landscape as other Chinese Hans, in which one plausible source was from southern indigenous and the other from northern Altaic or Tibeto-Burman people. Results from *f*_*4*_(YRB farmers, Siberian ancients; northeastern Hans, Mbuti)≥3*SE and *f*_*4*_(Hans1, Hans2; Eurasians, Mbuti)≤3*SE showed that northeastern Hans were one genetically homogeneous population, which showed a strong genetic affinity with Neolithic YRB farmers.

Additionally, we assessed the genomic formation of Northeast Hans, as well as geographically close northern minorities (Oroqen, Hezhen, Daur and Japanese), as well as southern Han Chinese and other ethnic groups (Miao, Jing, Tujia and Gelao) from South China using the qpAdm analysis with different representative ancestral sources in the merged 1240K dataset. We first modeled mixture proportion with early ARB ancients as the northern surrogate sources and Late Neolithic Coastal South East Asians (Tanshishan and Xitoucun) as the southern surrogate sources (**Fig. 5A**). Five Northeast Hans were mixed via 0.506∼0.586 ARB ancestry and 0.494∼0.414 southern East Asian ancestry. Compared with Northeast Hans, Northeast minorities harbored more Neolithic northeastern East Asian ancestry ranging from 0.599±0.047 in Heihe Ewenki to 0.851±0.043 in Oroqen. Southern Hans, as expected, had more southern East Asian ancestry ranging from 0.565±0.039 in HGDP Han to 0.583±0.039 in Zunyi Han, and southern ethnically specific minorities also harbored more southern East Asian ancestry ranging from 0.566±0.043 in Tai-Kadai-speaking Gelao to 0.812±0.045 in Tai-Kadai-speaking Zhuang. We following used late Neolithic YRB farmers as the northern representative ancestral source, as China_YR_LN shared the most ancestry with Northeast Hans in the PCA, ADMIXTURE, outgroup-*f*_*3*_-based allele sharing. Only HulunBuir and Harbin Hans could be successfully modeled as major (0.871∼0.842) ancestry related to YRB farmers and minor (0.129 ∼0.158) ancestry related to Guangxi LaCen people. All included populations sampled from the south of YRB could be fitted as 0.133∼0.613 YRB millet farmer ancestry and 0.867∼0.387 southern historic Guangxi ancestry. We further used two ancestral sources respectively from YRB and ARB to model the genetic formation of included northeastern populations (**Fig. 5C**). We confirmed the significant influence of Neolithic ARB ancestry on the formation of the gene pool of Northeast Hans, but Mongolic and Tungusic people possessed more ARB ancestry ranging from 0.086 in Ewenki and 0.881 in Oroqen people. The best-fitted qpGraph models focused on Hans also supported the limited gene flow from western Eurasian into Northeast Hans, in which Northeast Hans were fitted as the 24% ancestry related to Southeast coastal Iron Age Hanben people and 76% ancestry from Neolithic upper YRB farmer ancestry (**Fig. 5D**).

**Fig. 5.**
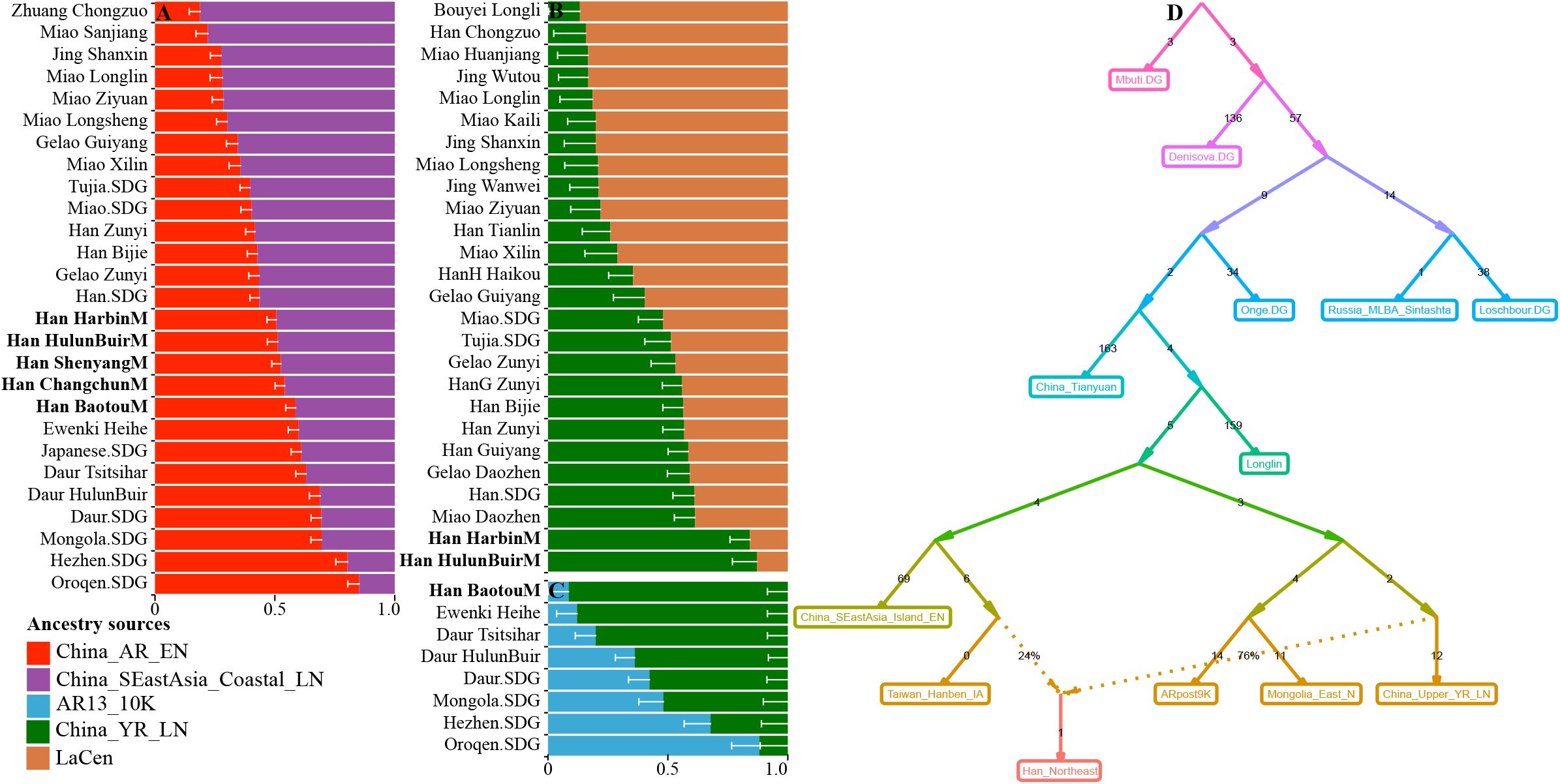
Genetic formation of Northeastern Han Chinese inferred from the genome-wide SNPs data in the merged 1240K dataset. (**A**) Two-way admixture model (Amur Neolithic sources+ Neolithic Coastal southern East Asian source). (**B**)North-to-south admixture model with two sources related to YRB farmers and 2000-year-old LaCen inland southern East Asian showed the differentiated ancestry composition of Han Chinese and southern Chinese indigenous populations. (**C**) Ancestry admixture landscape characterized via the two sources from the Yellow River farmers and Amur River Hunter-Gatherer. All fitted models with the p_rank1 values were larger than 0.05 and the bar plot represented the standard error. (**D**) The deep population formation history of Northeast Hans via the best-fitted qpGraph-based admixture graph.

### Dominant Neolithic Mongolian Plateau ancestry in Tungusic and Mongolic people in Siberia and Amur River Region

Ancient DNA researches from the MP, ARB, Baikal Lake regions and other eastern Eurasian steppe have demonstrated dynamic population admixture history of ancient eastern Eurasian populations. Sikora et al. recovered 34 31,000∼600-year-old ancient genomes and identified three ancient representative populations localized in northeastern Siberia, including Ancient North Siberians, Ancient Palaeo-Siberians, and Neo-Siberians(7). Jeong et al obtained large-scale ancient genomes from Outer Mongolia and Baikal Region and found the initial indigenous MP ancestry, the first genetic communication between western and eastern Eurasian, as well as the repeated and complicated mixed history of multi-ancestral sources since Bronze Age to historic empire periods(9). Mao et al. recently sequenced upper Paleolithic to the Iron Age genomes in the ARB and identified the population migration and replacement in this region 9000 years ago and observed a regional genetic continuity since then(14). Ancient findings from the easternmost Eurasian steppe also identified the genetic stability from 7700-year-old ancient DevilsGate people to modern Tungusic Ulchi and Nanai people. Recent linguistic and genetic evidence also evoked the controversial opinion of the origin of the modern Trans-Eurasian (Altaic) language and corresponding people. Linguists proposed modern Altaic people were originated from West Liao River regions associated with the dissemination of millet agriculture (farming-and-language-dispersal hypothesis). However, recent ancient DNA traces provided another possibility that the population expansion of the Hunter-Gatherer people in the MP dispersed the Proto-Altaic language. Although previous genome-wide SNPs have been used to solve this problem, modern comprehensive and deep population history focused on the genetic variations of modern people should be further conducted to illuminate how previously documented representative sources contributed to modern northern East Asians.

Except for the denser sampling of Mongolians, we also genotyped Mongolic Daur and Tungusic Ewenki and Hezhen people. We flowing combined our data with 25 previously published Tungusic and Mongolic populations in China, Mongolia and Russia to comprehensively characterize the potentially existed differentiated population history and their relationship to the ancient people, as well as the possible association between population migration and the spread of Trans-Eurasian language. HulunBuir and Tsitsihar Daur had a close genetic relationship with each other and clustered together in PCA, ADMIXTURE and FineStructure-based population clustering patterns (**Fig. 1∼2**) in the merged HO dataset. Hezhen and Ewenki were clustered closely with Mongolian people. Similar mixed ancestral sources and landscape of ancestry components were confirmed based on the 1240k dataset via asymmetrical-*f*_*4*_-tests in the form *f*_*4*_(pop1, pop2; Eurasians, Mbuti). Our studied and previously sequenced Hezhen and Daur could be fitted via three-way admixture qpAdm models, in which Hezhen people harbored more ancestry related to Neolithic ARB ancients (**Fig. 3**). Dominant Neolithic ARB ancestry in Mongolic and Tungusic people was further confirmed via the three-way admixture models focused on non-Meta populations in the merged HO dataset (**Fig. 4**).

Eastern Eurasian PCA has revealed the separation between northern and southern Mongolic people as well as between the Mongolic and Tungusic people (**Fig. 1**). Thus, to further comprehensively characterize the landscape of these populations, we collected 235 Altaic-speaking individuals from 26 populations to conduct population structure analyses based on the shared alleles and haplotypes except for Turkic people whose population genetic history has been recently explored(30). We could identify four population clusters respectively associated with the northern (pink circle) and southern (orange) Mongolic people, Japanese and Korean (dark-green), as well as the Tungusic people (yellow) based on PCA patterns reconstructed from the allele frequency spectrum and co-ancestry matrix (**Fig. 6A∼E**). Four ancestral sources were further confirmed via the ADMIXTURE results with four predefined ancestral sources. Here, we also confirmed the mixed genetic structure of Mongolian, Hezhen and Xibo consisting of major southern Mongolian ancestry and some of the Japanese and Tungusic ancestry, as well as Daur and Oroqen composing of four ancestries (**Fig. 6F**). Pairwise Fst within 26 populations first revealed significant genetic differentiation between Tungusic-speaking Ulchi, Negidal and Nanai with others, following showed the differences between Japanese with other references. Siberian Buryat people showed a close genetic relationship with each other, following with Mongols in Mongolia, and distant with southern Mongolic people (**Fig. 6G**). Longer sharing IBD observed in Nanai, Negidal and Ulchi than others showed their unique genetic structure and higher inbreeding phenomenon. Similar patterns of longer sharing IBD fragments were further identified in Siberian Buryat and Mongols. Other Chinese Mongolic and Tungusic people had relatively shorter IBD chunks, suggesting more frequent population movement and admixture (**Fig. 6H**). Differentiated population genetic history of geographically/ethnolinguistically diverse Altaic populations and corresponding clustering patterns were further confirmed via the sharing number of IBD chunks in FineStructure-based dendrogram and TreeMix-based phylogeny (**Fig. 6I∼K**).

**Fig. 6.**
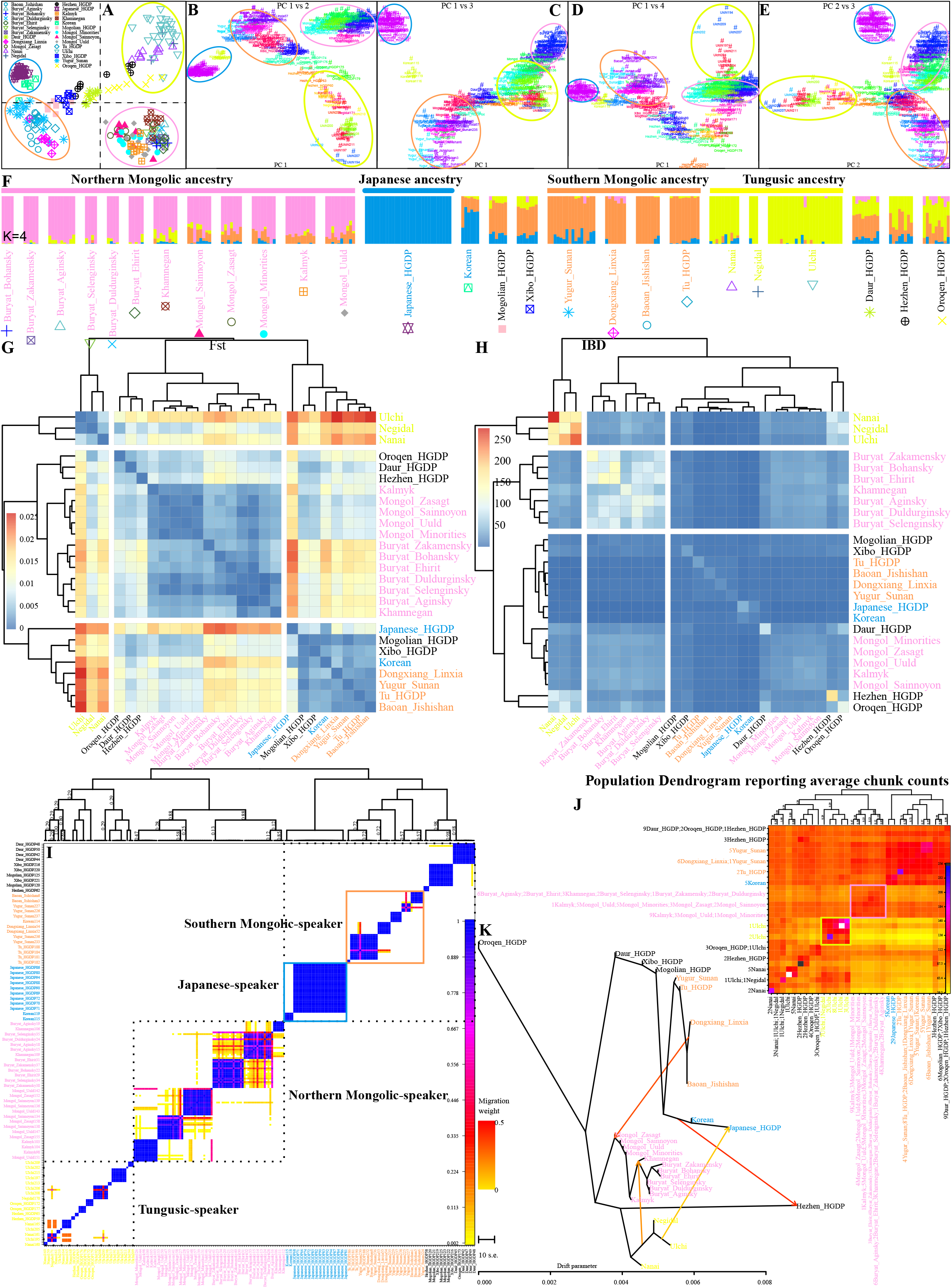
Fine-scale population structure of 238 northern Mongolic and Tungusic people from 26 populations inferred from dense haplotype data of 600K SNPs in the HO dataset. (**A∼E**) Two-dimensional plots based on the allele frequency spectrum (**A**) and coancestry matrix (**B∼E**) showed four genetic clusters within the studied populations. (**F**) ADMIXTURE results for four predefined ancestral sources. (**G∼H**) The pairwise Fst genetic distance and share IBD fragments among 26 Mongolic and Tungusic populations. (**I∼J**) Pairwise coincidence matrix output by FineSTRUCTURE based on the chunk counts and the estimated average chunk counts. (**K**) TreeMix-based phylogenetic relationship with four fitted admixture events.

We following used three-way and two-way admixture models to assess the genetic sources and admixture proportion of 42 Altaic-speaking populations (included Turkic people). All included populations except for four Tungusic-speaking populations (Negidal, Ulchi, Evenk-FarEast and Nanai) could be well-fitted via the three-way admixture models with three representative ancestral sources from northeastern Asia (Russia_OldBeringSea_Ekven, Fofonovo_EN, DevilsCave_N and Boisman_MN), YRB (Xiaowu_MN, Pingliangtai_LN and Haojiatai_LN) and western Eurasia (Russia_Andronovo, Iran_C_TepeHissar and Russia_EBA_Yamnaya_Samara). In the Palaeo-Siberians-YRB-farmers-western-Eurasians models (**Fig. 7A**), Altaic-speaking populations possessed major Neolithic YRB Xiaowu farmer ancestry ranging from 0.462±0.013 in Even to 0.907±0.020 in Yugur (Sunan) and non-ignorable Palaeo-Siberian ancestry (0.053±0.021 in Yugur to 0.339±0.014 in Yukagir) and western Andronovo pastoralist ancestry (0.033±0.011 in Daur to 0.915±0.009 in Veps). Mongolic and Tungusic people harbored more Middle YRB Xiaowu farmer ancestry than Turkic people (0.61∼0.865 versus 0.069∼0.597). Most of included Mongolic and Tungusic populations could be well-fitted via the Neolithic-Mongolians-YRB-farmers-western-Eurasian models (**Fig. 7B∼C**) with dominant Neolithic Mongolia Fofonovo ancestry in non-Chinese ethnic groups (ranging from 0.661±0.070 in Mongol_Uuld to 0.880±0.030 in Khamnegan) and dominant Late Neolithic YRB Pingliangtai ancestry in Chinese groups (0.648±0.057 in Dongxiang to 0.806±0.064 in Yugur). Different from the ancestry composition of Tungusic and Mongolic people, we identified a significant effect of both western Eurasian (Iran_C_TepeHissar: 0.228±0.013 in Kazakh_Aksay to 0.762±0.014 in Nogai) and Neolithic Fofonovo ancestry (0.168±0.060 in Nogai to 0.638±0.02 in Kazakh_Aksay) on the genetic formation of Turkic people. Similar patterns of qpAdm-based genomic affinity were confirmed via the DevilsCave_N-Haojiatai_LN-Yamnaya three-way admixture model (**Fig. 7C**). we also confirmed that Chinese Mongolic and Tungusic people had dominant Neolithic Yellow farmer ancestry via the well-fitted two-way admixture models (**Fig. 7D∼F**). Our findings supported that geographically different Altaic-speaking populations harboring different genetic landscapes with differentiated demographic history: Northern one with dominant Neolithic Mongolia gene flow, southern one possessing significant influence from YRB farmers, and western one harboring extensive genetic admixture with steppe pastoralists and Iran farmers.

**Fig. 7.**
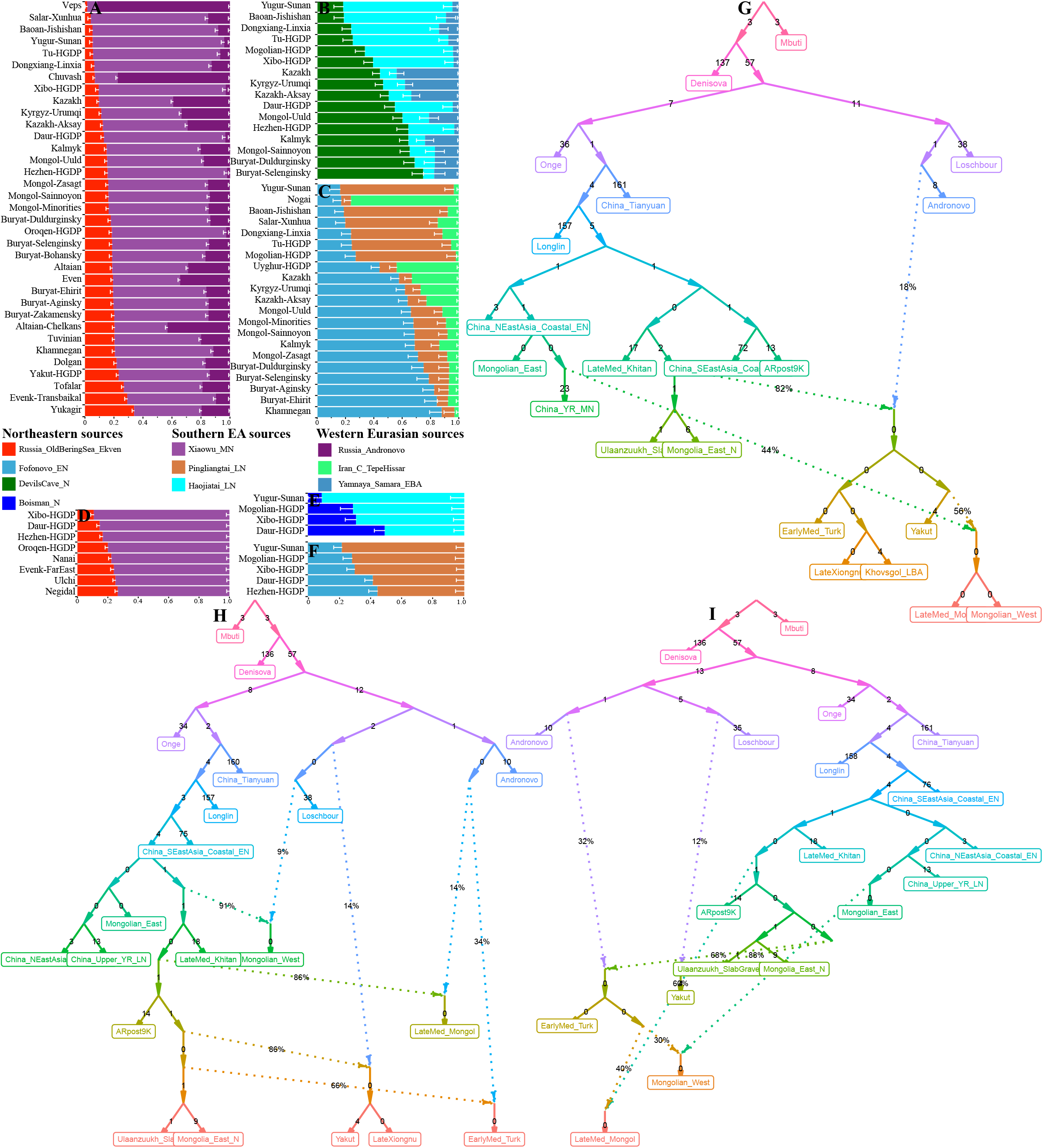
The estimated ancestry admixture coefficient of modern and ancient eastern Eurasians via qpAdm and qpGraph. (**A∼C**) Three-way ADMIXTURE models focused on the Altaic-speaking populations with ancestry from northeastern East Asia, Yellow River Basin and western Eurasia. (**D∼F**) Two-way admixture models with one source from Mongolian Plateau or Amur River Basin and Yellow River Basin. (**G∼I**) The best fitted qpGraph models with different population compositions showed the genetic relationship between Neolithic, historic and modern northern East Asians. The estimated genetic drift (1000×f2 values) was marked along each branch. The dot line represented the admixture events and the admixture proportion was also marked along the dot line.

**Fig. 8.**
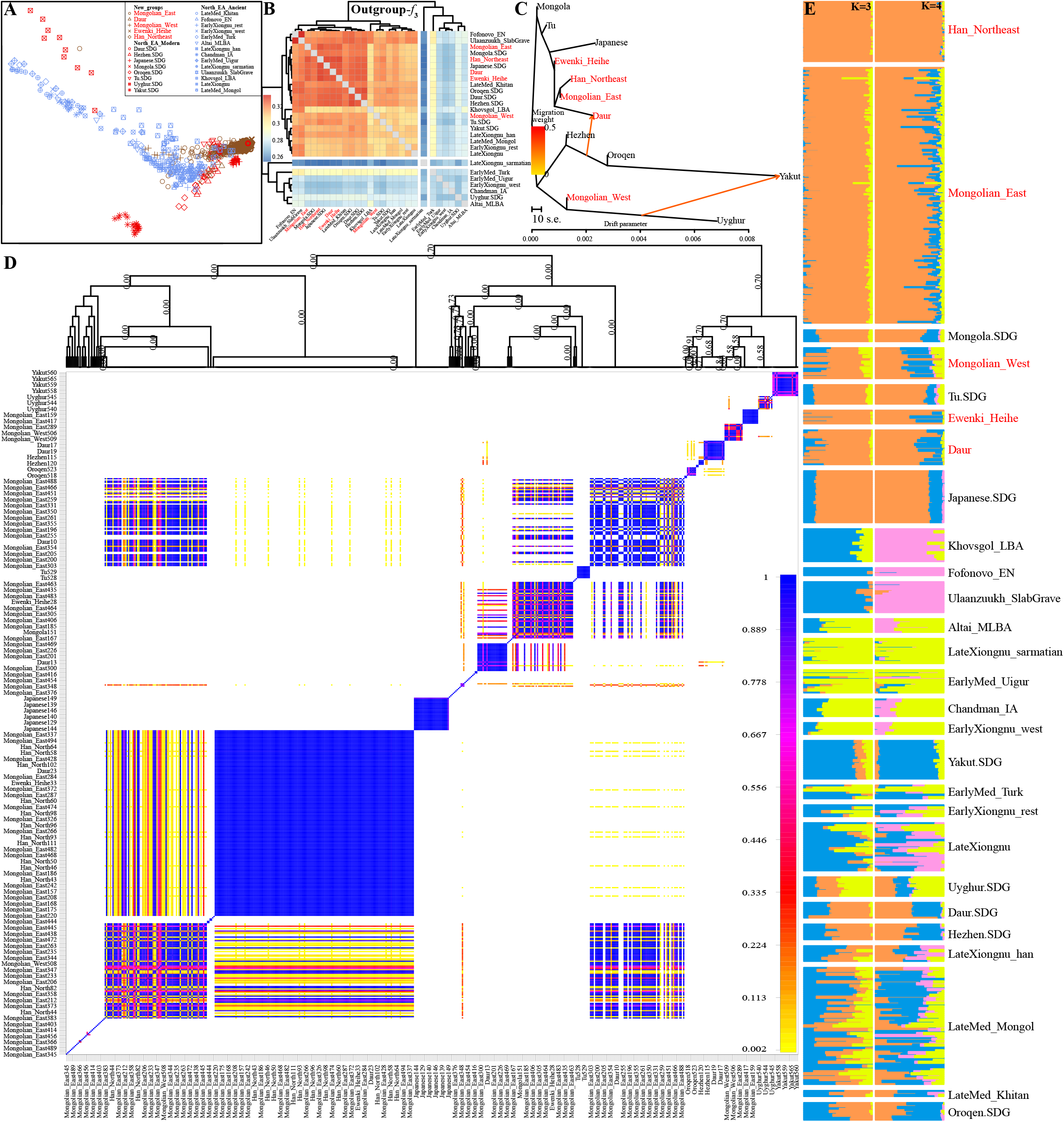
Fine-scale population genetic structure showed the genetic relationship between newly studied Chinese populations. (**A**) Smartpca-based PCA results among 760 individuals from 27 populations. Variations from 566 individuals from 13 populations were used to provide the genetic background. Ancient Mongolian Plateau historic people were projected. (**B**) Pairwise shared genetic alleles estimated via outgroup-*f*_*3*_-statistics. (**C**) Phylogenetic relationship between five studied populations and seven northern East Asian populations included in the HGDP with two admixture events. (**D∼K**) Pairwise coincidence matrix and ADMIXTURE-based results of 13 included modern populations in the merged 1240K dataset.

**Fig. 9.**
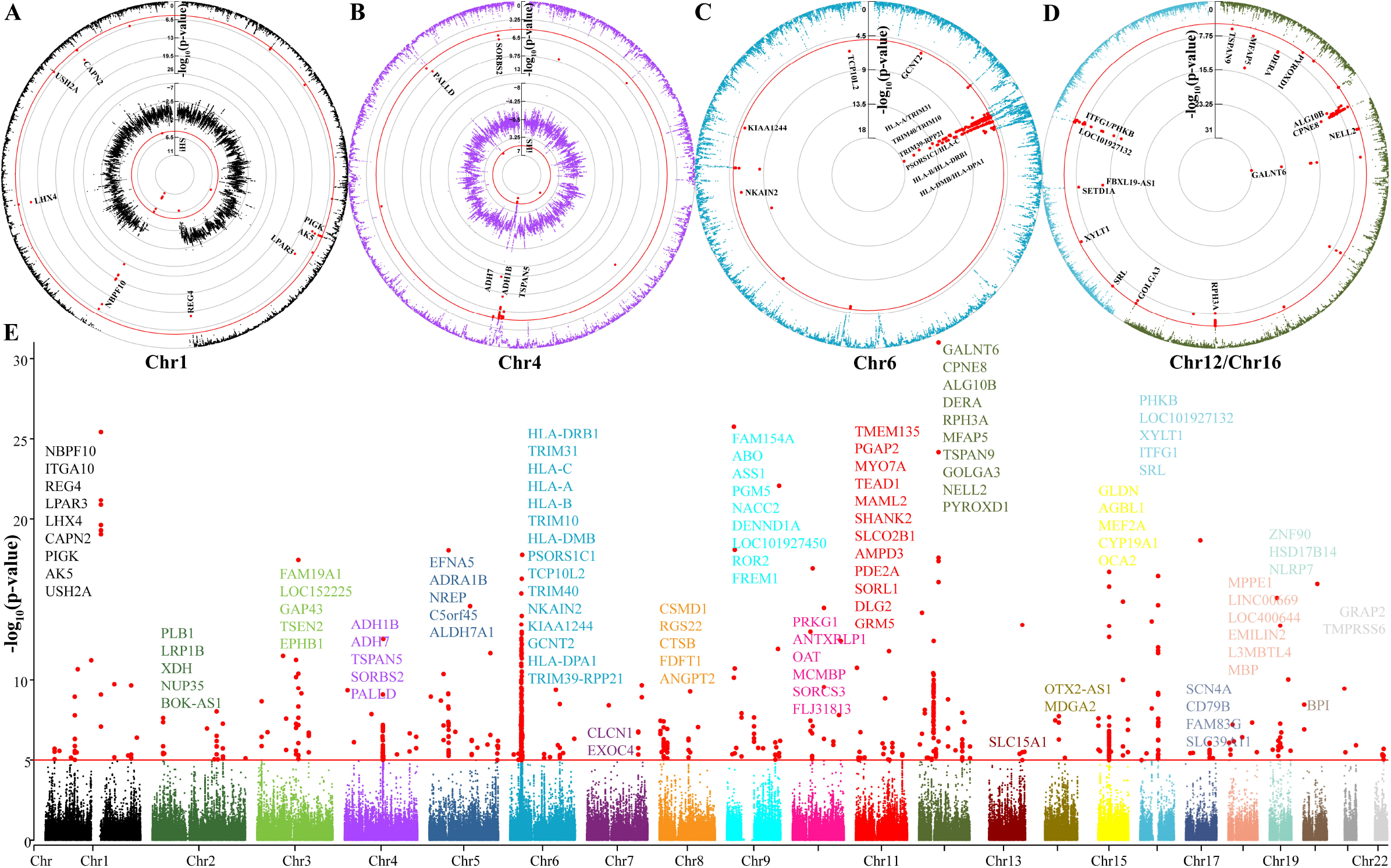
P values of the estimated EXEHH focused on Mongolian populations with the Guizhou Hans as the reference population. (**A∼D**) Circle plots of chromosomes 1, 4, 6 12 and 16 showed the selection signatures. (**E**) Manhattan plots showed all tested candidate loci under natural selection.

### Genetic relationships between northern Chinese populations and historic hierarchical and centrally organized empires in the Eastern steppe

More and more ancient genomes of historic pastoralists were recently reported(9), including Xiongnu, Uyghur, Khitan, Turkic and Mongols. Jeong et al. used the ancient and modern genome-wide data from Siberia and Mongolia to perform the individual-based qpWave analysis and demonstrated that genetic profiles of northern Mongolic-speaking populations have not changed since the Mongol Empire(9). However, the genetic contribution from historic pastoralists in the MP into modern northern Chinese ethnic groups needed to be further characterized.

To explore the shared ancestry between modern East Asians and historic ancient populations, we merged our data with modern Mongolic and Tungusic people in the HGDP dataset, as well as the ancient genomes from the MP, which included some Bronze Age populations and historic Xiongnu, Khitan, Turk, Uigur and Mongol. PCA analysis showed the genetic differentiation between studied populations and ancient Mongolian populations, but some late Medieval Mongols overlapped with eastern Mongolians, and some western Mongolians overlapped with Turk and Xiongnu people (**Fig. 8A**). Shared genetic drift estimated using outgroup-*f*_*3*_-statistics revealed that eastern Mongolians possessed stronger East Asian affinity with northern Hans, Heihe, Daur, Japanese and Hezhen, but distant with western Mongolians (0.3177). Western Mongolians, Daur, Han and Ewenki also shared more ancestry with each other and then with modern northern East Asians not with ancient populations. Heatmap based on the outgroup-*f*_*3*_ matrix showed two major branches, one consisted of ancient populations harboring more western ancestry (Chandman, Turk and Uigur), and the other comprised modern northern Asian populations and Ulaanzuukh_SlabGrave, Xiongnu and Mongol populations (**Fig. 8B**). Admixture signatures inferred from admixture-*f*_*3*_-statistics focused on the eastern Mongolians revealed that the best ancestral sources were from Hans and Mongol ancients. Different from the eastern Mongolians, western Eurasian-related populations (LateXiongnu_sarmatian, Chandman_IA, EarlyMed_Uigur, Altai_MLBA and EarlyMed_Turk) in MP were evidenced as one of the best sources for western Mongolians. Daur and Ewenki possessed the most ancestral sources deriving from the southern source related to Hlai and the northern source linked to Yakut. Ulaanzuukh_SlabGrave people were evidenced as the ancestry surrogate for all studied northern East Asians. Our results from the shared genetic ancestry showed that ancient Xiongnu and Mongols contributed more ancestry into modern northern Tungusic and Mongolic people. But the proportion of ancestry contribution was different, as the differentiated population genetic structure revealed via the TreeMix-based phylogeny and fine-scale substructure inferred from the FineSTRUCTURE (**Fig. 8C∼D**). Under the genetic variations of these included modern and ancient populations, we conducted the model-based ADMIXTURE analysis to explore the ancestry composition. We found three ancestry components maximized in Han Chinese, Neolithic Mongolian ancestry (Fofonovo, Ulaanzuukh and SlabGrave), and western Eurasian-related Xiongnu ancestry respectively. And Yakut ancestry was separated from Neolithic ancestry when we predefined four ancestral sources in the ADMIXTURE-based model fitting. We found a significant genetic difference between Mongolia historic ancients and modern northern Chinese populations. We finally reconstructed the best-fitted qpGraph-based admixture graph and found different genetic lineages between Mongolia historic ancients and modern northern Chinese groups with different population compositions (**Fig. 7G∼I**). We found eastern Mongolians shared the major lineages with YRB farmers and northern Yakut, and western Mongolians shared common lineages with historic Turkic and Xiongnu. Here, we found that western Mongolians shared a similar ancestry history with late Medieval Mongols, which was descended 0.440 ancestry from YRB farmers, 0.4592 from ARB Hunter-Gatherers, and 0.1008 from western Eurasian Andronovo (**Fig. 7G∼H**). We also confirmed that early Medieval Turkic derived 68% ancestry from Neolithic MP people. Ancient Turkic people also contributed 30% of genetic materials to western Mongolians whose remaining ancestry derived from eastern Mongolian-related ancient sources (**Fig. 7I**).

### East Eurasian origin of Mongolic-speaking Kalmyk in northern Caucasus Mountain

More genome-wide-scale population genetic analysis was needed to explore the genetic impacts of the westward expansion of the Mongol empire in the Eurasian steppe on modern central Eurasians. GLOBETROTTER-based admixture sources and dates showed that eastern Mongolian-related haplotype was directly observed in modern central Uyghur, Hazara and others with different proportions, such as 8% in the Turkish(31). Recent findings based on the forensic STRs and SNPs identified the genetic link between geographically distinct Torghut Mongolians and 3000 kilometers away Jalaid Mongols(32). The identified shared alleles between Chinese Mongolians and Hazara also supported the hypothesis of modern Hazara were the descendants of ancient Mongols(33). Kalmyk is one of the Mongolic-speaking populations residing in the Yashkul, Republic of Kalmykia Russia. The genetic origin, population structure and admixture history were formally explored here.

Although the geographical origin of our focused Kalmyk was in North Caucasus, the clustering patterns inferred from PCA and ADMIXTURE showed that the genetic localization of Kalmyk was East Eurasia (**Fig. 1**). The model-based ancestry composition revealed that Kalmyk people possessed major ancestry related to Neolithic ARB people (0.426), and some related to Neolithic YRB Longshan (0.187), western Eurasians (0.196) and Neolithic MP (0.100), and minor related to southern East Asian indigenes. These observed landscapes were consistent with the genetic structure observed in the northern Mongolic people (**Fig. 1F**). The small allele frequency difference between Kalmyk and Xinjiang Mongolian, Siberia Mongols and Buryats (0.0005∼0.0034), suggested their genetic affinity and the common evolutionary history. Outgroup-*f*_*3*_ values further showed that Kalmyk shared the most genetic drift with Tungusic-speaking populations and Neolithic ARB ancient of DevilsCave_N (*f*_*3*_>0.2050). Admixture-*f*_*3*_ analysis showed the ancestral sources of Kalmyk were eastern Tungusic people and western Indo-European groups, such as *f*_*3*_(Nanai, French; Kalmyk)=-38.165*SE. Genetic relationships within Mongolic and Tungusic populations inferred from PCA, TreeMix, heatmap and FineSTRUCTURE (**Fig. 6**) grouped Kalmyk with Mongolians. Kalmyk also shared longer IBD chunks with eastern Eurasians than with others. Formal testing of ancestry proportion estimation confirmed that Kalmyk had an East Asian origin with an admixture of major YRB millet farmers or Neolithic Mongolian ancestry and some western Eurasians (**Fig. 7A∼C**).

### Natural selection signatures in northern East Asian Mongolian and Tungusic people

The cross-population extended haplotype homozygosity (XP-EHH) was calculated using our newly genotyped Mongolians as the target and Guizhou Hans as the reference populations (**Fig. 1**). Many gene-coding candidate selection regions from Chromosomes 1, 4, 6 12 and 16 were identified via the XP-EHH. Here, we highlighted the top 803 SNPs which were clustered and located in different candidate regions of the genomes. Nine SNPs (1: 77854238-77895554) located in the AK5 gene and six SNPs (1: 145444556-145524224) in NBPF10 were the major candidate regions in Chromosome 1, other signatures of natural selection from ITGA10 (3 SNPs) and CAPN2 (3 SNPs) were also identified in this chromosome. Linked haplotype block consisting of fifty-one SNPs located in the CYP19A1 (15: 51502844-51591204) was the longest genomic region under selection in Mongolian populations, which were associated with the etiology of breast cancer. Twenty-six SNPs were located in HLA-C (6: 31236998-31239912) with XPEHH values ranging from -6.121 to -4.805. Twenty-four SNPs located in HLA-A (6: 29910276-29913542) and thirteen SNPs in HLA-B (6: 31321882-31324889) and ten loci in HLA-DPA1 (6: 33035771-32555987) also showed significant natural selection signatures. Twenty-three SNPs in CPNE8 (12: 39050589-39261882) possessed the XPEHH values ranging from -5.692 to -7.260. Thirteen SNPs located in the gastric alcohol dehydrogenase gene (ADH7, 4: 100333932-100349135) showed obvious selection signatures, suggesting that alcohol metabolism in Mongolian people underwent selection during the pastoralist subsistence strategy, which was also evidenced via our estimated iHS results. Besides, at least five functional SNPs in these genes (TRIM31, PHKB, TRIM40, HLA-DRB1, SCN4A, CTSB, GLDN, NBPF10, PGM5, NELL2, GALNT6, ITFG1, PLB1, EPHB1 and NREP) also showed selection signatures, most of which were further comfired via the estimated high iHS values.

### Paternal and maternal admixture history of northern East Asians

Finally, we explored gendered dimensions of the population history of studied northern East Asians. All 362 Mongolians were assigned into 204 terminal matrilineal haplogroups. Haplogroup D was the most predominant lineage (27.07%), followed by B (11.60%), F (10.77%), Z (8.01%), G (7.73%), C (6.91%), A (6.08%), N (5.25%), and M7 (5.25%), other haplogroups (HV, H, I, M8, M9, M10, M11, R, T, U, W, and Y) were sporadically distributed in studied Mongolians with frequencies of no more than 1.66%. Target Mongolic-speaking Daur people were assigned into 19 unique matrilineal lineages. Haplogroup B was the most dominant lineage (26.32%), followed by G (15.79%), F (10.53%), D (10.53%), and A (10.53%), in addition, haplogroups C, N, M8, M11, and R were observed once respectively. Haplogroups B (2/10), C (1/10), D (3/10), F (1/10), M8 (1/10), N (1/10), and R (1/10) were observed in studied Tungusic speakers. All 119 Han Chinese were assigned to 94 terminal matrilineal haplogroups. Haplogroup D was the most common lineage (23.53%), followed by B (12.61%), F (10.08%), M7 (9.24%), N (7.56%), Z (6.72%), G (6.72%), A (6.72%), and C (5.88%), other haplogroups (M8, M9, M10, M11, T, U, and Y) were sporadically distributed in Tungusic individuals with frequencies of no more than 3.36%. Among 175 male Mongolians, we identified 80 terminal paternal lineages with frequencies ranging from 0.0057 to 0.0629 (O2a1c1a1a1a1e: 11). The most frequent paternal haplogroup in target Mongolians was O2 (49.14%), followed by C2 (22.86%), O1 (12.00%), and N1 (6.29%). Furthermore, haplogroups D1, E, I, G, Q, and R were sparsely distributed in studied Mongolian populations. We observed the distributions of haplogroups C2, N1, O1, and O2 in Mongolic-speaking Daur individuals and haplogroups C2, O1, and R in studied Tungusic groups. Sixty Han Chinese men were assigned into 42 diverse paternal lineages. Haplogroup O2 was the most prevalent lineage (45.00%), followed by C2 (26.67%), N1 (10.00%), O1 (10.00%), Q (5.00%), D1 (1.67%), and J (1.67%).

## Discussion

The genetic, archaeological and anthropological findings have demonstrated that the Eastern Eurasian populations had experienced several waves of population turnover. Ancient North Siberians (Yana)(7), ancient northern Eurasians (Mal’ta)(3) and 330,00-year old ARB served as the first batch of pre-LGM Paleo-Neolithic populations, which possessed higher East Asian Tianyuan-related ancestry. Ancient Palaeo-Siberians, Neo-Siberians(7) and other East Asians from YRB, Guangxi and Fujian served as the second batch of founding lineages that worked as the key components to participate in the formation of Holocene to modern northern East Asians. Furthermore, the Yamnaya and Afanasievo people in the Bronze Age had dominated the eastern Eurasian steppe, and these steppe herders gradually expanded eastward to the Altaic Mountains(11, 22) and even reached Shirenzigou of Xinjiang in northwestern China(24) in the Iron Age and central region of Mongolia in the Bronze Age(9). Besides, the ancient genomes from MP also documented the genetic legacy of Iran farmer-related ancestry. Thus, ancient western Eurasians related to northern steppe pastoralists and southern agriculturalists served as the third batch of ancestral sources. The reconstructed deep population history in North-East Asia was dynamic. Recent findings from ARB supported over 13,000-years of genetic stability in ARB, and complex interactions and admixture history between local ancestry and surrounding incoming ancestry were spatiotemporally documented in YRB, MP and Baikal Lake regions. Damgaard et al. provided the first batch of ancient genomes from 74 ancient individuals across Inner Asia and Anatolia and found that there was a distinct East-West cline in the ancient Eurasian steppe populations, namely, eastern European Hunter-Gatherers harbored the highest proportion of western ancestral components, and Neolithic populations (such as Shamanka) of Lake Baikal harbored the highest proportion of eastern ancestral components(12). The genetic findings from 101 newly recovered ancient genomes that dated to between 3000 and 1000 years ago showed that the Bronze Age was a highly dynamic period involving large-scale population migrations and replacements, which also provided supporting evidence for the spread of Indo-European languages during the Early Bronze Age(26). In 2018, Damgaard et al. further sequenced the genomes of 137 ancient humans in Eurasia and found that population substructures existed in the Scythian groups that dominated the Eurasian steppe throughout the Iron Age(10). However, the early population migrations in Eurasia had not significantly affected eastern Eurasia. A recent paleogenomic study of 22 Khovsgol individuals dated to between 5300 to 4700 years ago indicated that dairy pastoralism was introduced through a process of cultural transmission rather than population replacement(23). Highly spatiotemporally sampling ancient DNA from Mongolia further characterized the dynamic history of the Eastern Steppe, which documented the tripartite population structures in the Bronze Age and multiple population admixture in historic periods(9). How these ancient admixture events influenced the modern genetic landscape in North-East Asia needed to be comprehensively explored.

We have presented a comprehensive characterization of genetic variations in 510 northern East Asian Mongolic, Tungusic and Sinitic samples by merging with all available Eastern Eurasian modern and ancient genomes, which is the largest genome-wide genetic study of the Altaic-speaking populations in the boundary regions between South China and Siberia. We documented population stratification within northern East Asians, but all of them possessed the dominant ancestry related to Neolithic MP/ARB people. This documented common genetic legacy in Altaic-speaking populations was consistent with the ancient findings of genetic continuity from 13,000-year-old ARB people to 7700-year-old DevilsGate/Boisman and then to modern Tungusic Ulchi. Shared common ancestry from eastern Eurasian lineage among modern Altaic people was also evidenced via the longer length shared IBD fragments among modern central Asian Turkic people from the southern Siberians(30). Previous Trans-Eurasian language origin hypothesis stated that the language subfamily of Mongolic, Tungusic, Turkic and others shared cultural elements with the Hongshan culture in the West Liao River Basin. If the hypothesis is correct, we expect to observed dominant Neolithic Hongshan ancestry in modern Altaic people in the ADMIXTURE, FineSTRUCTURE, qpAdm and qpGraph-based admixture models. However, we observed dominant Neolithic Hunter-Gatherer-related ancestry from MP/ARB in modern northern East Asians. Considering the co-dispersal of language, gene and culture, as well as the established genetic continuity of deep population history in ARB and surrounding regions, our established genetic landscape in North-East Asia supported Altaic language originated from Northeast Asia. Moreover, archeologically documented stability in the material cultures and anthropologically attested similarity of morphology in MP/ARB further supported our observed dominant local ancestry in northern Altaic people in ARB and neighboring regions. Here, we also noted that we lack more detailed robust evidence to confirm the exact geographical locations of the Proto-Altaic language in Northern East Asia. More complex originated landscapes maybe existed in East Eurasia, such as Robbees et al. provided evidence supporting for new ‘farming Hypothesis’ and against the traditional ‘Pastoralis Hypothesis’ by integrating linguistics, archaeological and genetic phylogenetic findings(34). Deeper and more comprehensive studies based on cultural reconstruction, linguistic diversity, lexicostatistics, Bayesian phylogeography and other interdisciplinary approaches should be conducted to reconstruct a more accurate and complete picture of the evolutionary history of Proto-Altaic people and languages.

Western Eurasian gene flow significantly shaped the genetic structure of western and northern Altaic-speaking populations. Three-way qpAdm models provided good fitness for Turkic and northern Mongolic people, which was also observed in our qpGraph-based phylogenetic framework. ALDER-based and GLOBTTOTER-based date results supported the admixture events that occurred in the historic times. We should pay attention that only single simple or complex models with double dates and sources were considered, which provided the simplest framework of evolutionary history. The truth admixture scenarios may be continuous admixture, which could make a more recent admixture times in the ancient admixture event dating. Ancient population contact between western pastoralists and eastern Eurasians has indeed been attested in the early ancient populations. Early paleo-genomic studies have shown that significant genetic differences existed in ancient Eurasian individuals at different times and geographic scales. But the Yamnaya and Afanasievo populations spread eastward and westward across the Pontic Caspian steppe between 3000 BC to 2100 BC, which was accompanied by the diffusion of the early and middle Bronze Age steppe population-related ancestral components(26). The ancient individuals related to Sintashta, Srubnaya and Andronovo cultures further continued to spread westward and eastward to generate the middle and late Bronze Age steppe population-related ancestral components(10). From 100 BC to 200 BC, Scythians with significantly different genetic structures were active in the eastern, central and western of the Eurasian steppe, respectively(35). These westward movements may also leave genetic legacies in modern eastern Eurasians as identified in our qpAdm and qpGraph admixture models.

With this unprecedented data of Mongolians, we also comprehensively revealed the genetic origins, admixture history, and population structure of geographically diverse Chinese Mongolian populations. Our results showed that the Mongolian people were most closely related to East Asian populations compared with other modern and ancient global populations, especially the northern Tungusic and Mongolic people, suggesting their eastern Eurasian origin hypothesis. Moreover, eastern Mongolians shared most of the genetic makeup with the geographically close Han Chinese populations and Neolithic YRB farmers. Interestingly, eastern Mongolians and Han Chinese were genetically distinguishable from each other, as well as there was a different genetic structure between eastern and western Mongolians, which could attribute to the differentiated Siberian and western Eurasian ancestry in eastern and western Mongolians, and different proportions of YRB ancestry in Northeast Hans. Indeed, all results from allele-shared *f*-statistics and haplotype-shared chromosome painting found remarkable differences in the genetic makeup between Mongolians and Hans, as well as eastern and western Mongolians. Specifically, at least three major ancestral components were identified in the Mongolic and Tungusic people, which were potentially derived from ancestral populations in Western Eurasia, YRB, and MP/ARB. In contrast, two major ancestral components from YRB and southern China were identified in Northeast Hans. QpAdm/qpGraph-based modeling admixture history of northern Altaic and western Mongolians indicated that these three identified ancestral components were derived from Neolithic ancestors or their admixed descendants. The precise source of eastern ancestry for Mongolians was difficult to constrain but our results showed that Neolithic YRB people contributed substantial genetic materials to the eastern Chinese Mongolic populations. The intense population movement and gene flow between ancient Xinjiang people and western Eurasians, as well as the deep connection between Xinjiang local Xiaohe people and South Siberians, have been reported via autosomal and mitochondrial DNA, which provided an explanation for our observed differentiated genetic structure in western Mongolians possessing more ancestry from western Eurasia and ARB/MP.

Our findings also demonstrated that the genetic landscape of modern westernmost Mongolic Kalmyk people was a result of the genetic origin, migration, and admixture history of multiple ancestral sources. Interestingly, among the Mongolic populations in Eurasia, the genetic differentiation between Chinese eastern and western Mongolians and others was even larger than that between Kalmyk and other eastern Mongolic populations, though ARB and Pontic-Caspian steppe were geographically apart. Our results suggested that Kalmyk was genetically closer related to Tungusic people in ARB. According to historical documents, the Kalmyk people were the Mongolian-related nomadic pastoralist regimes who migrated from South Siberia into North Caucasus during the Yuan and Qing dynasties. ADMIXTURE analysis suggested that there was no considerable gene flow between Kalmyk and surrounding populations after Kalmyk people migrated from Siberia into West Asia, which could be attributed to that they speak the Mongolic language, whereas most of the surrounding populations speak Indo-European language. Focused on the relationship between historic pastoral empires and Chinese Mongolians, we found a significant effect of these historic and prehistoric admixture events on the ethnolinguistic diversity of northern and western modern Altaic people, but the limited contribution to Chinese eastern Mongolians compared to the influence from the Han expansion. The Xiongnu confederations grew strong in eastern Eurasia and moved westward about the second or third century BC(27, 36). Subsequently, Turkic, Mongolian and other people successively dominated inner Asia from 600 AD to 1500 AD(27, 36, 37). The genetic impact of the intense east-to-west population movement on the genetic landscape of modern Altaic speakers was stronger than that caused by the north-to-south population migration although historic Chinese Yuan and Qing dynasties were established via the predecessors of modern Mongolic and Tungusic people. Finally, considering the unique and complex history of Mongolic and Tungusic populations, the genome-wide SNP data generated in this study is of great significance for the genetic studies of northern East Asians and serves as a useful control data set for genetic association studies.

## Conclusions

Our population genomic results suggested a complex scenario of admixture history of Altaic-speaking populations in North-East Asia via comprehensively analyzing the newly generated genome-wide SNP data of 31 Mongolic-speaking, two Tungusic-speaking, and five Sinitic-speaking populations in China. The results showed that there were significant population substructures within Altaic-speaking populations (northern and southern Mongolic, Tungusic, and Turkic-speaking populations) and between eastern and western Mongolians. All Altaic-speaking populations were a mixture of dominant Siberian Neolithic ancestry and non-negligible YRB ancestry, suggesting that Altaic-people and their language were more likely to originate from the Northeast Asia (mostly likely the ARB and surrounding regions as the primary common ancestry identified here) and further experienced influence from Neolithic YRB farmers. All Altaic people but eastern and southern Mongolic-speaking populations possessed a high proportion of West Eurasian-related ancestry, in accordance with the linguistically documented language borrowing in Turkic language. Our findings supported the ‘‘ARB-origin hypothesis’’ over the ‘‘Baikal-origin hypothesis’’ for the origin of Mongolic and Tungusic-speaking populations, as they possed documented dominant ARB ancestry. Moreover, the genetic makeup of Mongolic-speaking populations, especially southern, central and eastern Mongolic-speaking populations, harbored more YRB Neolithic farmer ancestry and a stronger genetic affinity with modern Northeast Hans, suggesting the extensive genetic admixture between Chinese Mongolians and adjacent Hans. Finally, we identified a close genetic connection between the Mongolic-speaking populations from North Caucasus and ARB, suggesting long-distance migration of Altaic-speaking people since the Mongolian Empire periods, and these Mongolic groups kept relatively genetically isolated with surrounding Indo-European-speaking populations.

## Materials and methods

### Ethics statement

This study was approved by the Medical Ethics Committee of Xiamen University. Informed consent was obtained from all included individuals before the saliva or blood collection. The procedures of sample collection and experiment of this research were in accordance with the recommendations provided by the revised Helsinki Declaration of 2000(38). All subjects were required to be indigenous self-declared ethnic groups.

### Sample collection and genotyping

The reported dataset consisted of 510 unrelated subjects from 31 Mongolic-speaking populations (Mongolian: 362, HulunBuir Daur: 9, and Tsitsihar Daur: 10), two Tungusic-speaking populations (Heihe Evenki: 8 and Jiamusi Hezhen: 2), and five Sinitic-speaking Han Chinese (119, **Fig. 1**). Large-scale geographically different Mongolians were obtained from 29 groups: Jilin (18), HulunBuir (22), Chaoyang (21), XilinGol (19), Xinjiang (8), Chifeng (43), Heilongjiang (21), Hohhot (38), Tongliao (42), Fuxin (13), Xingan (18), Shenyang (9), Ordos (8), Nanyang (6), Beijing (9), Dalian (3), Qinghai (3), Alashan (3), Shanxi (8), Hebei (7), Tianjin (4), Gansu (3), Yunnan (4), Jiangsu (4), Shandong (12), Bayannaoer (5), Guangdong (2), Ulanchabu (2), and Baotou (7). Han individuals were collected from five northeastern Chinese populations: Baotou (25), Shenyang (25), Changchun (24), Harbin (24), and HulunBuir (21). PureLink Genomic DNA Mini Kit (Thermo Fisher Scientific) was used to isolate genomic DNA from saliva or anticoagulant-treated peripheral blood samples following the manufacturer’s recommendations. Quantifiler Human DNA Quantification Kit (Thermo Fisher Scientific) and Applied Biosystem 7500 Real-time PCR System (Thermo Fisher Scientific) were used to quantify the DNA concentration. Genotyping of 529,790 variants, including 18,711 parentally lineage informative single nucleotide polymorphisms (LISNPs), 4,448 maternally LISNPs, and 506,616 autosomal and X-related SNPs, was carried out using the Affymetrix WeGene V1 Arrays.

### Data purification

We used PLINK v1.90b6.13 with the parameters of genotyping success large than 95% and the minor allele frequency large than 0.05% to perform quality control. We used PLINK and King to identify the close genetic relationships within degree relatives and ran Genome-wide Complex Trait Analysis (GCTA) version 1.92.2 to remove outliers. After quality filtering, we kept 478 individuals in the following population genetic analysis.

### Reference population datasets

For comparative purposes, we merged our data with publicly available modern and ancient Eurasians included in the 1240K, HO, and other recent public databases into three datasets: Affymetrix dataset, the merged HO dataset, and the merged 1240K dataset. The reference populations in the Affymetrix dataset consisting of Chinese populations which were genotyped using the same array in our groups, which included two Hainan populations (Han and Hlai)(39), 14 Tai-Kadai-speaking populations (three Jing, four Zhuang, and seven Sui groups) and seven Hmong-Mien-speaking Miao from Guangxi province(40). The reference groups in the merged HO dataset included 5081 individuals included in the 1240K database and 7744 individuals from the 1240k+HumanOrigin database obtained from David Reich Lab(1, 7, 13, 17, 41). These datasets included all publicly available ancient DNA data published before 2021 and 2054 individuals from the 1000 Genomes Project, 300 individuals from 142 worldwide populations from Simons Genome Diversity Project, and 928 sequence genomes in the sequenced HGDP. Genotype data obtained using the Affymetrix Human Origins were also included here(42-44). The used reference populations in the merged 1240K dataset included all publicly available ancient genomes and the sequenced genomes in the HGDP project(45).

### PCA and Model-based ADMIXTURE analysis

We used PLINK v1.90b6.13(46) and SmartPCA(47) to conduct Eurasian and regional PCA in the context of modern and ancient Eurasians or eastern Eurasians with one additional parameter (Project: YES). Ancient individuals were projected onto the two-dimensional scaling plots. To characterize the individual and population ancestry compositions and reconstruct population genetic history, we used genetic clustering algorithm implemented in ADMIXTURE, one STRUCTURE-like but applying the maximum likelihood-based approach to conduct the model-based clustering analyses in the population genetic and evolutionary genetic studies (48). The analysis was performed based on the Eurasian dataset and other sub-dataset comprising regional East Asians. Plink v1.9 (46) was used to exclude SNPs in strong Linkage Disequilibrium (LD) with the following parameters settings: pairwise SNP loci r^2^ > 0.4; window: 200 SNPs; and sliding windows: 25 (--indep-pairwise 200 25 0.4). A total of 199,753 variants out of 346,634 variants passed filters and quality control (QC) and remained in the merged 1240K. 90,965 out of 120,894 variants passed filters and QC in the merged HO. We assumed the number of predefined ancestry populations (K values) ranging from 2 to 20 and ran unsupervised ADMIXTURE with the 10-fold cross-validation (--cv=10). 100 randomly seeded runs were employed, and the log-likelihood scores and cross-validation errors were used to find the most appropriate K values. Ancestry compositions among modern and ancient people were conducted using the aforementioned settings.

### Three-population test (admixture f_3_-statistics and outgroup-f_3_-statistics)

All possible pairs of source populations in three datasets were used to conduct the admixture-*f*_3_-statistics in the form *f*_3_(Source1, Source2; Studied populations) using the qp3Pop in the AdmixTools (49). Formal tests of admixture with statistically negative *f*_3_-statistic values (Z≤-3) indicated that target populations were an admixture of two predefined ancestral populations. Genetic affinity was estimated using outgroup-*f*_*3*_-statistics in the form *f*_*3*_(reference, studied populations; Mbuti).

### Four-population test (f_4_-statistics)

We employed the formal test of the four-population test (also called *D-statistics* or *f*_*4*_*-statistics*) implemented in the ADMIXTOOLS(49) to test whether our studied populations harbor more shared alleles or genetic drift with targeted reference populations than other worldwide reference populations. We used the statistics in the form *f*_*4*_*(W, X; Y, outgroup)*, where W represents the Asian populations belonging to the Tungusic-, Mongolian-, Turkic-, Hmong-Mien-, Tai-Kadai-, Sinitic-, Tibeto-Burman-, Austronesian-, and Austroasiatic-speaking populations, X represents worldwide reference populations, and Y represents studied populations. The central African of Mbuti was used as the outgroup. Generally, the statistical index was calculated as the differences between counts of BABA sites and counts of ABBA sites divided by the sum of counts of BABA sites and counts of ABBA sites. The null hypothesis is that W and X form a clade and descend from a homogeneous ancestral population separated from Y and Outgroups. Thus, there are no differences in the rate of allele sharing existing with Y, and the D-statistic should be expected to be 0. If there is an excess of shared alleles or gene flow between either or both pairs (W, outgroup) or (X, Y) since separation from the others, a negative D value will be produced. Similarly, a positive D value can be obtained if a large number of the shared allele or gene flow exist between (X, Outgroup) or (W, Y) since separation from the others. Standard errors were estimated using the weighted block jackknife with the block size of 5Mb in a single Hotelling’s T^2^ test. Z-scores were computed as the ratio between D and standard error. |Z| > 3 was regarded as significantly different from 0 and the null hypothesis could be rejected.

### TreeMix and qpGraph

We used the TreeMix version 1.12(50) to explore the topology of included populations with gene flow events ranging from 0 to 10. Phylogenetic trees with Yoruba as the toot and un-root trees were established. We used ADIXTOOLS2 to explore the best qpGraph-based models with different population compositions. We also reconstruct the neighboring trees used Mega 7.0(51) based on the pairwise Fst and inverse outgroup-*f*_*3*_. PCA and MDS based on the genetic distance matrixes were conducted via the IBM SPSS Statistics 21(52).

### Mitochondrial DNA haplogroup and Y-chromosome haplogroup assignment

A total of 18,435 phylogenetic informative Y-chromosomal SNPs and 4,418 phylogenetic mitochondrial-related SNPs were used to classify the haplogroups using an in-house script.

### Pairwise qpWave and qpAdm-based admixture models

We merged geographically and ethnically defined populations into new genetically homogeneous populations (Mongolian_East, Daur, Mongolian_West, Ewenki_Heihe, and Han_Northeast) and performed qpWave analyses(49) to explore the distribution of the *f*_*4*_-matrix via rank tests. Nine continental-representative populations were used as outgroups (Mbuti, Ust_Ishim, Kostenki14, Papuan, Australian, Mixe, MA1, Onge, and Atayal). We also conducted pairwise qpWave analysis used the new-studied populations and previous sequenced HGDP Mongolic and Tungusic populations (Daur, Hezhen, Tu., Yakut, Mongolian, and Oroqen) as the right populations.

### IBD segments inference and ROH

The different datasets were phased using ShapeIT and then we used Refined IBD(53) to characterize the identical by descent (IBD), population-level shared IBD were corrected via the product of sample size of the focused population pairs. We used Plink to calculate the consecutive stretches of homozygous SNPs (ROH).

### Estimating admixture times with MALDER

We estimated admixture times using MALDER(54) with the default parameters or with two additional parameters (mindis: 0.0005 and jackknife: YES). Admixture mediated linkage disequilibrium can be decay with the recombination that occurred after the initial admixture. We only captured the simple characteristics of the admixture process with the assumption of a single pulse-like admixture. We used ancestral sources from southern East Asians, eastern Siberians, and western Eurasians as the possible ancestral proxy sources.

### Chromosome painting and FineSTRUCTURE

To infer finer-scale scenarios of the admixture landscape, we used ChromoPainterv2 to paint the targeted chromosome using all other included donor chromosomes based on the phased haplotype data(55). Chromosome painting could identify haplotypic distribution to further perform admixture dating and population identification. We conducted chromosome painting using the merged 1240K dataset, Altaic people in the HO dataset, and the Affymetrix dataset. Model-based Bayesian clustering instrumented in the FineSTRUCTURE was used to identify population substructure. FineSTRUCTURE version4 and FineSTRUCTURE R scripts based on the reconstructed coancestry matrix were conducted to dissect the fine-scale population structure via heatmap, clustering dendrogram, and PCA.

### ChromoPainterv2 and GLOBETROTTER admixture modeling

R program of GLOBETROTTER(31) was used to identify and date admixture events based on the merged 1240K dataset, which included around 366K genome-wide SNPs. Both full analysis and regional analysis were conduct based on the estimated coping vectors from ChromoPainterv2. GLOBETROTTER used the sampled surrogated populations to paint the haplotypic mixture of the truth unsampled source populations and then to identify whether the targeted Altaic populations descend from any north-to-south or west-to-east admixture events (uncertain, no admixture, single admixture models, or complex admixture models with multiple dates or sources), and -if so -it can statistically calculate precisely when the admixture events occurred and the admixing sources involved.

### Nature selections signatures identifying

We used to selscan and Rehh to calculate the iHS and XP-EHH as the selection indexes and then annotated the identified selection candidate loci using the PLINK. We highlighted the top selection SNPs using the Manhattan.

## Data availability

The variation data reported in this paper can be shared via personal communication with corresponding authors. We make the data available upon request by asking the person requesting the data to agree in writing to the following restrictions: 1, the data can be only used for studying population history; 2, the data cannot be used for commercial purposes; 3, the data cannot be used to identify the sample donors; 4, the data cannot be used for studying natural/cultural selections, medical or other related studies.

## Acknowledgments

This study was supported by the National Natural Science Foundation of China (31801040), Nanqiang Outstanding Young Talents Program of Xiamen University (X2123302), China Postdoctoral Science Foundation (2021M691879), Fundamental Research Funds for the Central Universities (ZK1144), and Science and Technology Program of Guangzhou, China (2019030016).

