## Supplementary Figures for "Genomic insights into the differentiated population admixture structure and demographic history of North East Asians"

### Supplementary Figures S1~39:

**Genomic insights into the differentiated population admixture structure and demographic history of North East Asians**

**Guanglin He[1,2,3*,#], Mengge Wang[2,4,5,*], Xing Zou[2,6], Renkuan Tang[7], Hui-Yuan Yeh[3], Zheng Wang[2], Xiaomin Yang[1], Ziyang Xia[1], Yingxiang Li[1,8], Jianxin Guo[1], Rui Wang[1], Jing Liu[2], Kongyang Zhu[1], Jing Chen[9], Meiqing Yang[9], Qu Shen[1], Jinwen Chen[1], Jing Zhao[1], Hao Ma[1], Lan-Hai Wei[1], Ling Chen[10], Changhui Liu[4], Chao Liu[4,5,10,#], Gang Chen[11,#], Yiping Hou[2,#], Chuan-Chao Wang[1,12,13,#]**

[1]Department of Anthropology and Ethnology, Institute of Anthropology, National Institute for Data Science in Health and Medicine, State Key Laboratory of Cellular Stress Biology, School of Life Sciences, State Key Laboratory of Marine Environmental Science, Xiamen University, Xiamen, 361005, China

[2]Institute of Forensic Medicine, West China School of Basic Science and Forensic Medicine, Sichuan University, Chengdu, 610041, China

[3]School of Humanities, Nanyang Technological University, Nanyang Avenue, 639798, Singapore

[4]Guangzhou Forensic Science Institute, Guangzhou, 510080, China

[5]Faculty of Forensic Medicine, Zhongshan School of Medicine, Sun Yat-sen University, Guangzhou, 510080, China

[6]College of Basic Medicine, Chongqing University, Chongqing, 400016, China

[7]Department of Forensic Medicine, College of Basic Medicine, Chongqing Medical University, Chongqing, 400016, China

[8]AnLan AI, Shenzhen, 518000, China

[9]Department of Forensic Medicine, Guizhou Medical University, Guiyang, 550000, China

[10]Department of Forensic Genetics, School of Forensic Medicine, Southern Medical University, Guangzhou, 510515, China

[11]Hunan Key Lab of Bioinformatics, School of Computer Science and Engineering, Central South University, Changsha, 410075, China.

[12]School of Basic Medical Sciences, Zhejiang University School of Medicine, Hangzhou, 310000, China

[13]Institute of Asian Civilizations, Zhejiang University, Hangzhou, 310000, China

*These authors contributed equally to this work and should be considered co-first authors.

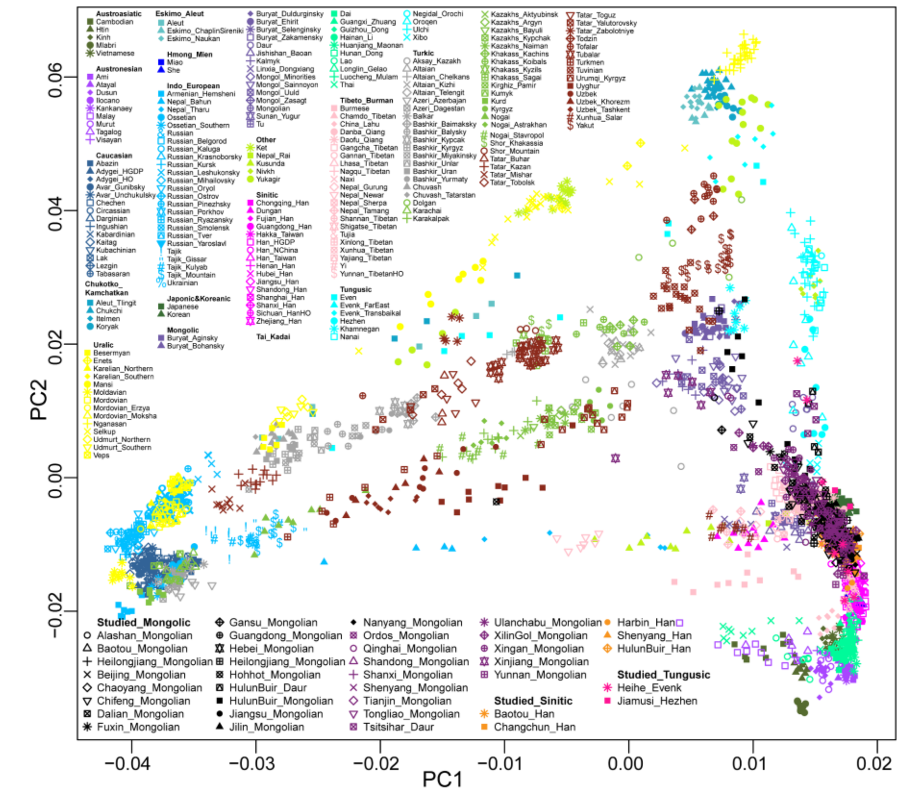


**Supplementary Fig. 1** Genetic affinity between newly studied populations and reference Eurasian populations revealed by principal component analysis.

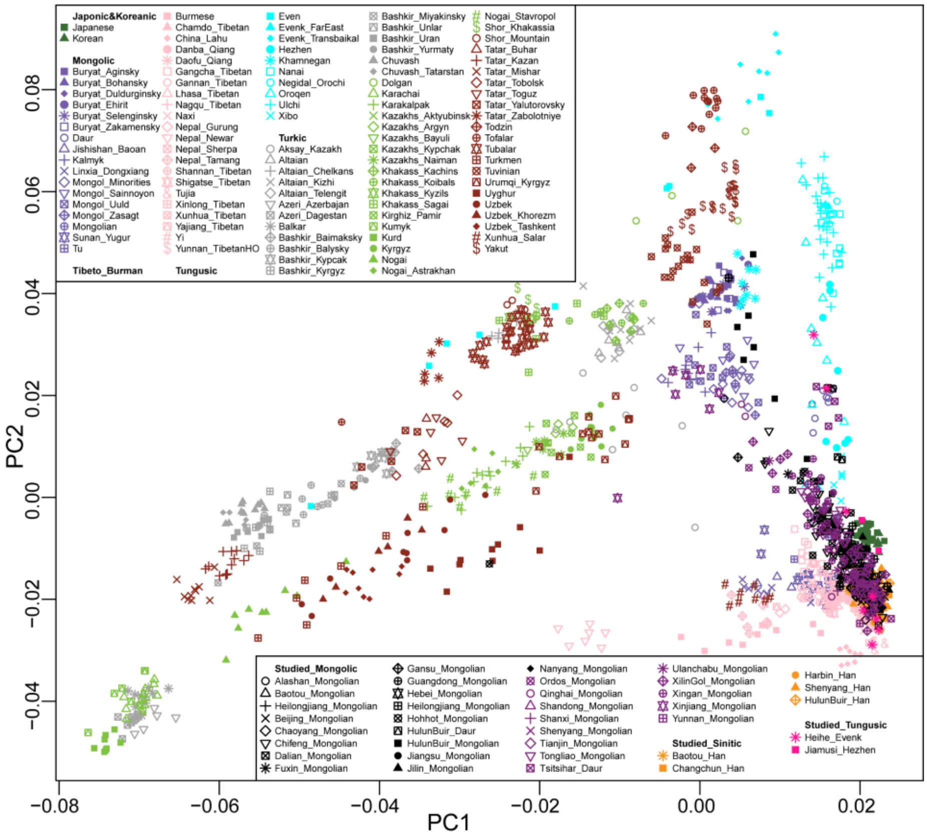


**Supplementary Fig. 2** Genetic affinity between newly studied populations and extracted Tibeto-Burman and trans-Eurasian-speaking populations revealed by principal component analysis.

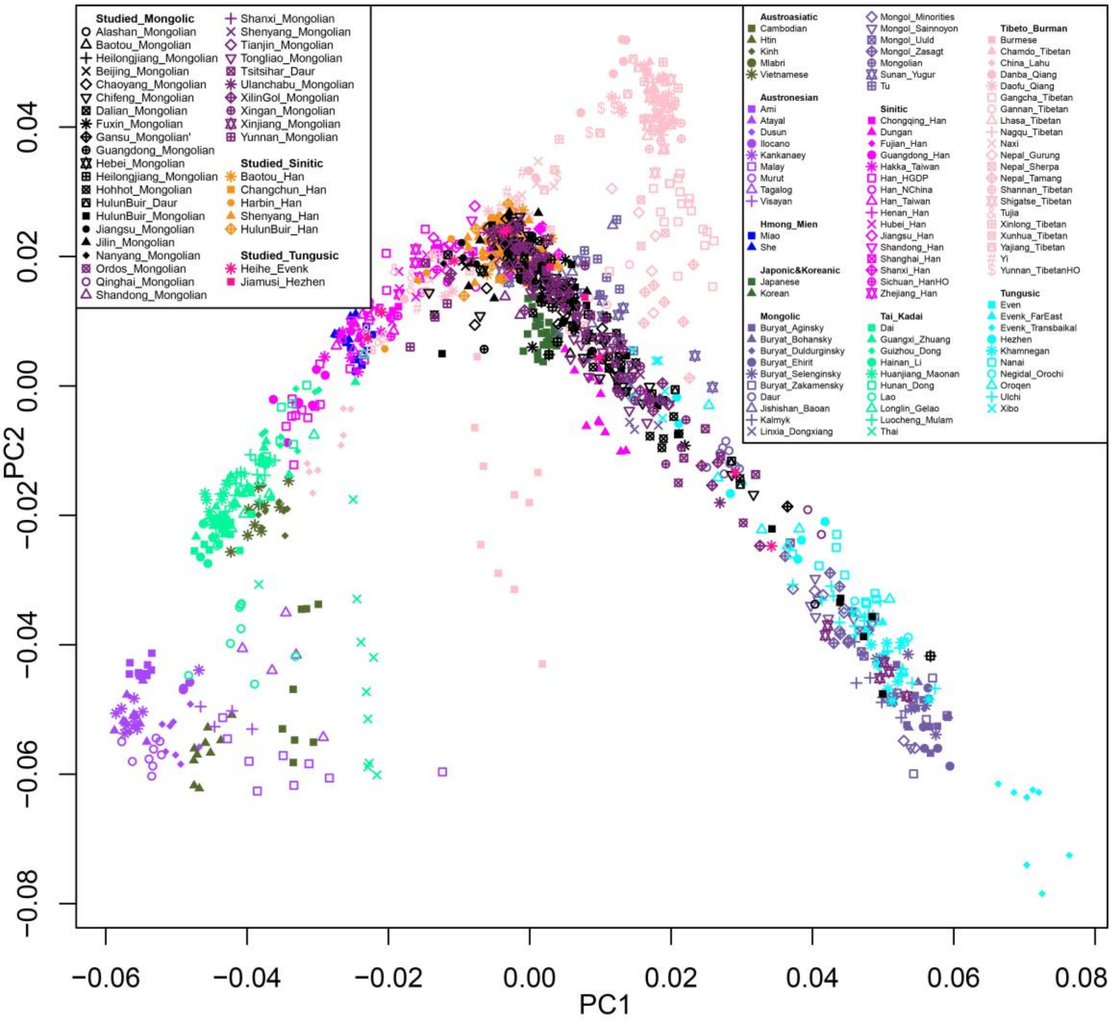


**Supplementary Fig. 3** Genetic affinity between newly studied populations and reference East Eurasian populations revealed by Principal Component Analysis.

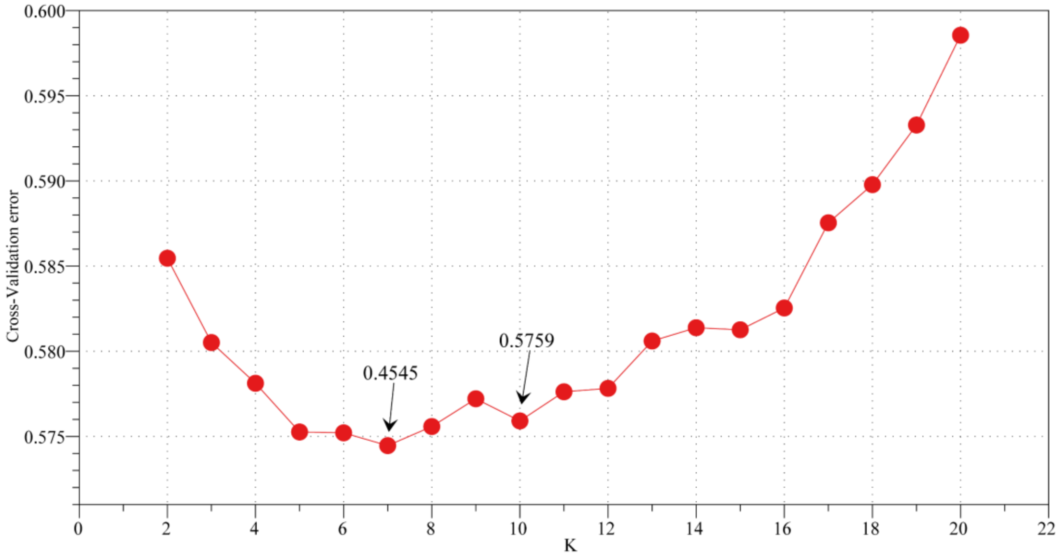


**Supplementary Fig. 4** The cross-validation error values for ADMIXTURE analysis. The optimal k value is 7.

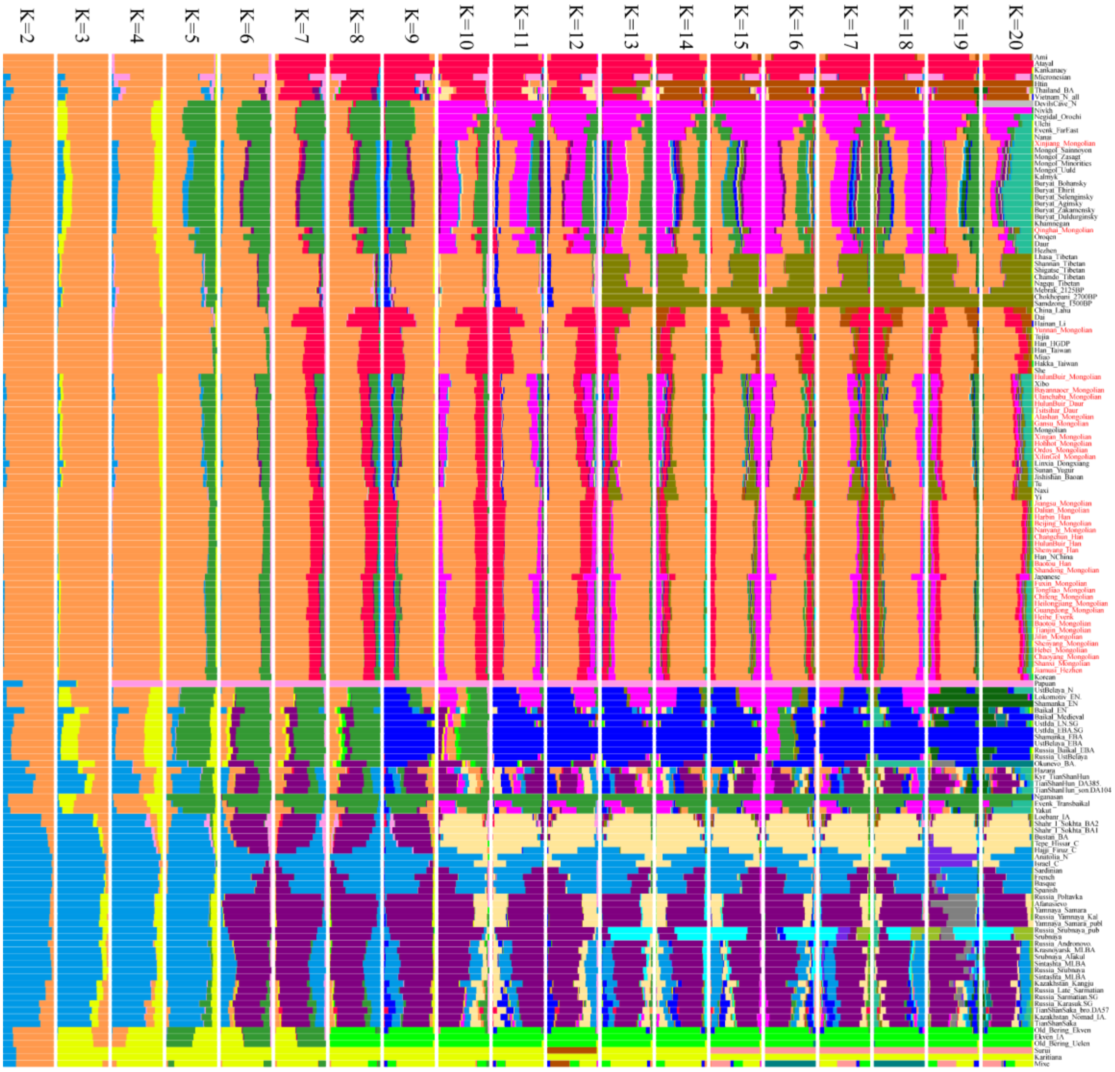

**Supplementary Fig. 5** Mean ancestry composition of newly studied populations and reference ancient and modern non-African populations revealed by ADMIXTURE analysis.

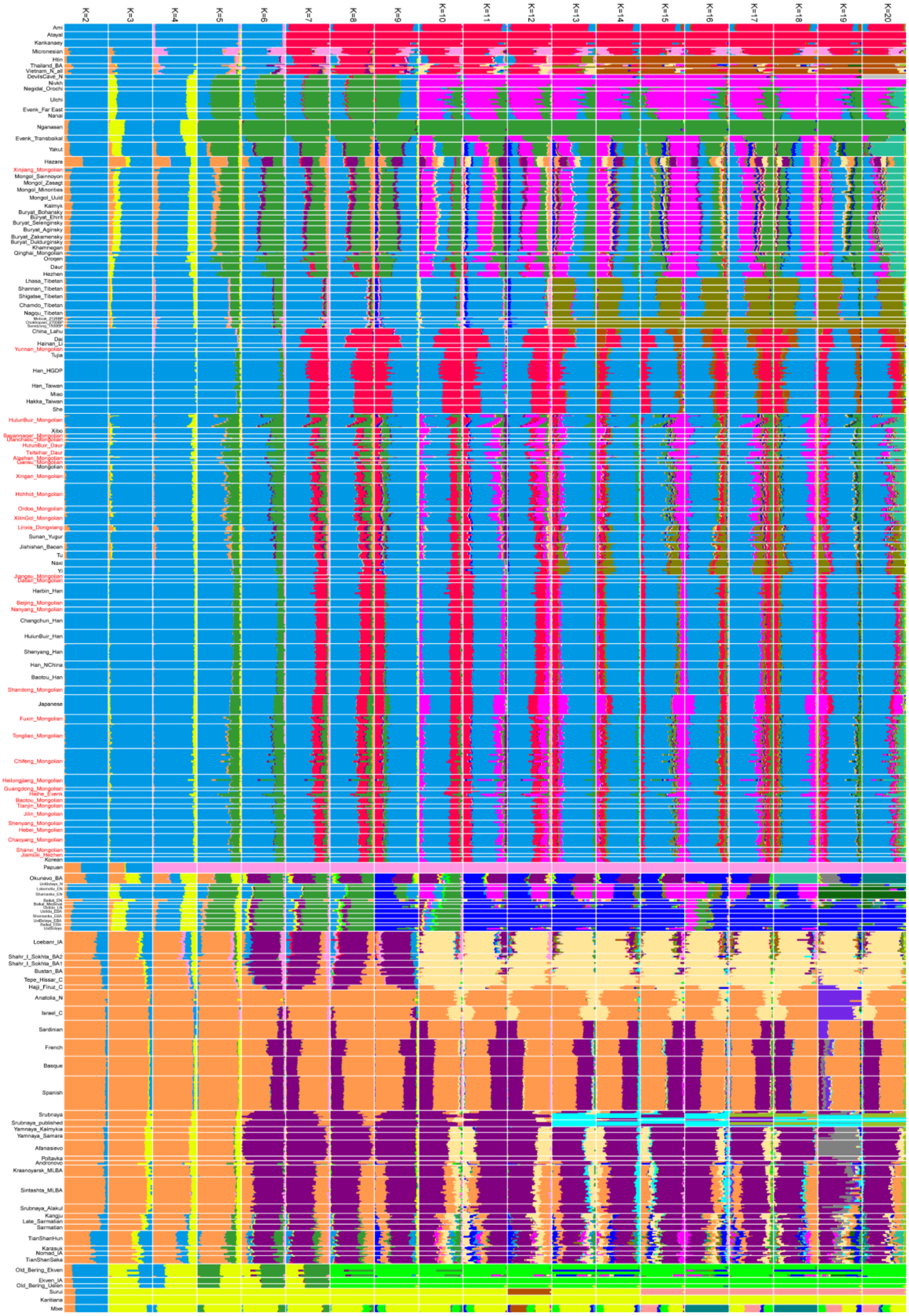


**Supplementary Fig. 6** Individual ancestry composition of newly studied populations and reference ancient and modern non-African populations revealed by ADMIXTURE analysis.

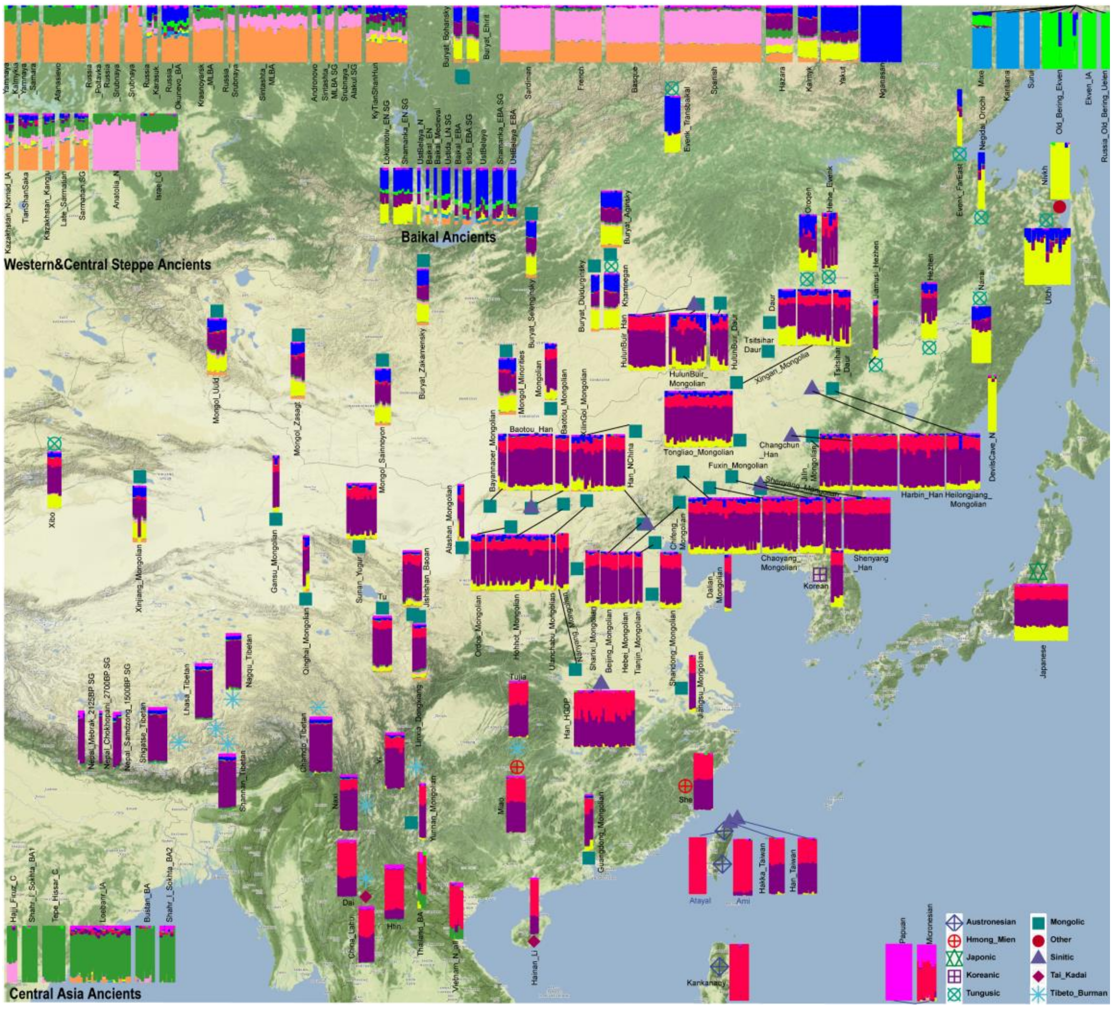


**Supplementary Fig. 7** The ancestry proportions of newly studied populations and reference ancient and modern non-African populations revealed by ADMIXTURE analysis with K = 7.

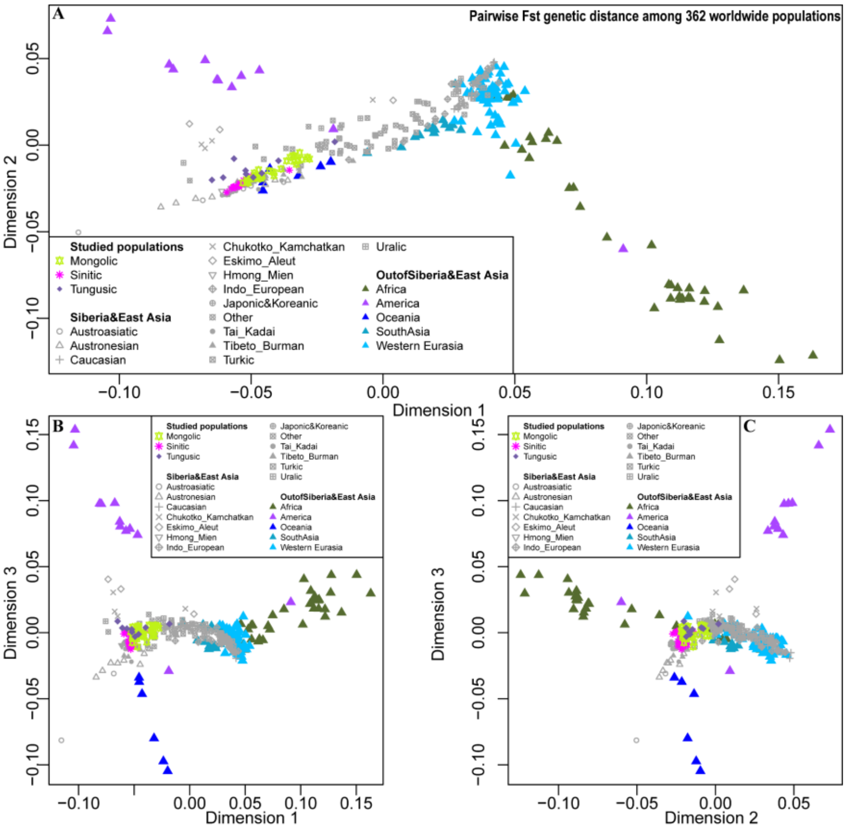


**Supplementary Fig. 8** Multidimensional scale analysis based on the Fst genetic distance matrix of 362 global populations.

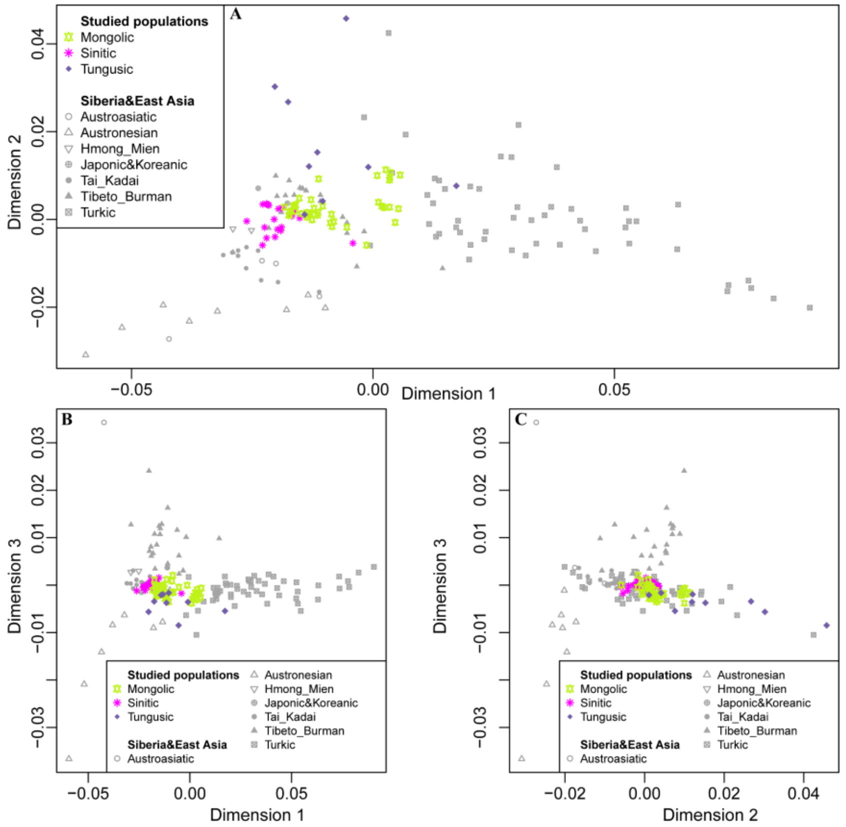


**Supplementary Fig. 9** Multidimensional scale analysis based on the Fst genetic distance matrix of East Asian populations.

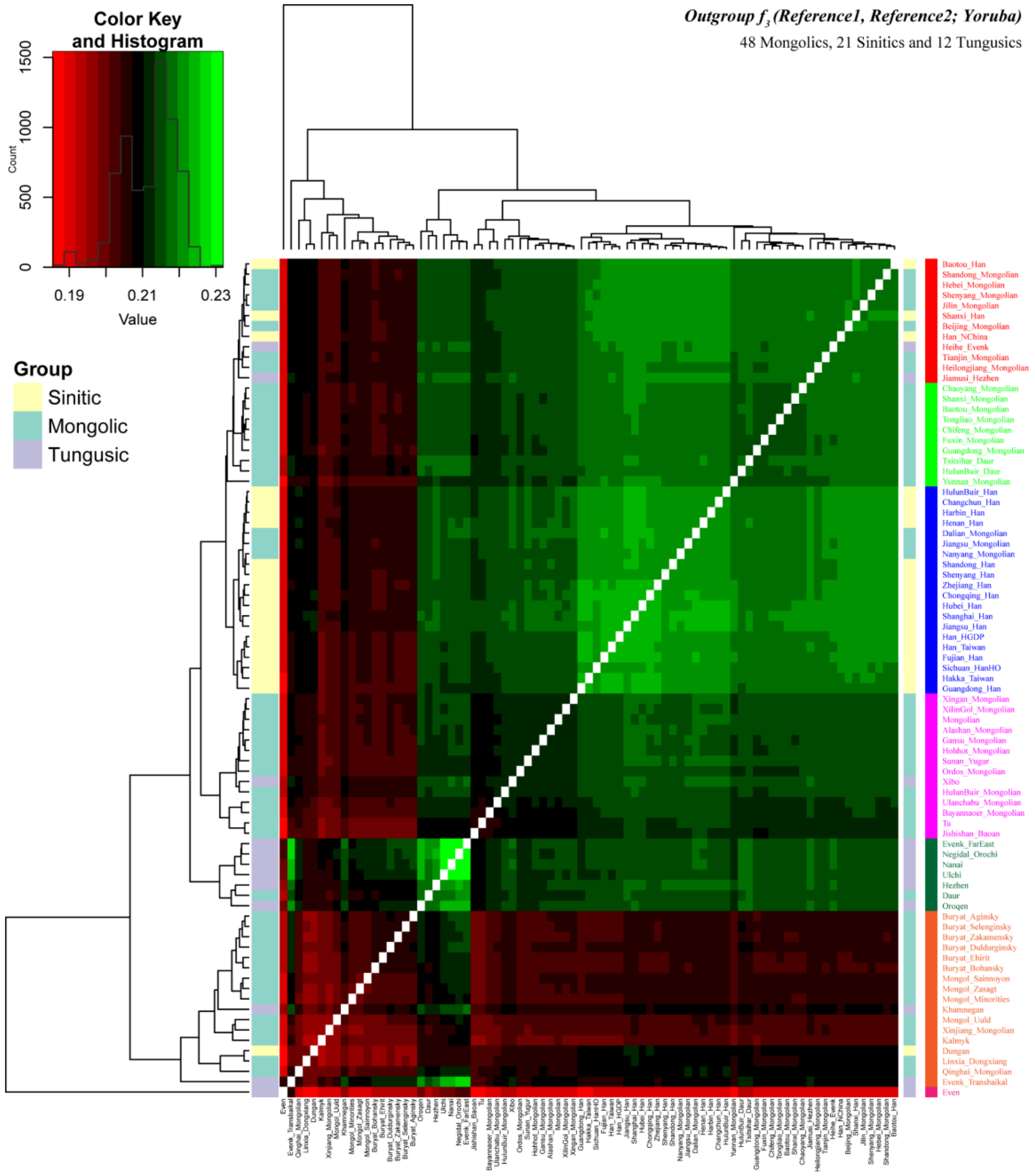


**Supplementary Fig. 10** The genetic affinity among Sinitic, Mongolic and Tungusic populations revealed by outgroup $f_3$-*statistics*.

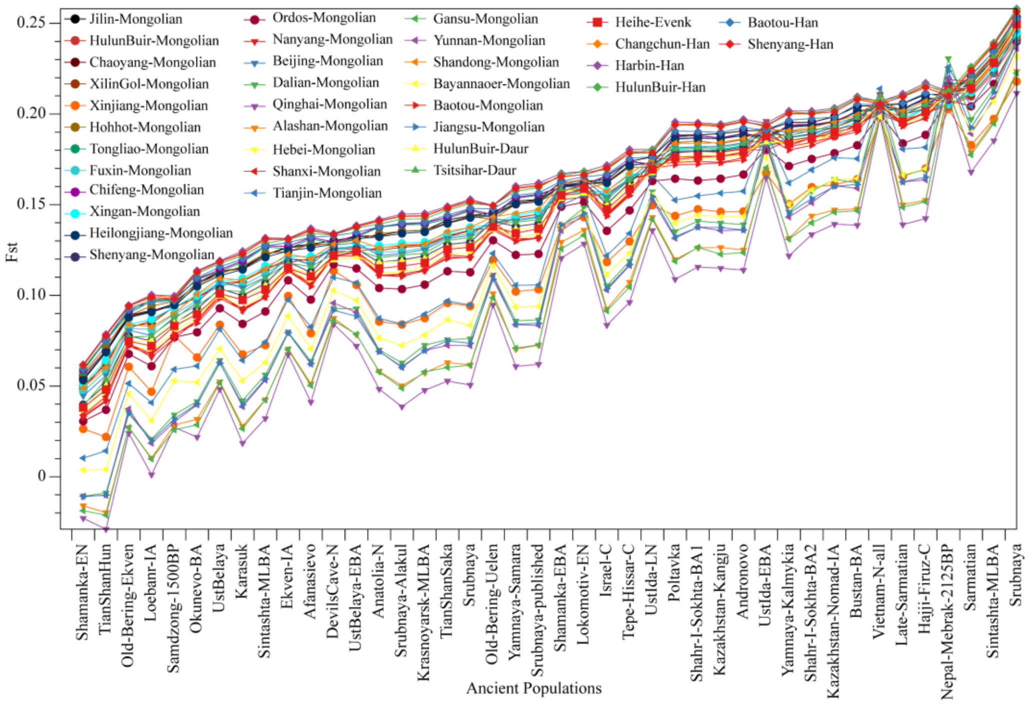


**Supplementary Fig. 11** The distributions of pairwise Fst genetic distances between newly studied population and ancient Eurasian populations.

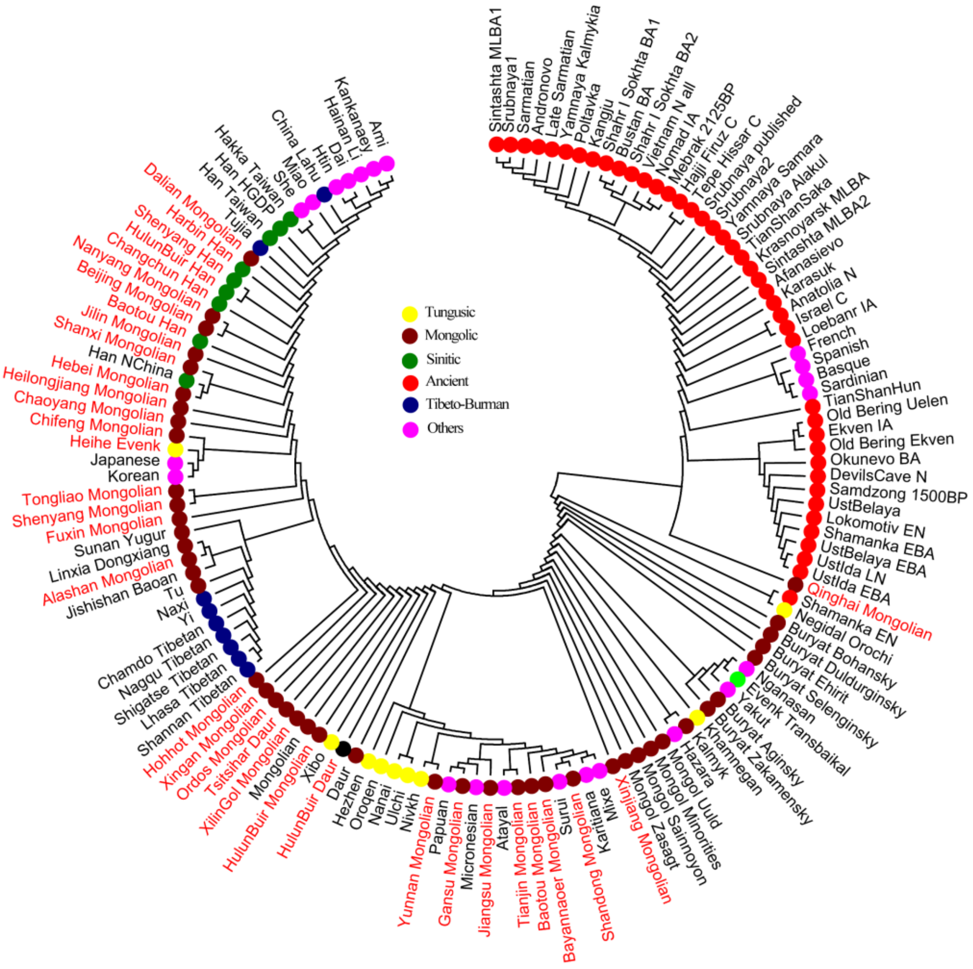


**Supplementary Fig. 12**The phylogenetic tree constructed based on pairwise Fst genetic distances among 138 modern and ancient populations.

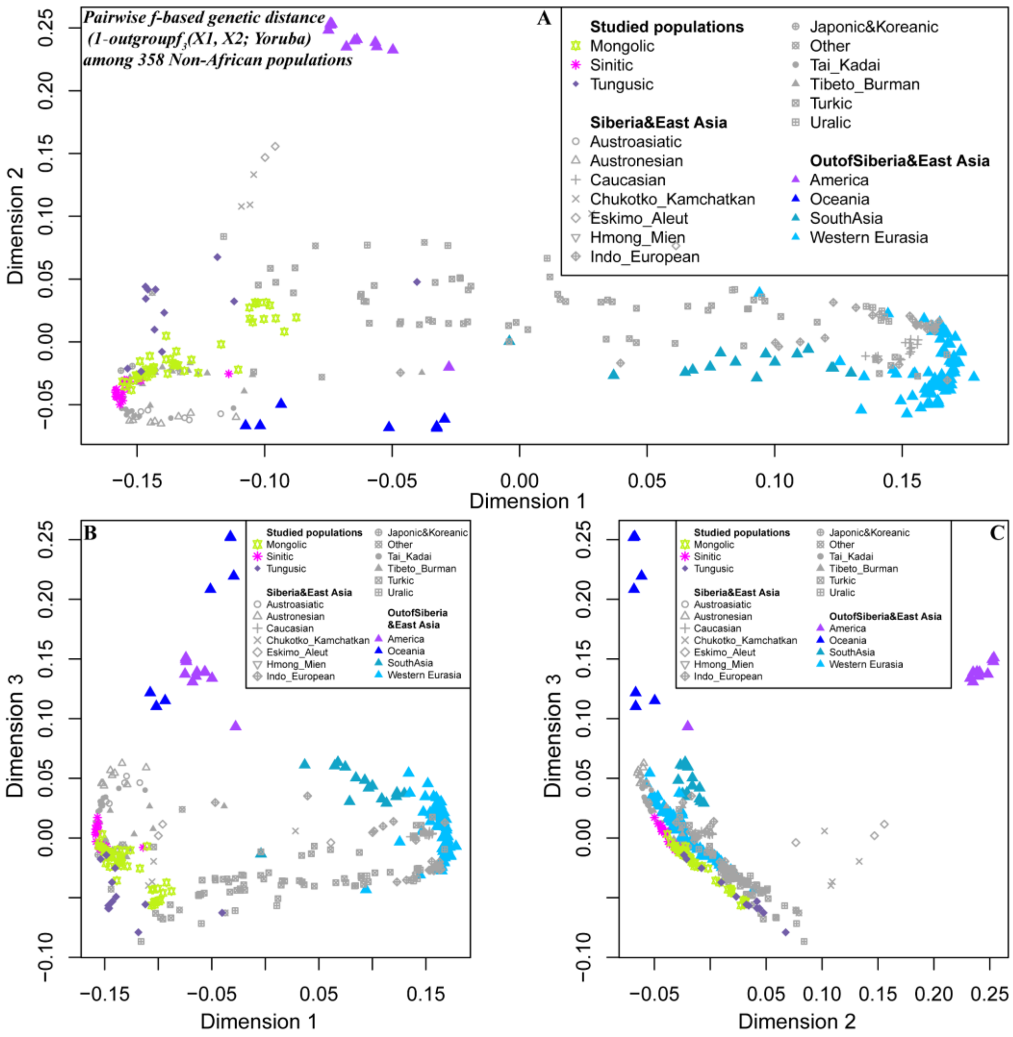


**Supplementary Fig. 13** Multidimensional scale analysis based on the 1 − $f_3$-value of 358 worldwide populations.

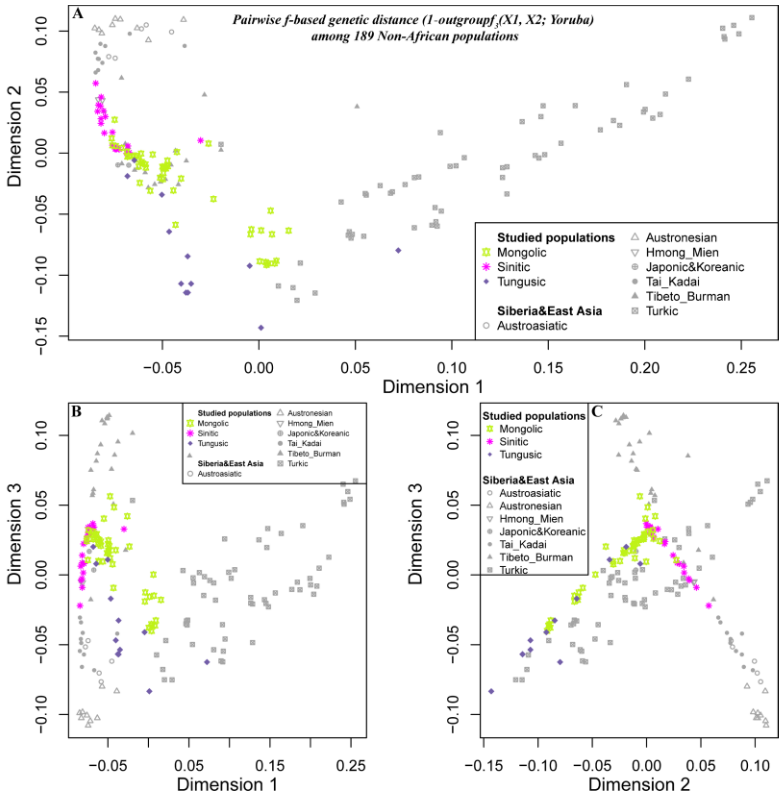

**Supplementary Fig. 14** Multidimensional scale analysis based on the1-$f_3$-value of 189 Eurasian populations.

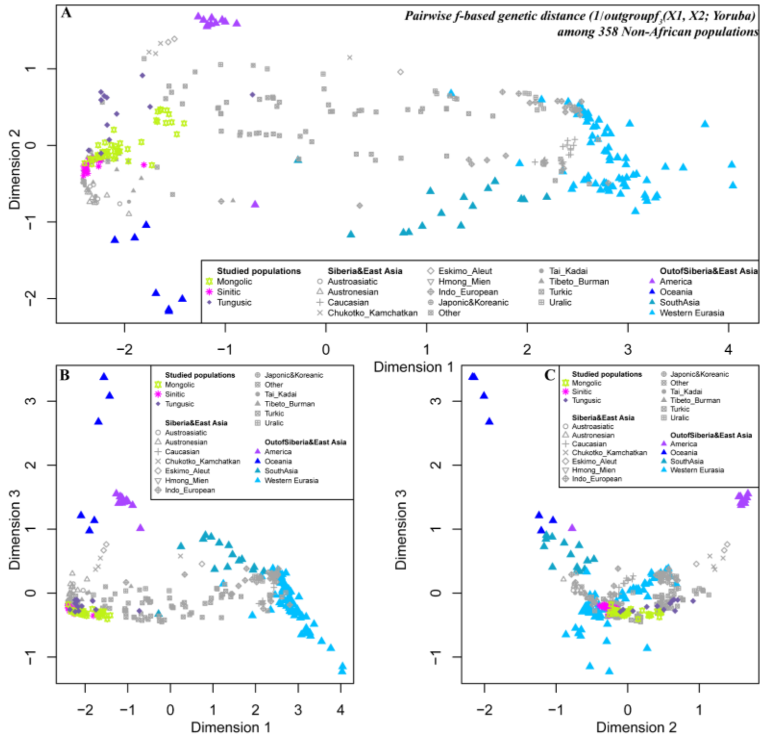

**Supplementary Fig. 15** Multidimensional scale analysis based on the 1/$f_3$-value of 358 worldwide populations.

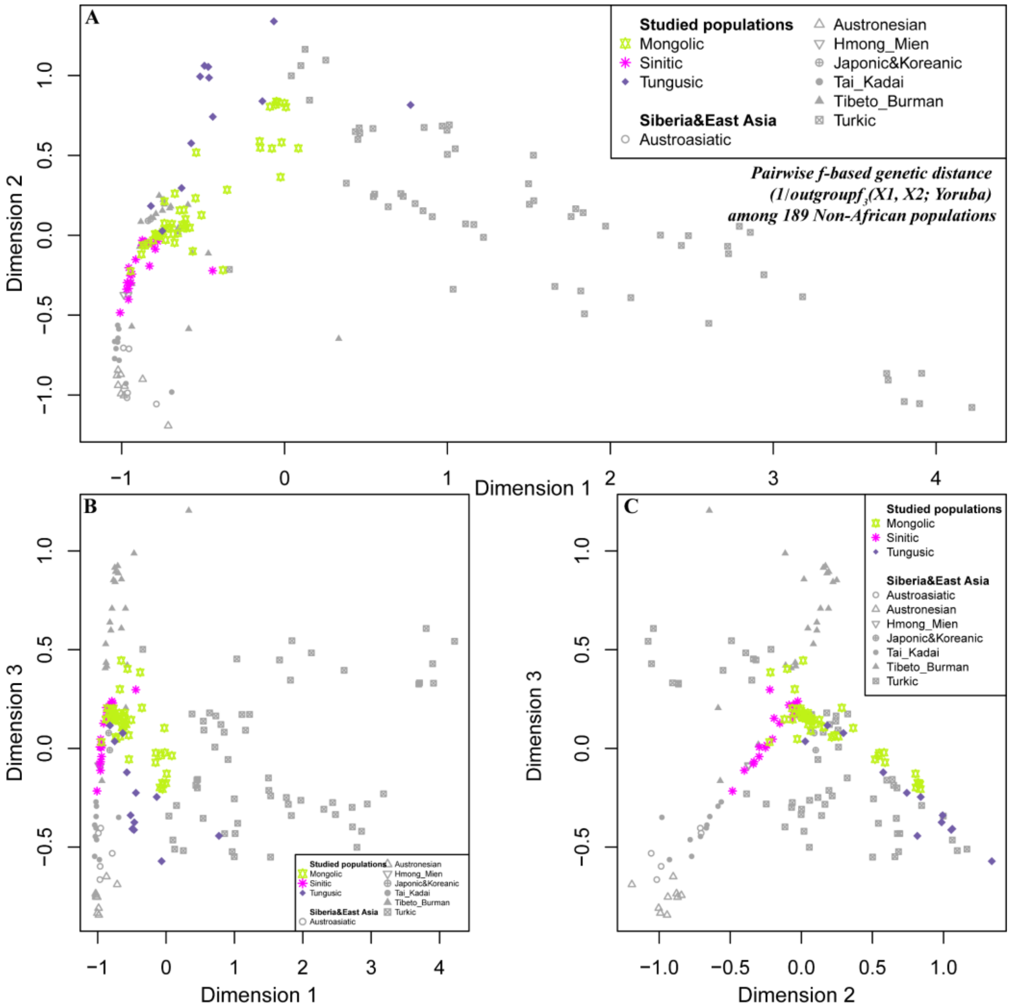


**Supplementary Fig. 16** Multidimensional scale analysis based on the 1/$f_3$-value of 189 Eurasian populations.

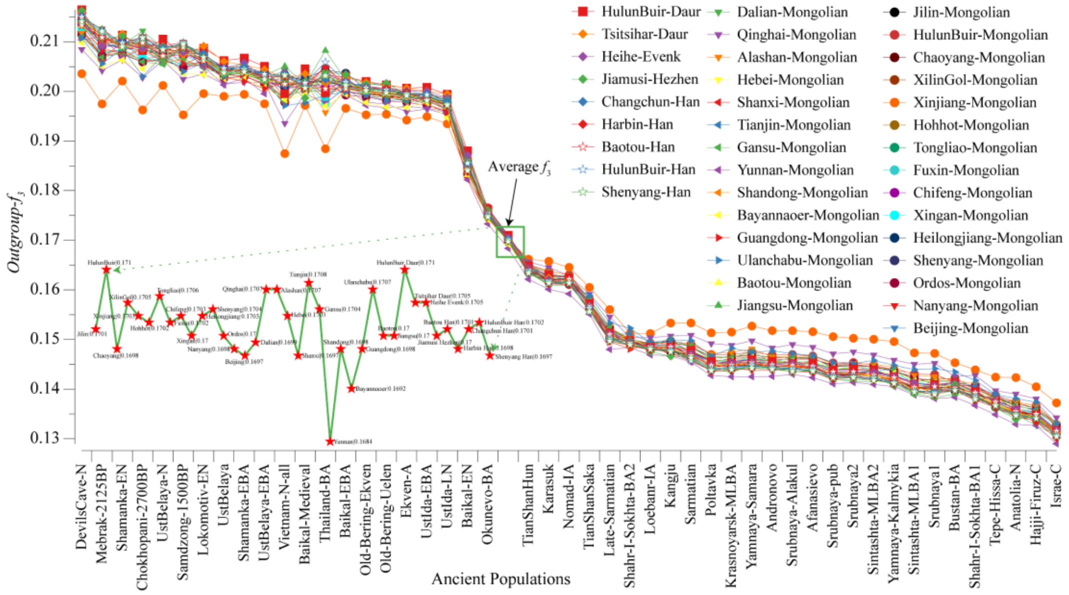


**Supplementary Fig. 17** The distributions of outgroup $f_3$-value between 38 newly studied population and ancient Eurasian populations.

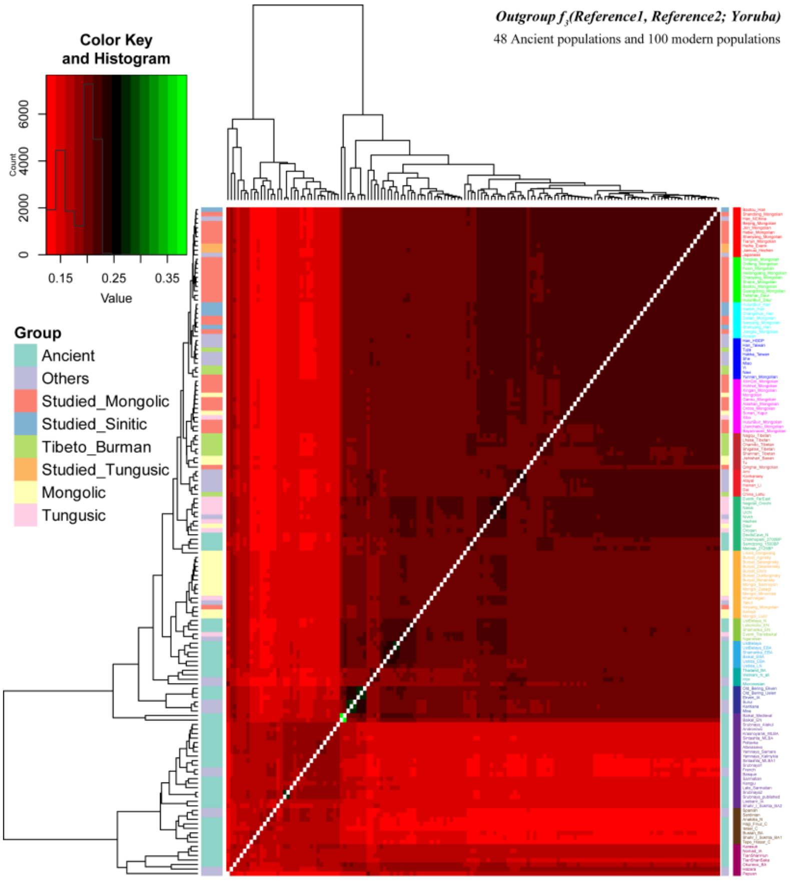


**Supplementary Fig. 18** Outgroup $f_3$-value between 38 newly studied population and ancient Eurasian populations.

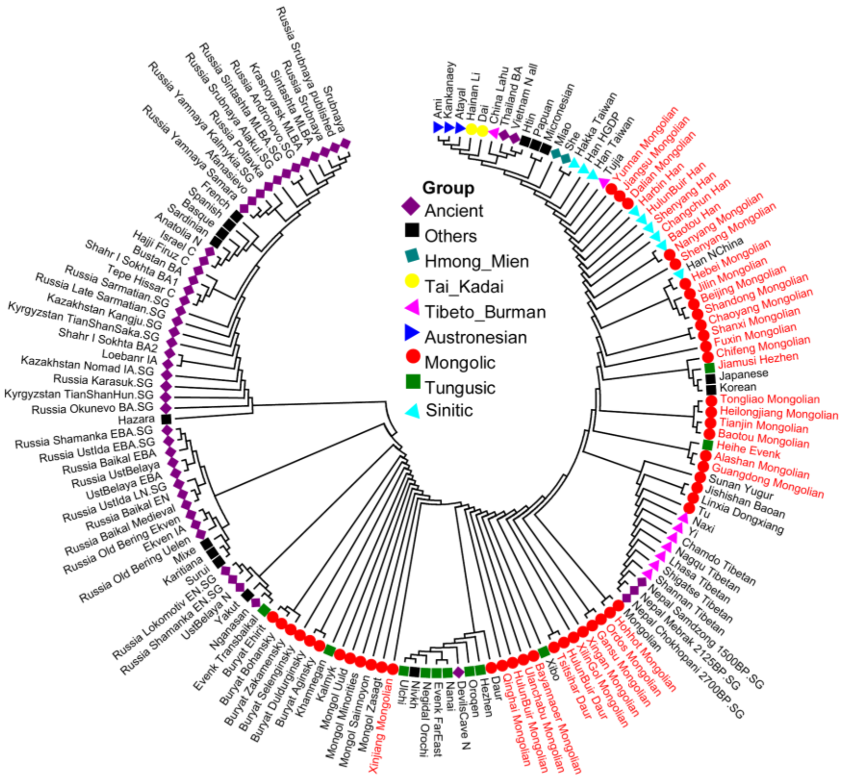


**Supplementary Fig. 19** The phylogenetic relationship among modern and ancient Eurasian populations.

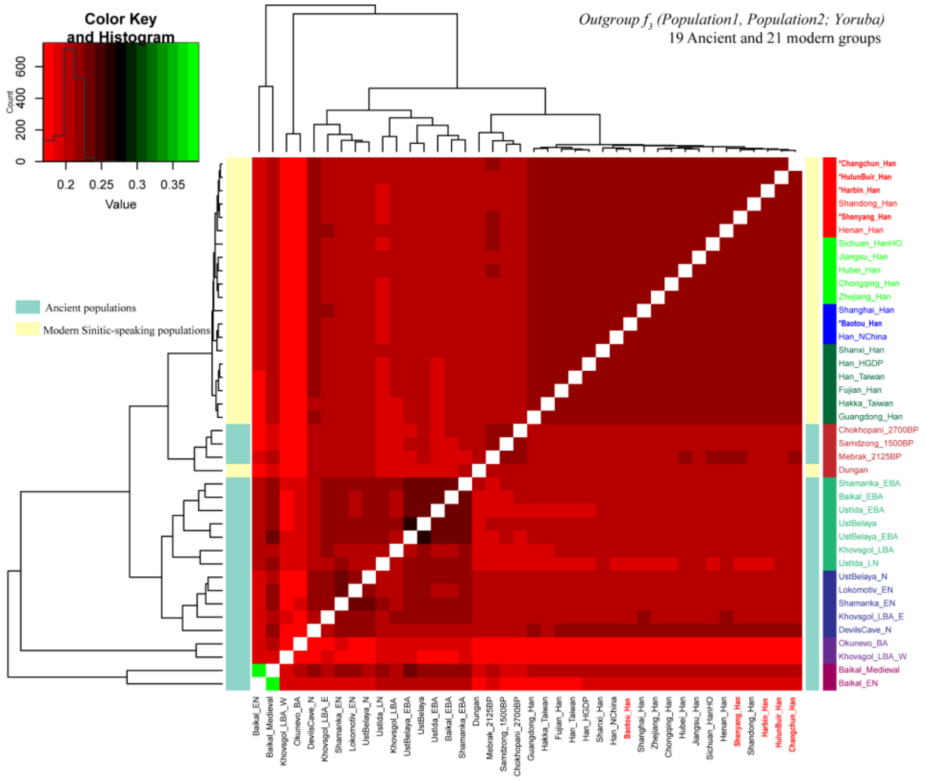


**Supplementary Fig. 20** The genetic relationship between Sinitic-speaking populations and modern and ancient East Eurasian populations revealed by outgroup $f_3$-statistics.

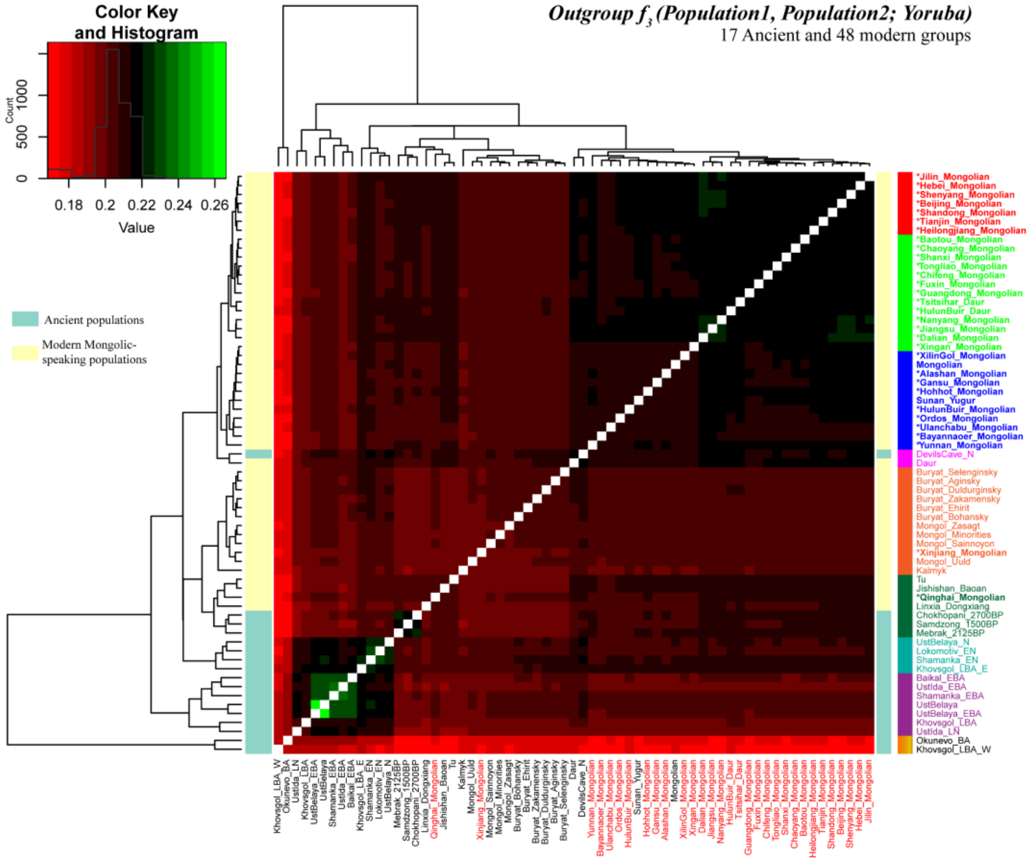


**Supplementary Fig. 21** The genetic relationship between Mongolic-speaking populations and modern and ancient East Eurasian populations revealed by outgroup $f_3$-statistics.

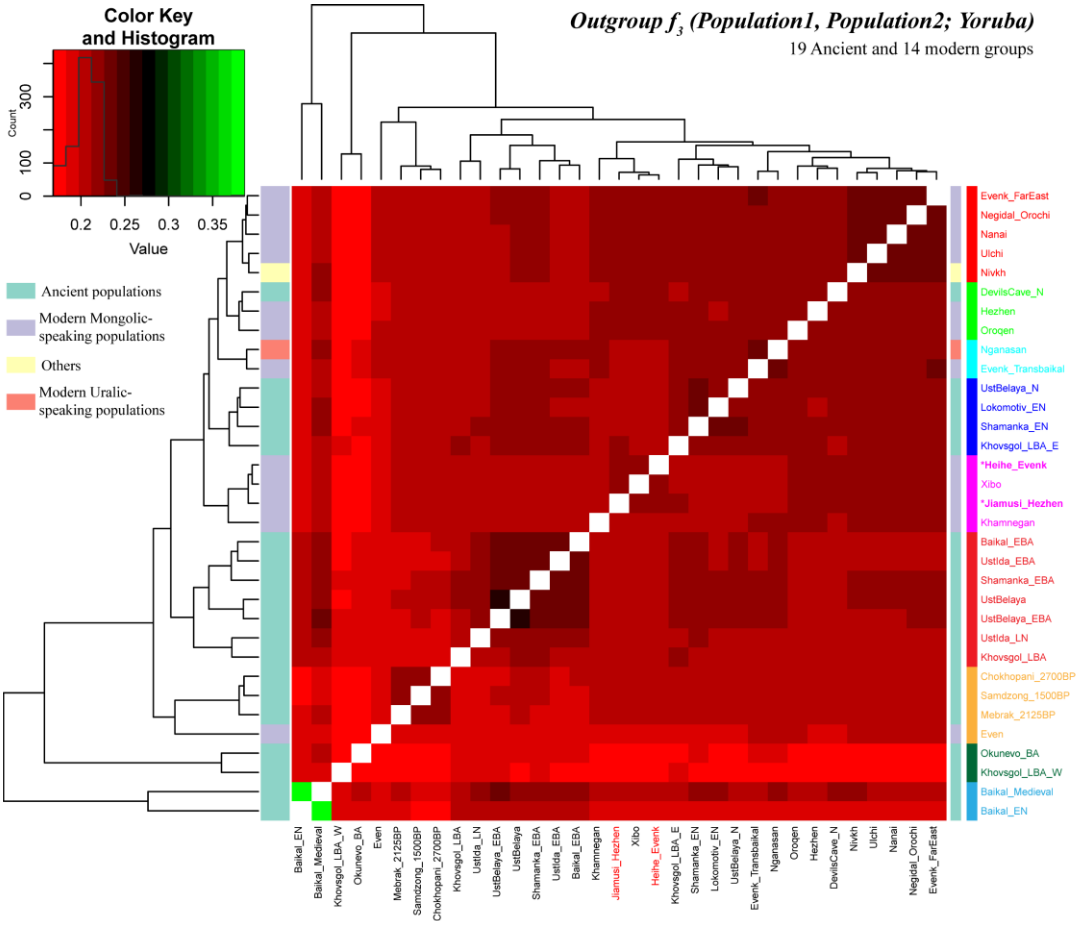


**Supplementary Fig. 22** The genetic relationship between Tungusic-speaking populations and modern and ancient East Eurasian populations revealed by outgroup $f_3$-statistics.

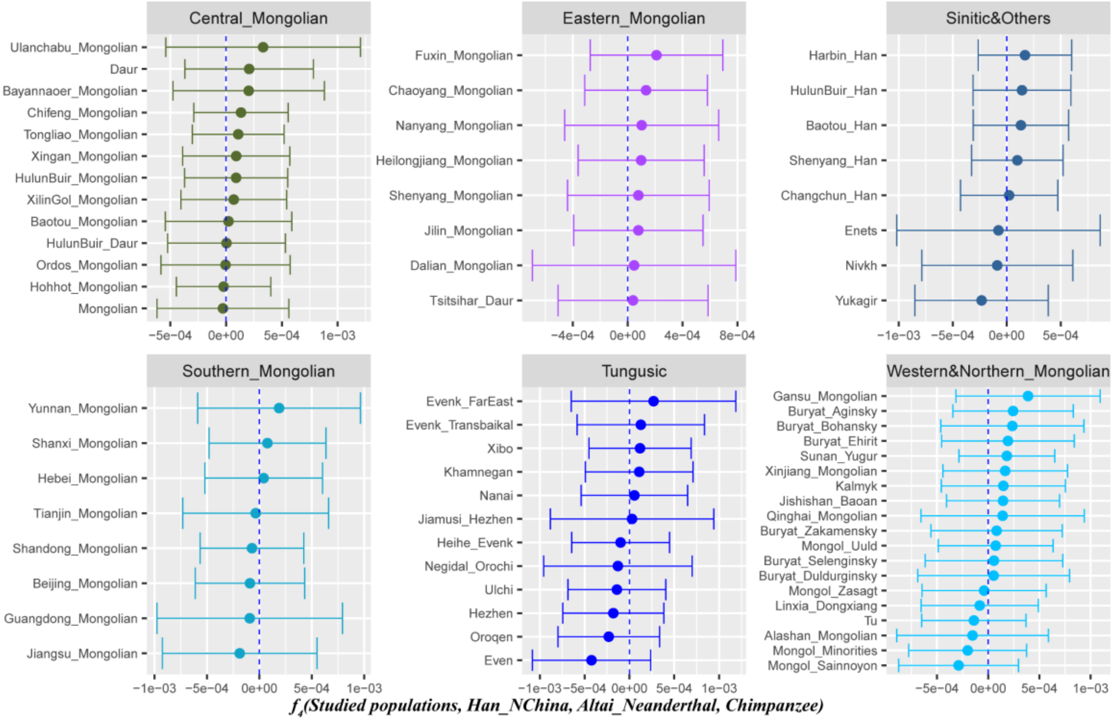


**Supplementary Fig. 23** The gene flow of Altai_Neanderthal into studied populations revealed by $f_4$-*statistics* in the form of ***$f_4$(Studied populations, Han_NChina; Altai_Neanderthal, Chimpanzee)***.

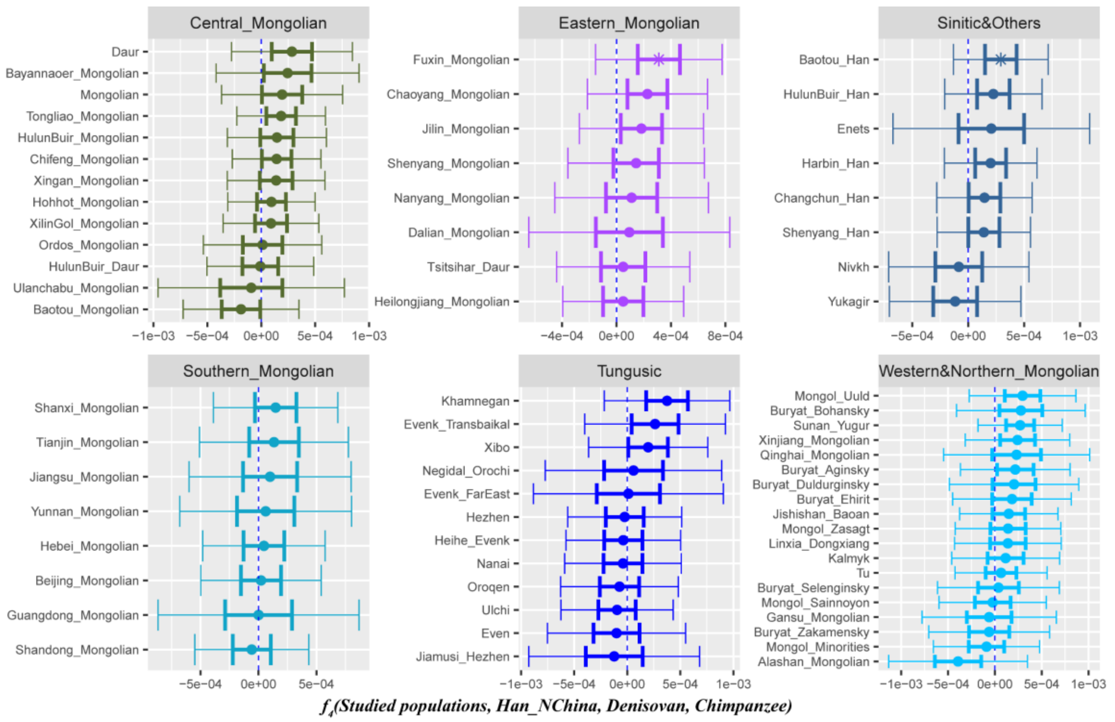


**Supplementary Fig. 24** The gene flow of Altai_Denisovan into studied populations revealed by *$f_4$-statistics* in the form of ***$f_4$(Studied populations, Han_NChina; Altai_Denisovan, Chimpanzee)***.

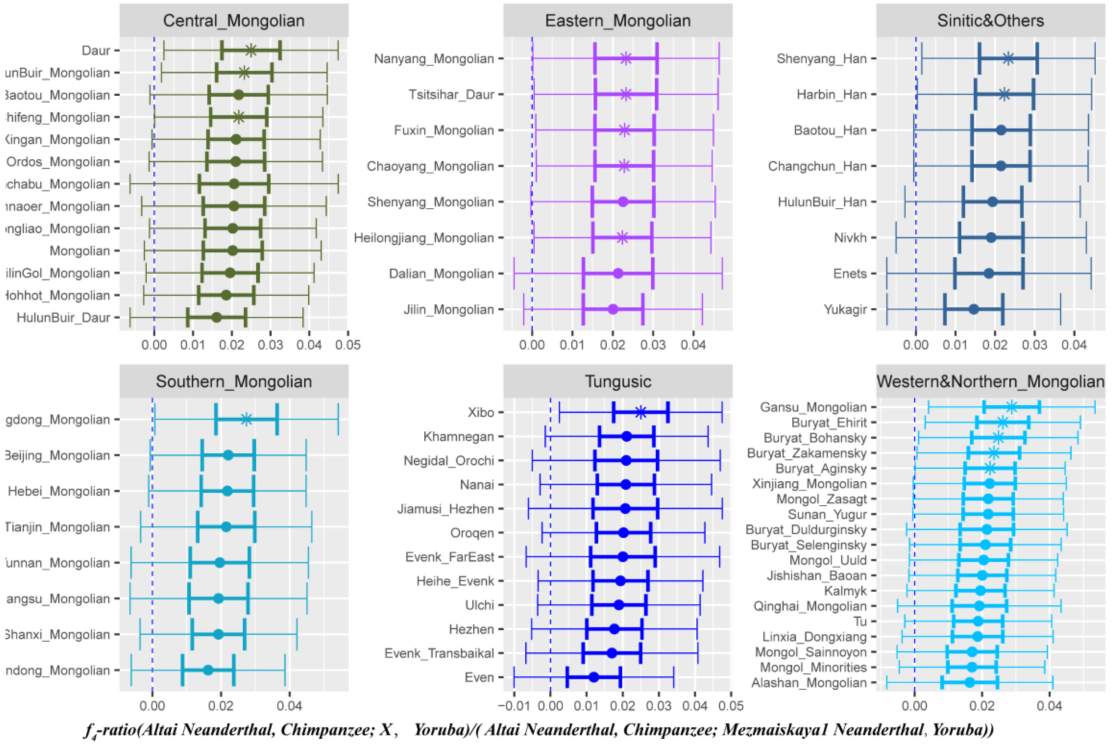


**Supplementary Fig. 25** The gene flow of Altai_Neanderthal into studied populations revealed by *$f_4$-ratio* in the form of ***$f_4$(Altai_Neanderthal, Chimpanzee; X, Yoruba)/$f_4$(Altai_Neanderthal, Chimpanzee; Mezmaiskayal_Neanderthal, Yoruba)***.

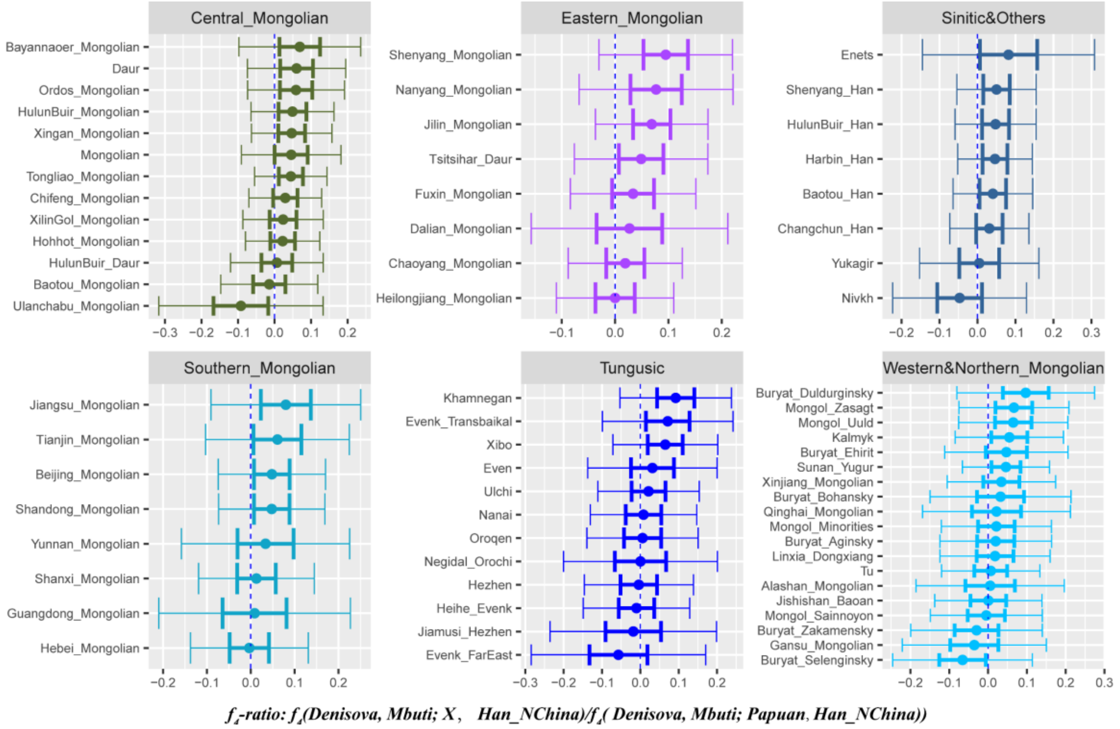


**Supplementary Fig. 26** The gene flow of Altai_Denisovan into studied populations revealed by *$f_4$-ratio* in the form of ***$f_4$* (Altai_Denisovan, Mbuti; X, Han_NChina)/*$f_4$* (Altai_Denisovan, Mbuti; Papuan, Han_NChina)**.

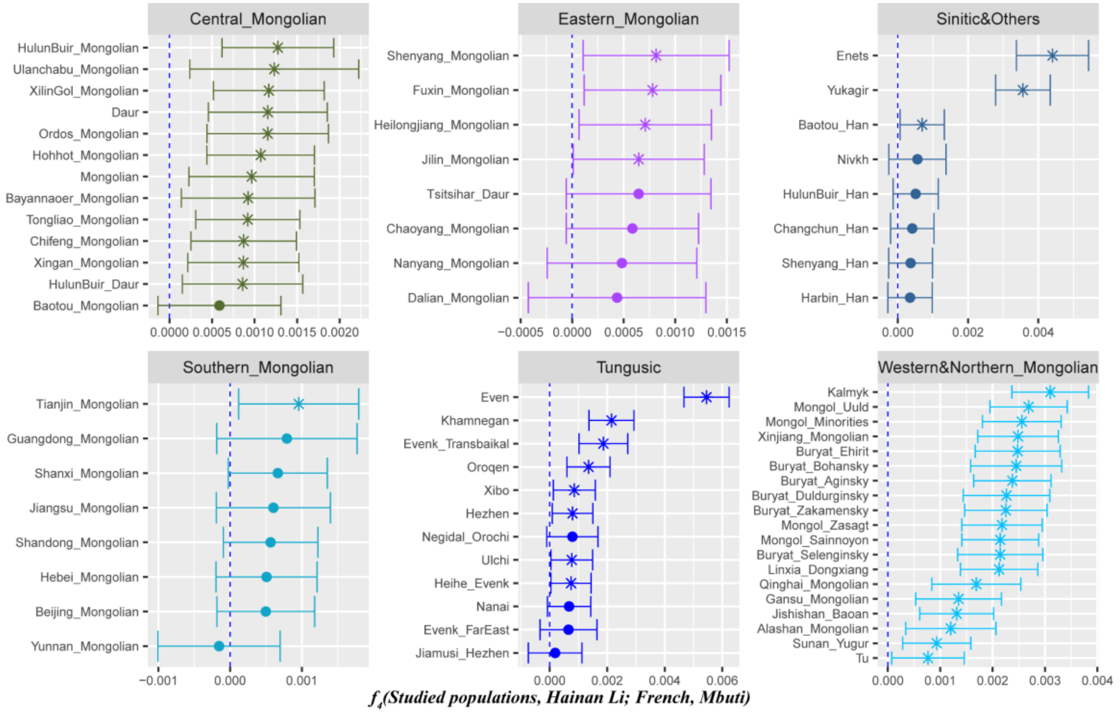


**Supplementary Fig. 27** The gene flow of West Eurasian populations (French) into studied populations revealed by *$f_4$-statistics* in the form of ***$f_4$*(Studied populations, Hainan Li; French, Mbuti)**, Hainan Li is used as the East Eurasian source population.

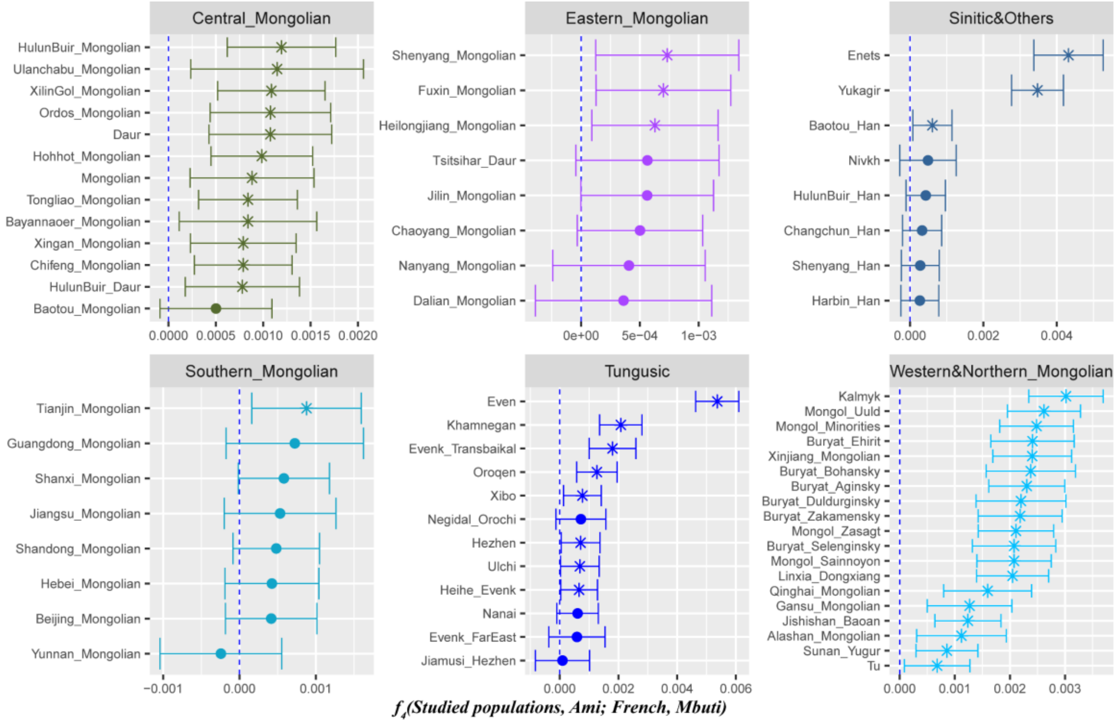


**Supplementary Fig. 28** The gene flow of West Eurasian populations (French) into studied populations revealed by *$f_4$-statistics* in the form of ***$f_4$(Studied populations, Hainan Li; French, Mbuti)***, Ami is used as the East Eurasian source population.

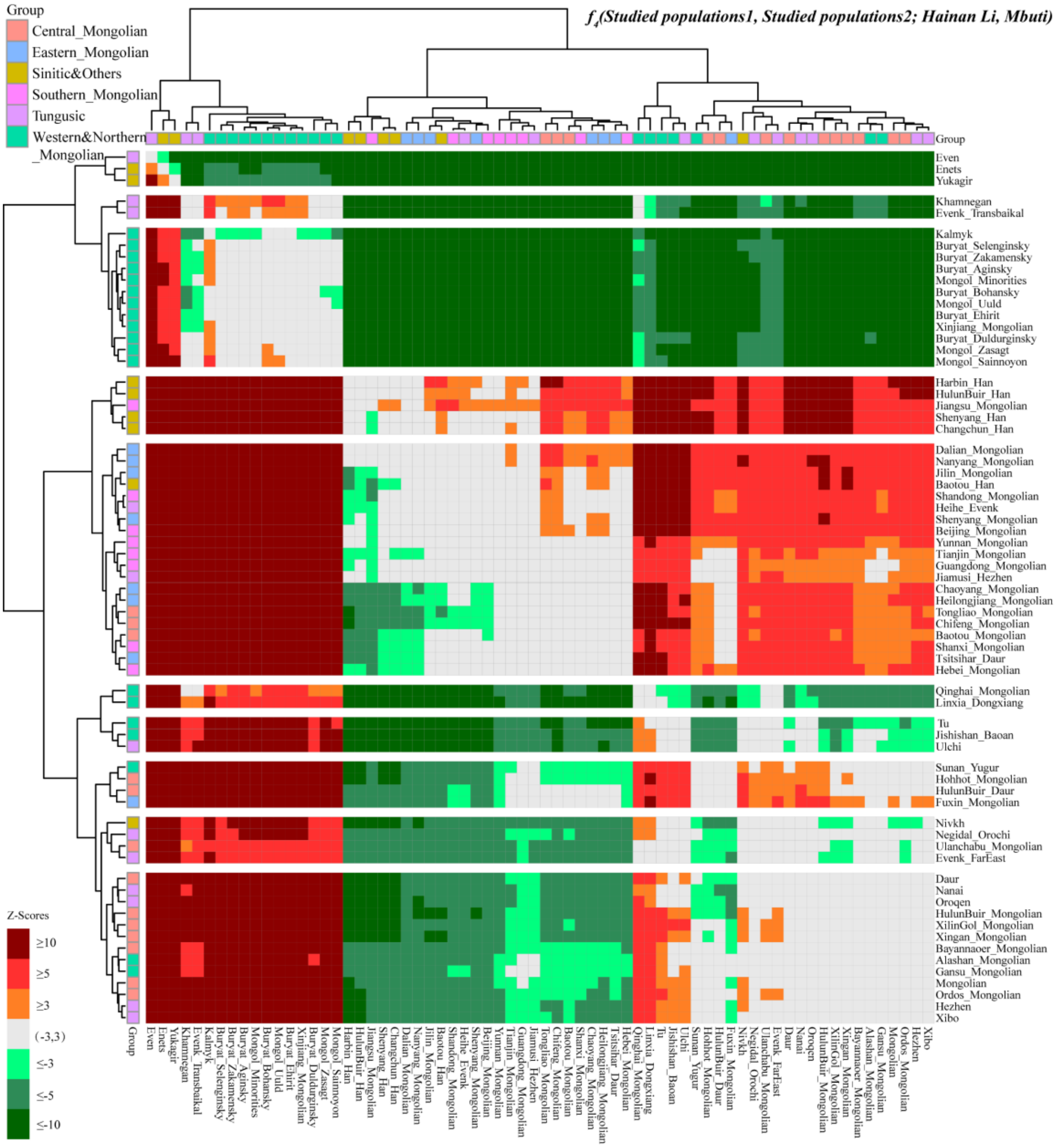


**Supplementary Fig. 29** Population substructures among studied populations revealed by *$f_4$-statistics* in the form of ***$f_4$(Studied populations1, Studied populations2; Hainan Li, Mbuti)***, Hainan Li is used as the reference baseline.

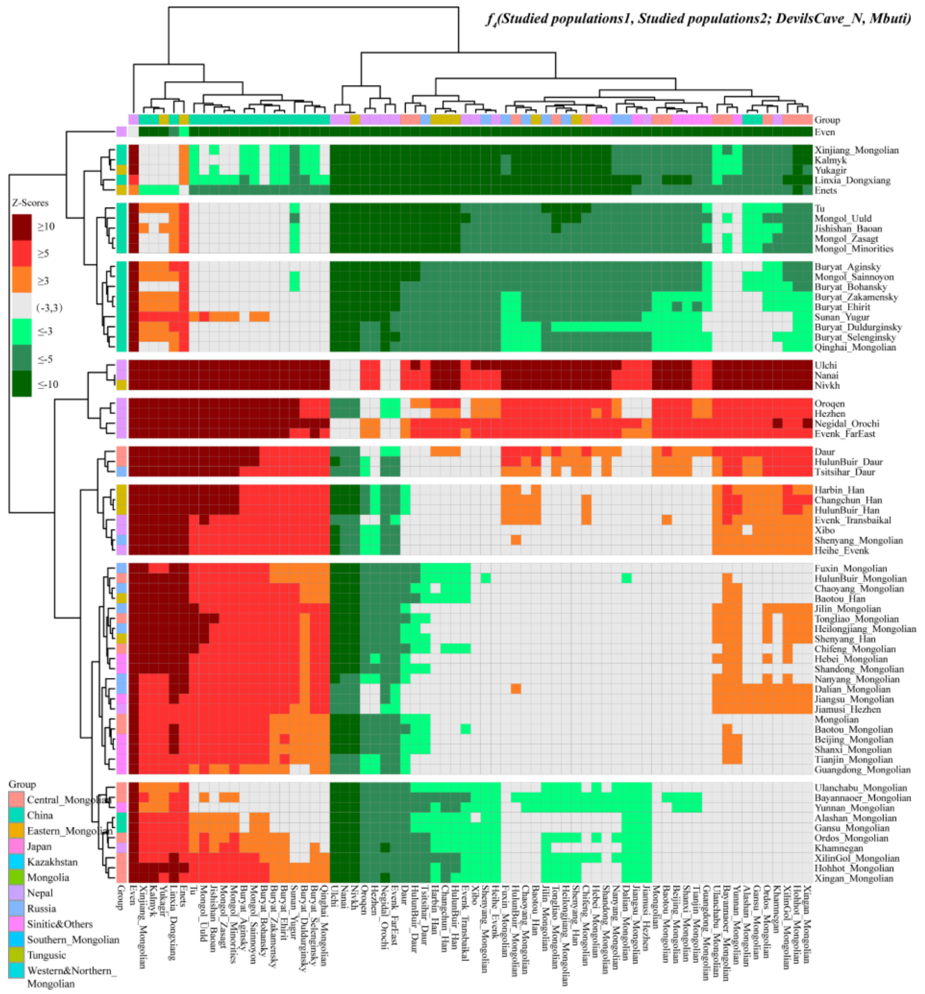


**Supplementary Fig. 30** Population substructures among studied populations revealed by *$f_4$-statistics* in the form of ***$f_4$(Studied populations1, Studied populations2; DevilsCave_N, Mbuti)***, DevilsCave_N is used as the reference baseline.

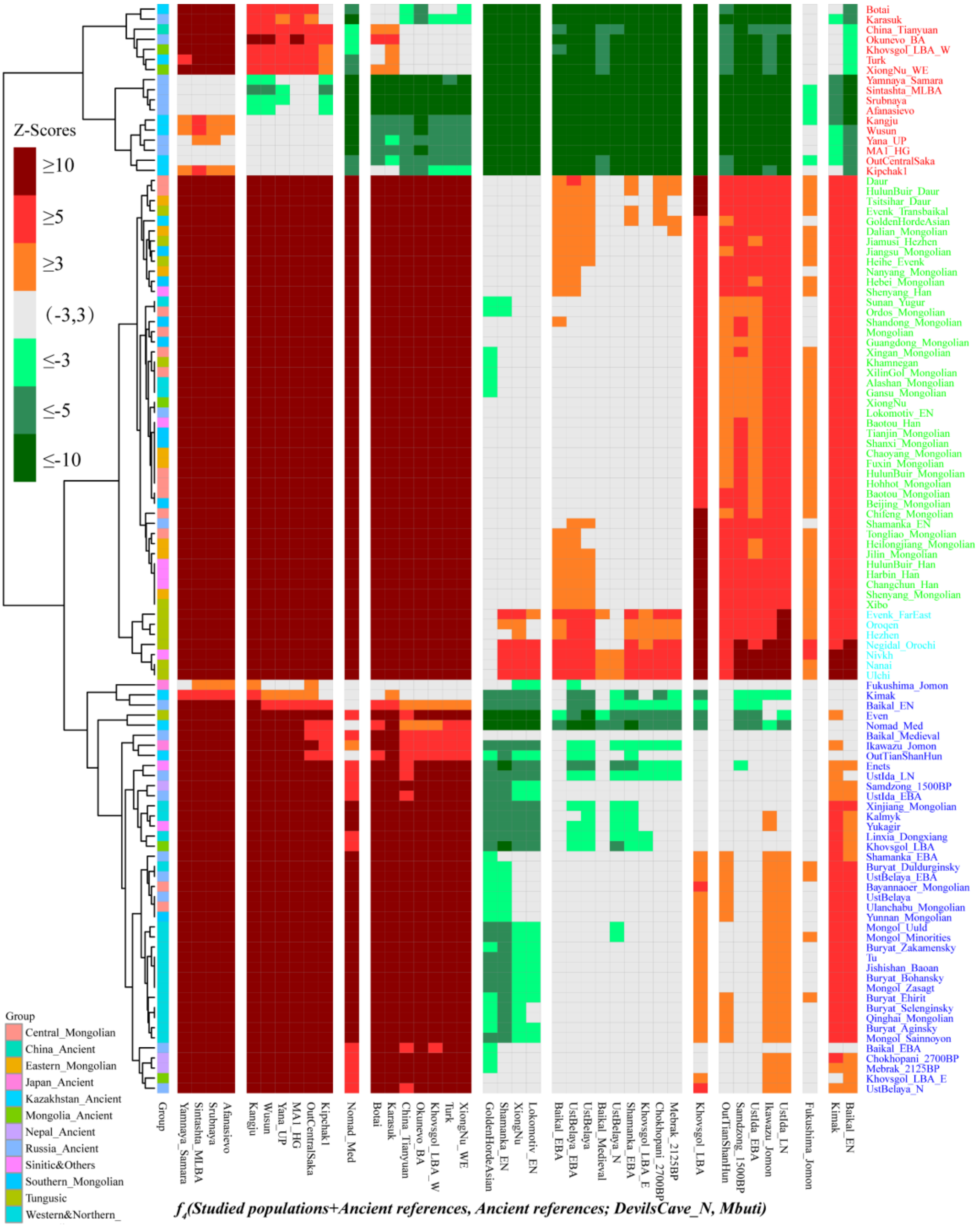

**Supplementary Fig. 31** Population substructures among studied populations revealed by *$f_4$-statistics* in the form of ***$f_4$(Studied populations + Ancient references, Ancient references; DevilsCave_N, Mbuti)***, DevilsCave_N is used as the reference baseline.

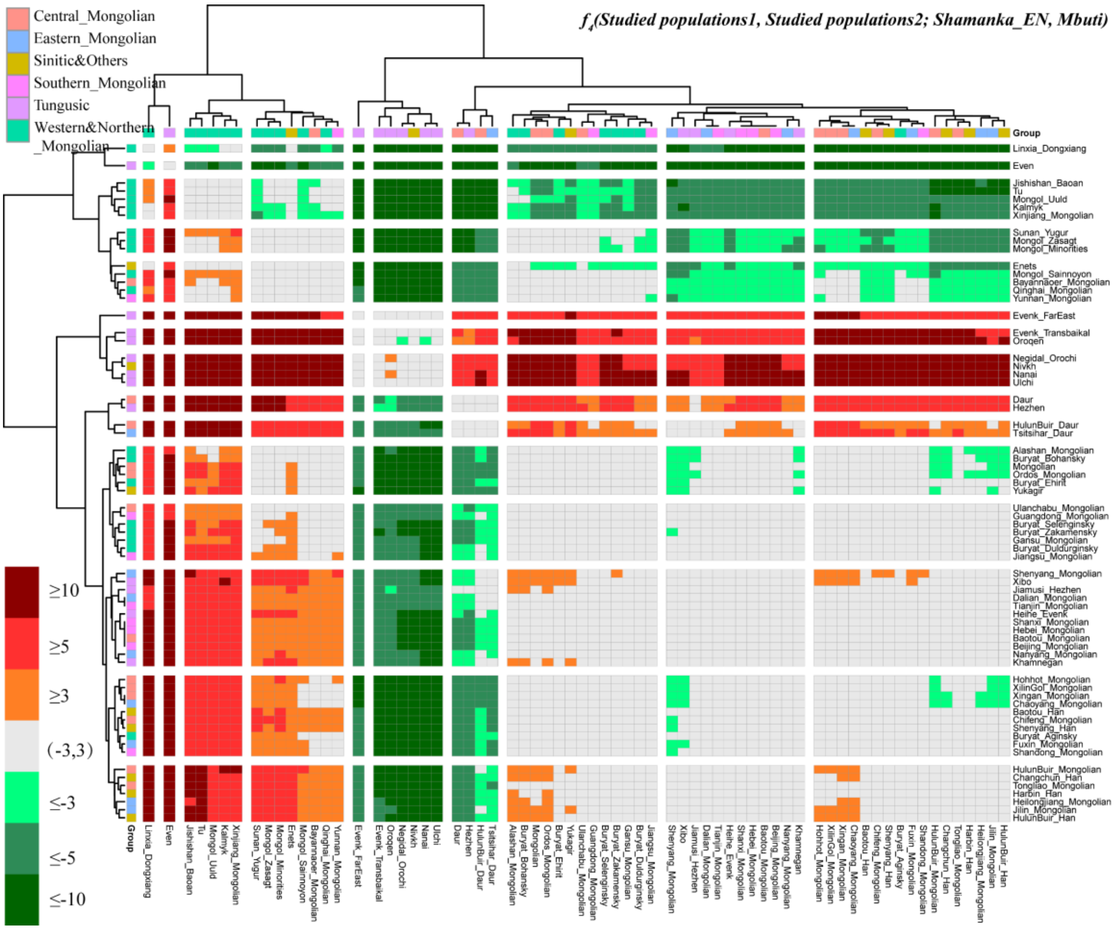


**Supplementary Fig. 32** Population substructures among studied populations revealed by $f_4$*-statistics* in the form of ***$f_4$(Studied populations1, Studied populations2; Shamanka_EN, Mbuti)***, Shamanka_EN is used as the reference baseline.

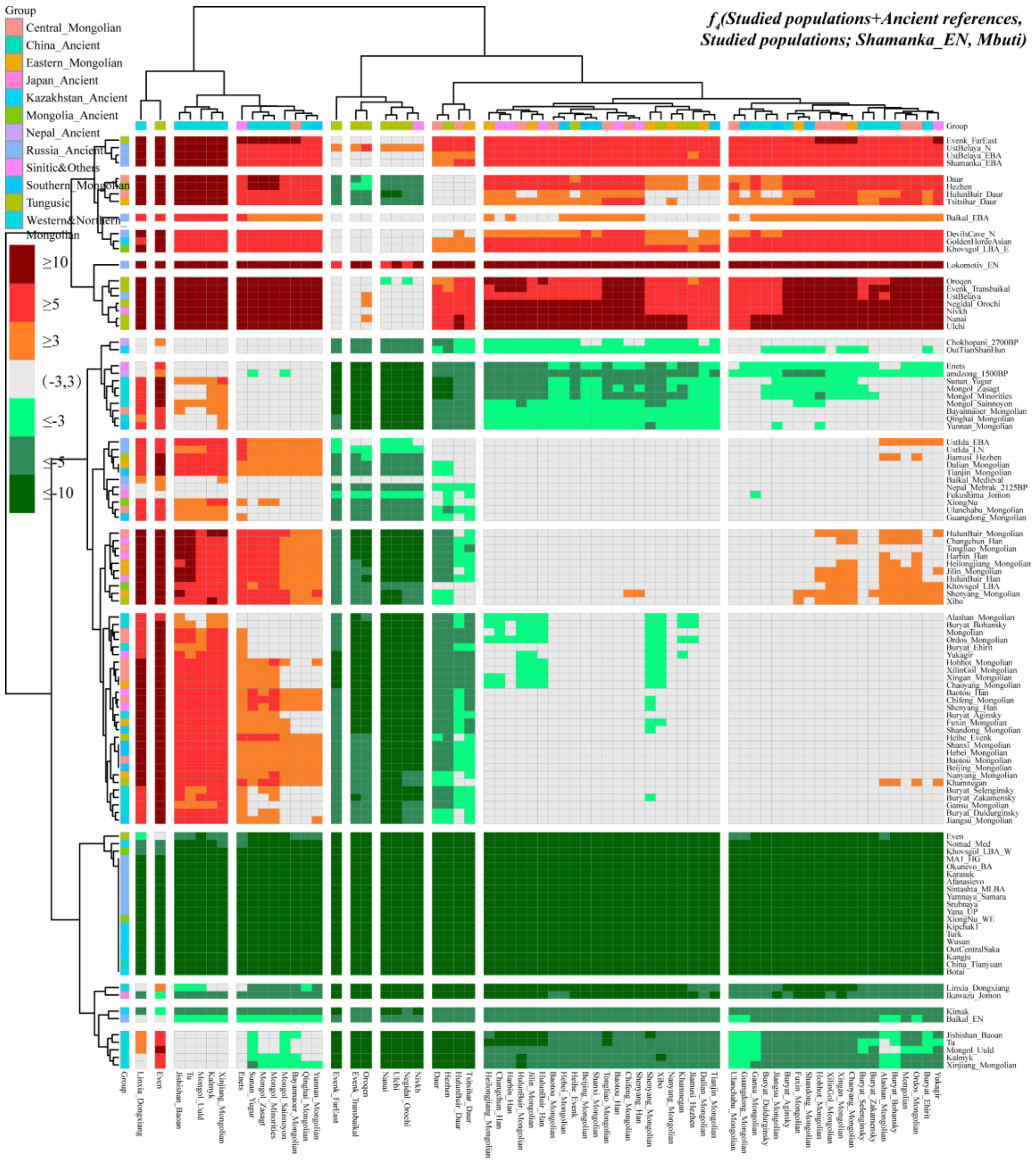


**Supplementary Fig. 33** Population substructures among studied populations revealed by *$f_4$-statistics* in the form of ***$f_4$(Studied populations + Ancient references, Ancient references; Shamanka_EN, Mbuti)***, Shamanka_EN is used as the reference baseline.

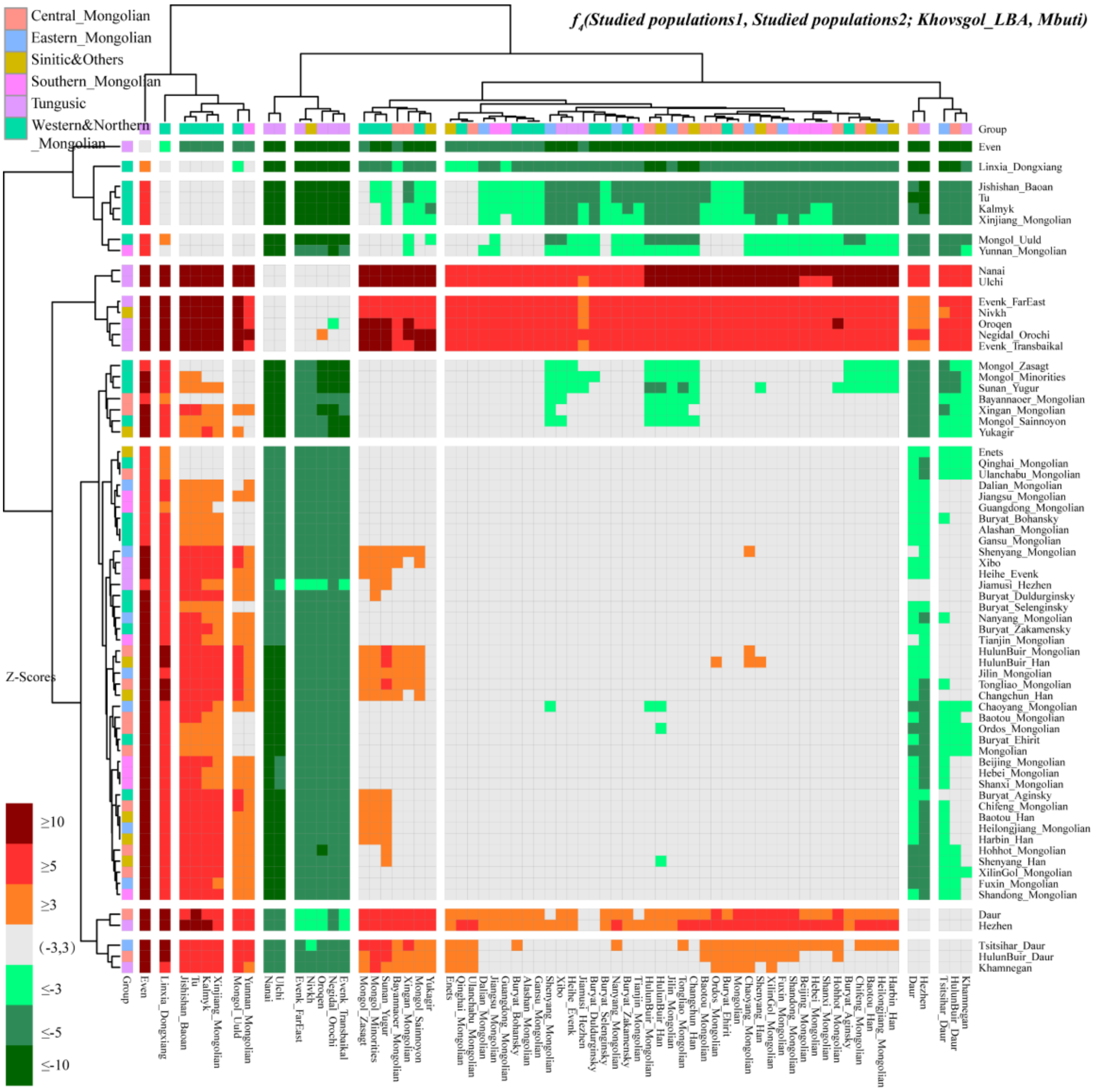


**Supplementary Fig. 34** Population substructures among studied populations revealed by *$f_4$-statistics* in the form of ***$f_4$(Studied populations1, Studied populations2; Khovsgol_LBA, Mbuti)***, Khovsgol_LBA is used as the reference baseline.

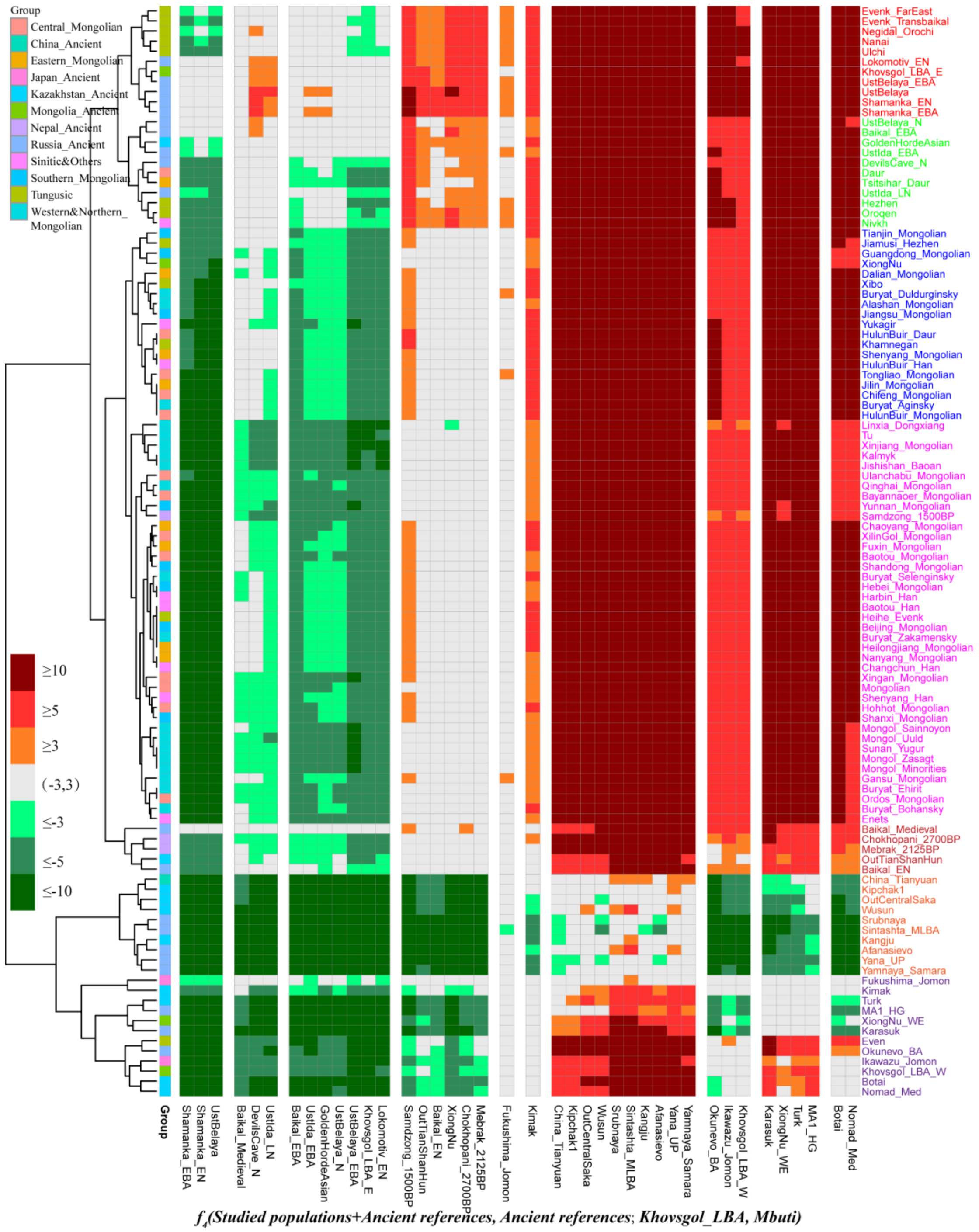


**Supplementary Fig. 35** Population substructures among studied populations revealed by *$f_4$-statistics* in the form of ***$f_4$(Studied populations + Ancient references, Ancient references; Khovsgol_LBA, Mbuti)***, Khovsgol_LBA is used as the reference baseline.

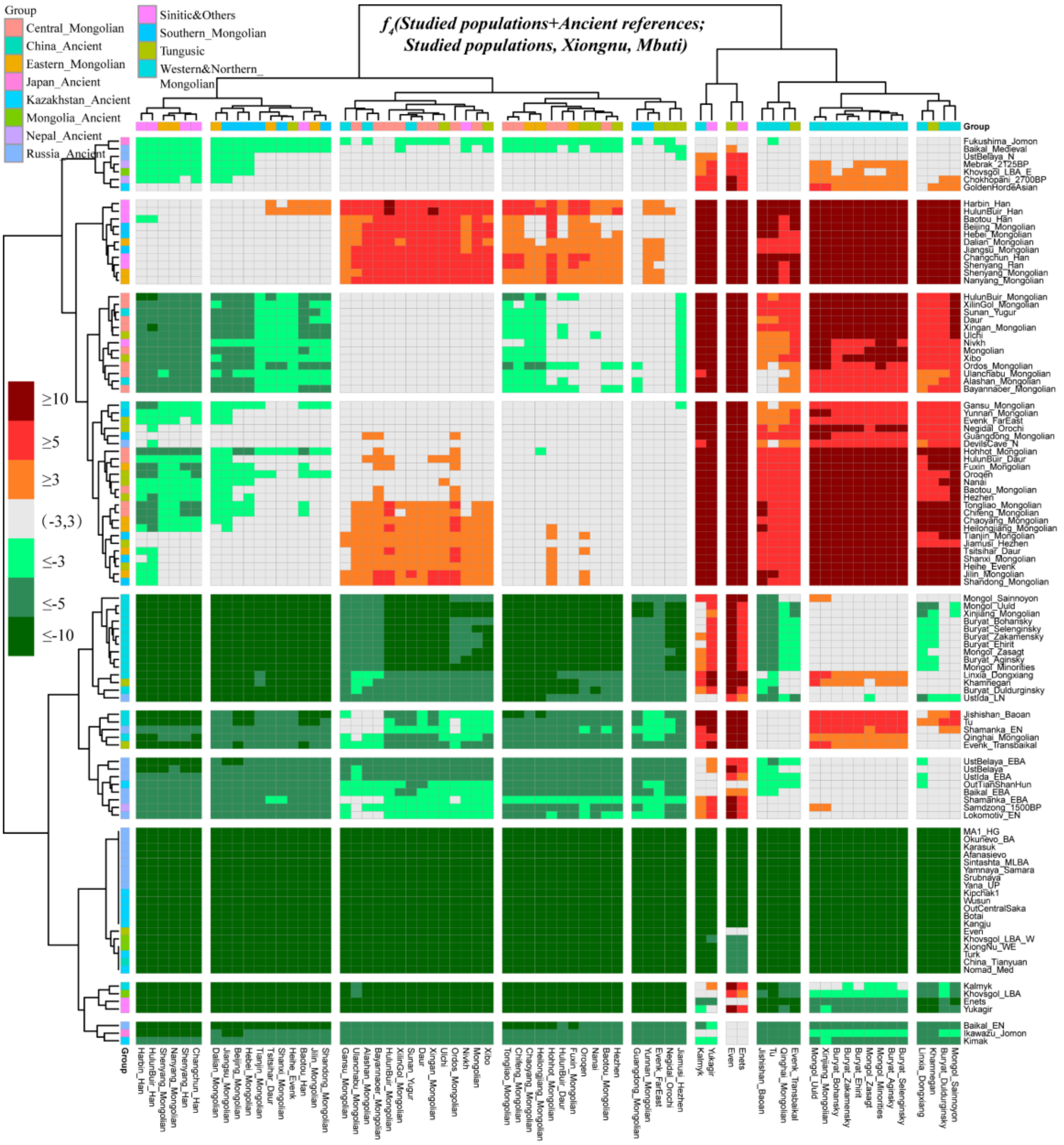


**Supplementary Fig. 36** Population substructures among studied populations revealed by *$f_4$-statistics* in the form of ***$f_4$(Studied populations + Ancient references, Studied populations; XiongNu, Mbuti)***, XiongNu is used as the reference baseline.

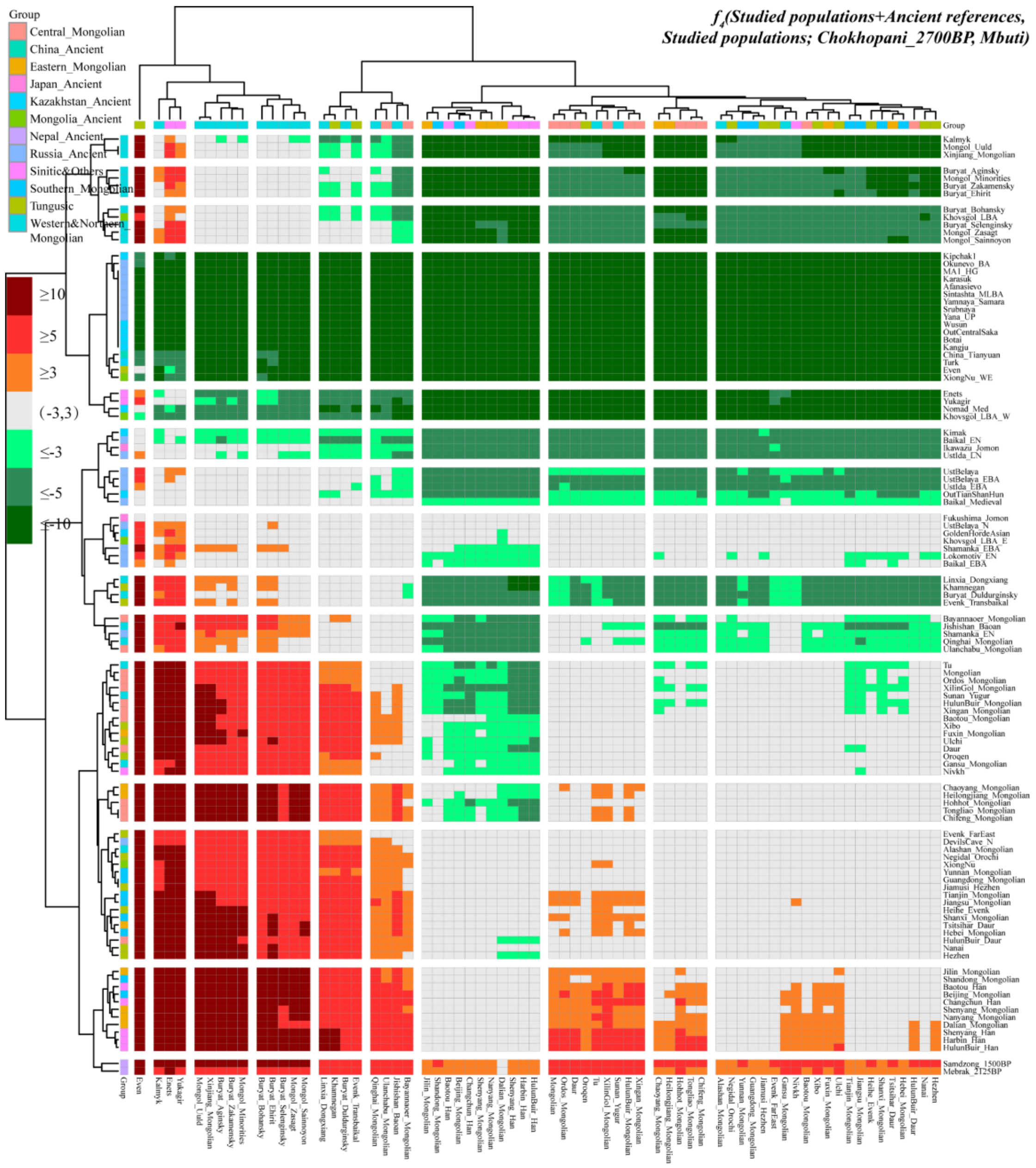

**Supplementary Fig. 37** The genetic contribution of ancient populations on the Tibetan Plateau to target populations revealed by *$f_4$-statistics* in the form of ***$f_4$(Studied populations + Ancient references, Studied populations; Chokhopani_2700BP, Mbuti)***, Chokhopani_2700BP is used as the reference baseline.

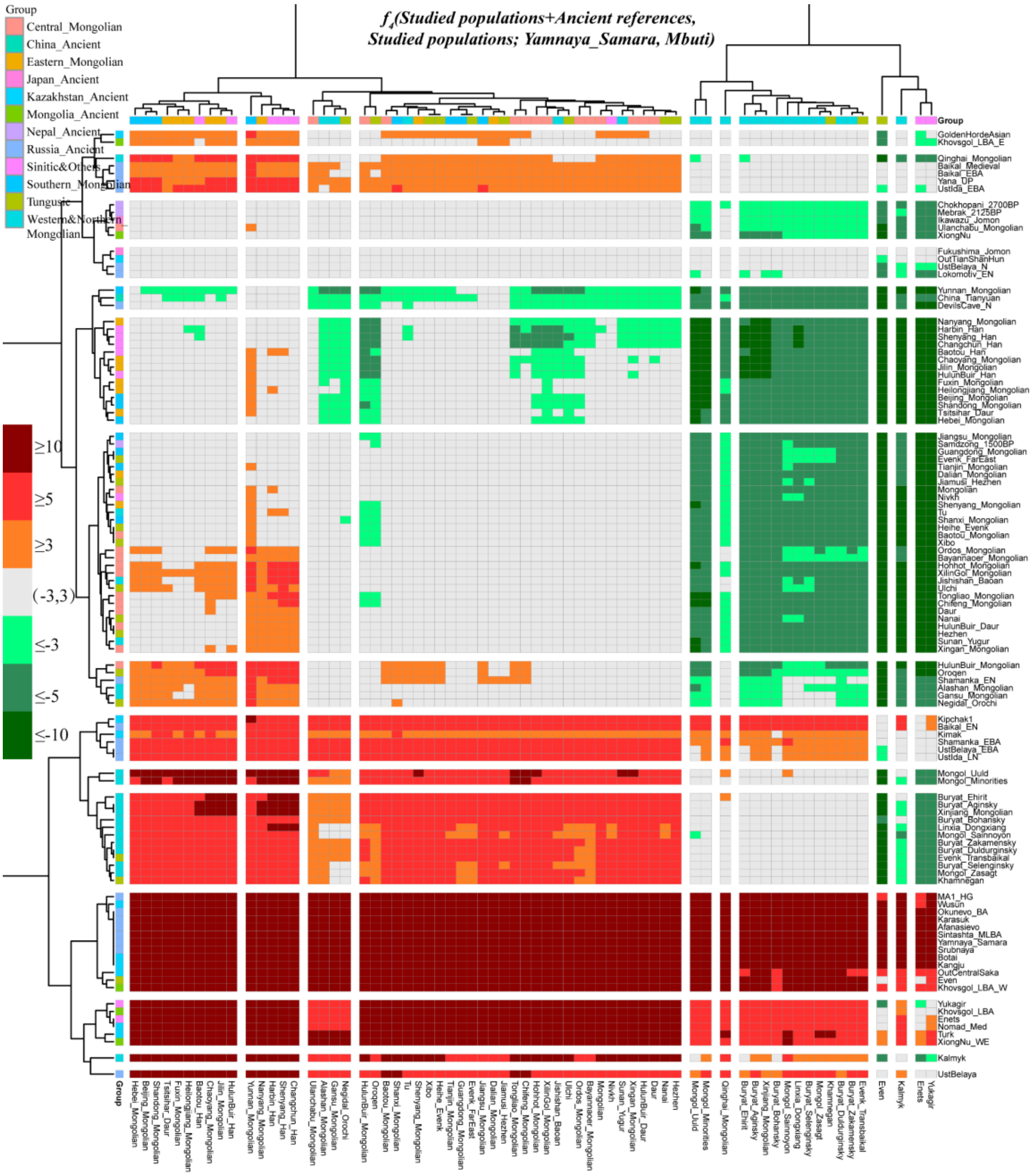


**Supplementary Fig. 38** The genetic contribution of western Eurasian populations to studied populations revealed by *$f_4$-statistics* in the form of ***$f_4$(Studied populations + Ancient references, Studied populations; Yamnaya_Samara, Mbuti)***, Yamnaya_Samara is used as the reference baseline.

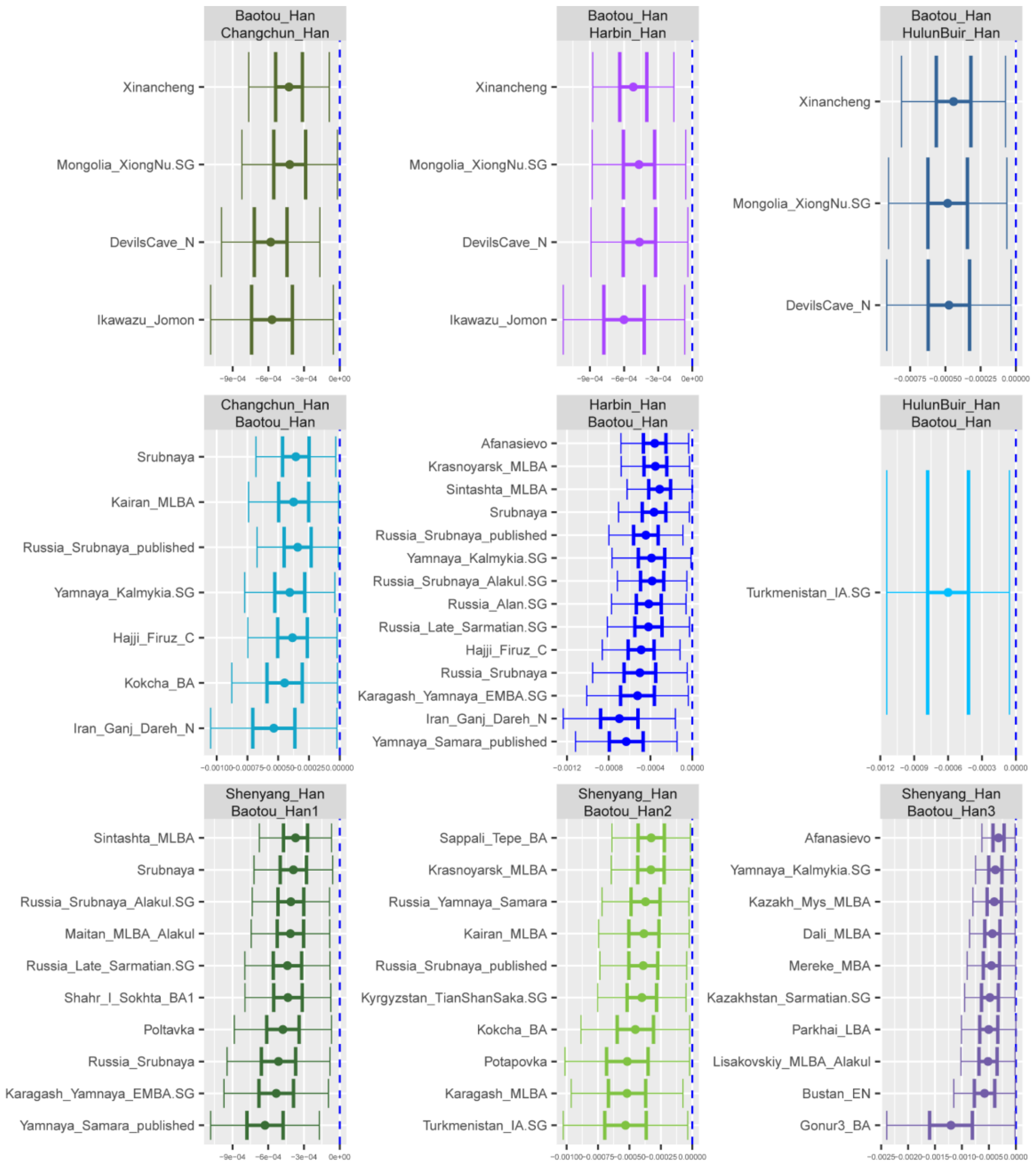


**Supplementary Fig. 39** The values of ***$f_4$(Studied Han Chinese1, Studied Han Chinese2; Ancient Eurasian populations, Yoruba)***.
